# Evolutionary genomics of local adaptation, climate vulnerability, and linarin metabolism in *Opisthopappus longilobus*

**DOI:** 10.64898/2026.01.20.700049

**Authors:** Ruyue Jing, Jiangshuo Su, Aiping Song, Wei Chen, Ming Wang, Jiayu Xue, Yaru Zhang, Xiaojin Su, Yanru Wang, Zhongyu Yu, Jun He, Liangju Ma, Fei Zhang, Haibin Wang, Zhenxing Wang, Sumei Chen, Weimin Fang, Jiafu Jiang, Fadi Chen

## Abstract

Global climate change and human activities are posing substantial threats to biodiversity. The genus *Opisthopappus* (*O. taihangensis* and *O. longilobus*), endemic to China’s Taihang Mountains, possesses great ecological, ornamental and medicinal value. However, it is confronted with the pressures of habitat fragmentation and climate change. Here, we present the first haplotype-resolved, chromosome-scale genome assembly of *O. longilobus* (∼2.95 Gb) and re-sequence 115 individuals across its range. Comparative analyses show *Opisthopappus* is sister to *Artemisia*-*Chrysanthemum*, with *Opisthopappus* and *Chrysanthemum* diverging at 5.15-5.18 million years ago. A profound genetic divergence is evident between *O. taihangensis* and *O. longilobus*, resulting in two distinct lineages within each species, driven by geography and climate. Our analyses indicate restricted gene flow, low diversity, and recurrent demographic bottlenecks collectively contribute to their endangerment. By integrating population genomics and environmental variables, we identified 4,437 core adaptive genes linked to water deprivation, and metabolism. Genomic offset predicts higher maladaptation risk in populations under drastic climate change. Furthermore, diverged promoters of two O-methyltransferase genes, *OMT250* and *OMT310*, likely underlie the differential acacetin/linarin accumulation between *C. morifolium* and *O. longilobus*. These findings advance understanding of *Opisthopappus*’ evolution and climate vulnerability, offering a model for genomics-guided biodiversity conservation.

## Introduction

Global climate change is driving the profound transformation of natural ecosystems. By accelerating habitat loss and fragmentation, it substantially increases the risk of local extinction and has become a major threat to the persistence of biodiversity (Thuiller et al., 2005; Harrison, 2020; Urban, 2024). The vulnerability of species to climate change is determined by their response strategies, which primarily involve two mechanisms: shifting their geographic ranges to track suitable climates or adapting in situ through phenotypic plasticity and genetic evolution (Aguirre-Liguori et al., 2021; Shen et al., 2024). However, for long-lived sessile organisms such as perennial plants, migration is virtually unfeasible in the face of rapid climate change, inevitably resulting in widespread and persistent maladaptation throughout their lifespans (Kooyers et al., 2025). Thus, unraveling how species genetically responded to past geological and climatic shifts is essential to interpret current adaptations and assess future evolutionary potential under climate change (Helmstetter et al., 2020). Encouragingly, the advancing affordability of genomic technologies now enables efficient production of population genomic datasets. These data are vital for identifying adaptive genetic variants associated with environmental gradients, which can be integrated into predictive models to better assess climate change vulnerability and proactively inform targeted conservation strategies (Hoffmann et al., 2021; Chen et al., 2022b; Hou et al., 2024a; Zhang et al., 2024).

The Taihang Mountains serve as a major phylogeographic boundary in northern China, where steep environmental gradients and rugged topography restrict gene flow and facilitate local adaptation (Wang & Li, 2008; Liu et al. 2018; Ye et al., 2021). The uplift of the Taihang Mountains, occurring rapidly after the Late Pleistocene, created a landscape that now provide a historical refuge for many species during climatic oscillations (Wang &Yan, 2014). *Opisthopappus* C. Shih (Asteraceae) is a narrowly endemic cliff-dwelling plant in the Taihang mountains, offering a natural system to investigate ecological evolution, species divergence, and potential decline in extreme rocky habitats. The genus comprises two sister species, *O. taihangensis* and *O. longilobus*, which inhabit cliffs and rock crevices and tolerate chronic stresses such as drought, low temperatures, and nutrient-poor substrates. Despite their shared habitat constraints, the two species exhibit significant niche differentiation (Wang & Yan, 2014; Yang et al., 2020; Chen et al., 2022a). Given its high habitat specialization, limited regeneration capacity, and exposure to dramatic climate change and destructive human activities, *Opisthopappus* faces a significantly elevated risk of population decline and genetic erosion (Li et al., 2009b; Ye et al., 2021; Liu et al., 2023). Consequently, it is recognized as a nationally prioritized conservation target among threatened plant species in China. However, its evolutionary history and adaptive genomics remain unresolved, and an integrative framework connecting population structure, local adaptation, and range-wide risk is lacking.

Flavonoids, a prominent class of polyphenolic secondary metabolites, play essential roles in plant adaptation by mediating stress responses, signaling, and interactions with microbial communities. A deeper understanding of their biosynthetic is therefore key to unraveling plant evolution and ecological adaptation (Pollastri and Tattini, 2011; Agati et al., 2012; Falcone Ferreyra et al., 2012; Šamec et al., 2021; Daryanavard et al., 2023; Agati et al., 2025). The biosynthesis of flavonoids is highly influenced by environmental factors, with various modifications such as methylation and glycosylation critically altering their solubility and bioavailability (Saito et al., 2013; Berim and Gang, 2016; Alseekh et al., 2020). Linarin (acacetin-7-O-rutinoside) is a characteristic flavonoid derivative of the Asteraceae family, listed in the Chinese Pharmacopoeia (Committee, 2025) for its traditional medicinal applications (Mottaghipisheh et al., 2021). Modern pharmacological studies confirm its multi-target efficacy against Alzheimer’s disease, cancer, inflammation, and diabetes, as well as its potential as a promising candidate in COVID-19 research (Lou et al., 2011; Ding et al., 2025; Liu et al., 2025; Wang et al., 2025a). Previous studies have proposed key steps in linarin biosynthesis of *C. indicum* (Jiang et al., 2019; Wu et al., 2022; Deng et al., 2024; Hou et al., 2024b). The elevated linarin content in *Opisthopappus* species relative to most wild chrysanthemum plants establishes their medicinal and economic importance (Wei et al., 2023; Fan et al., 2025). However, the complete biosynthetic pathway of linarin remains genetically unresolved in *O. longilobus*, and the connection between its linarin content and local adaptation presents an open question.

In this study, we assembled a haplotype-resolved genome of *O. longilobus* and resequenced 115 representative *Opisthopappus* individuals across its distribution range. Based on these genomic datasets, our research aims to: (1) reveal the phylogenetic context and evolutionary position of *O. longilobus* among angiosperms, particularly within the Asteraceae family; (2) characterize spatial patterns of genetic diversity, population structure, and demographic history, uncover the mechanisms underpin environmental adaptation and assess genetic vulnerability under future climate change; and (3) pinpoint critical biosynthetic pathway steps of linarin. These findings offer novel insights into genome evolution, local adaptation, and specialized metabolite biosynthesis in this endangered cliff-dwelling species, thereby establishing a strategic framework for its future conservation and molecular breeding.

## Results

### Haplotype-resolved genome assembly of *O. longilobus*

Cytological and genomic analyses indicated that the sequenced *O. longilobus* plant is a diploid, with the estimated genome size ranging from 2.92 to 3.36 Gb (2n = 2x = 18; Supplementary Figures 1-3). The genome survey revealed a high heterozygosity rate of 1.91% and high repeat content of 77.26% (Supplementary Figure 3A, B), which presents a challenge for accurate genome assembly. The HiFi reads assembly yielded a draft genome of 5.92 Gb (contig N50 of 169.20 Mb; Supplementary Tables 1, 2), which was scaffolded using 408 Gb Hi-C data (∼136×). The 3D-DNA pipeline successfully anchored 99.67% of the sequences into 18 pseudochromosomes (Figure 1A; Supplementary Tables 1 and 3), corroborating the karyotype evidence. The Hi-C interaction heatmap clearly demonstrated that the 18 pseudochromosomes could be organized into nine homoeologous clusters (Figure 1B; Supplementary Figure 4). Haplotype analysis revealed two haplotypes: Hap1 (2.947 Gb, scaffold N50 of 336.00 Mb) and Hap2 (2.951 Gb, scaffold N50 of 329.20 Mb) (Figure 1A; Table 1; Supplementary Table 4, 5), both representing the majority of the assembled genome size. A high degree of collinearity was observed between Hap1 and Hap2, aside from a minor inversion on chromosome 9 (Supplementary Figures 5, 6A, B, and 12D).

**Figure 1.**
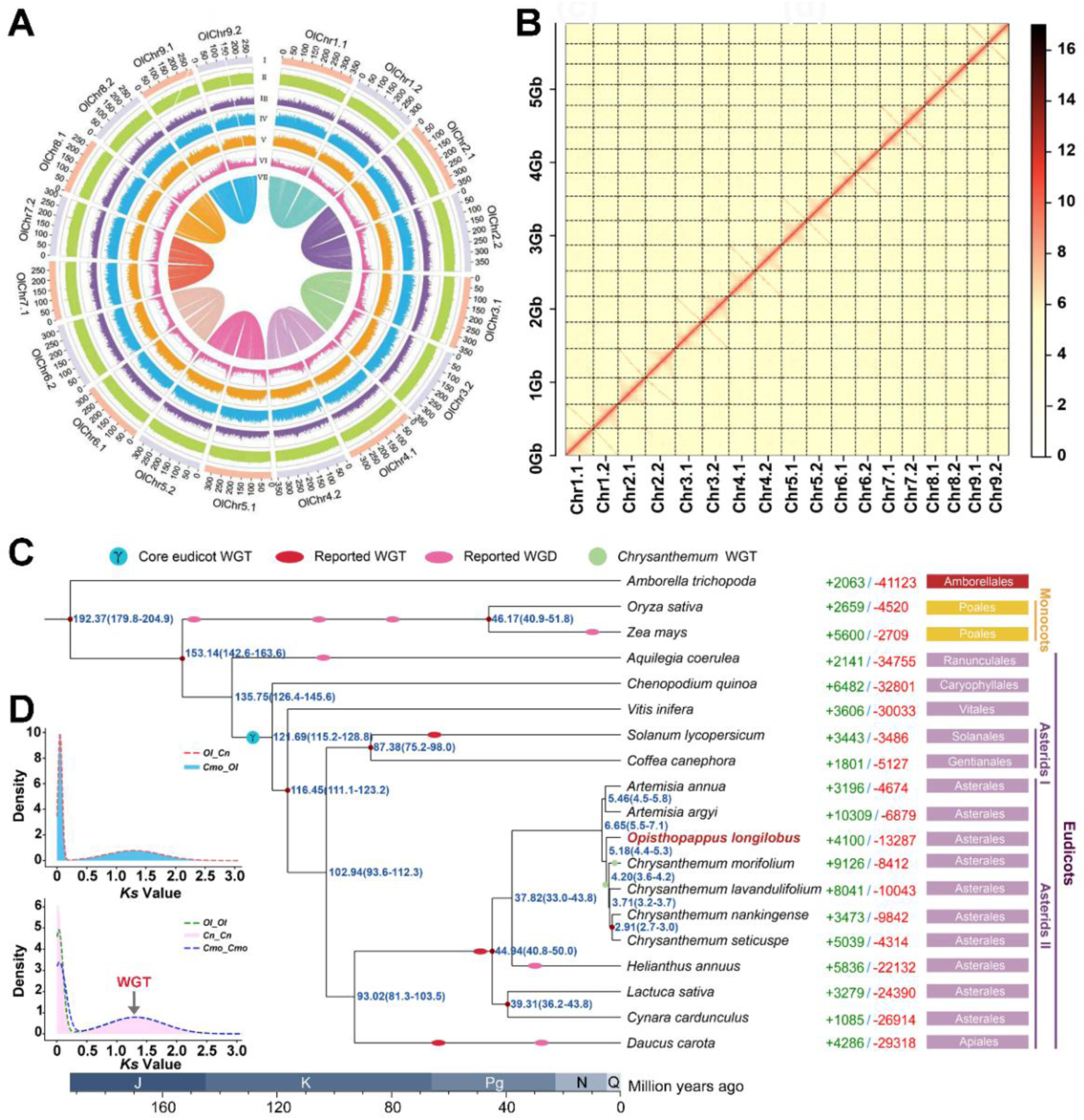
Genome assembly, evolutionary relationship, and whole-genome duplication history of *O. longilobus*. **A,** Circos plot displaying genomic features of *O. longilobus*. I. Pseudo-chromosome length (Mb); II. GC content; III. *Gypsy* transposon proportion; IV. *Copia* transposon proportion; V. LTR transposon proportion; VI. Gene density; VII. Collinear regions within the haplotype assembly. The two sets of haplotype pseudochromosomes are distinguished by the suffixes “.1” and “.2”. **B,** Hi-C contact map of *O. longilobus* genome. **C,** The phylogenetic tree illustrates the evolutionary relationship, divergence times, and gene family dynamics of *O. longilobus* and 18 other flowering plants. Gene family expansions (green) and contractions (red) are shown on the right. Fossil calibration points (red circles) are sourced from TIMETREE (http://www.timetree.org/). Divergence times, estimated via Maximum Likelihood (PAML), are labeled in blue with 95% confidence intervals in parentheses. J, Jurassic; K, Cretaceous; Pg, Palaeogene; N, Neogene; Q, Quaternary. **D,** *Ks* distribution within *O. longilobus* (Ol), *C. nankingense* (Cn), and *C. morifolium* (Cmo).

**Table 1.** Statistics for genome assembly and annotation of *O. longilobus*.

| **Taxa** | **Haplotype 1** | **Haplotype 2** |
| --- | --- | --- |
| **Genome assembly** |  |  |
| Assembly estimated genome size (Gb) | 2.92 | - |
| Assembled genome size (Gb) | 2.95 | 2.95 |
| GC content (%) | 36.44 | 36.43 |
| Contig N50 (Mb) | 169.20 | - |
| Scaffold N50 (Mb) | 336.00 | 329.20 |
| Total length of contig (Gb) | 3.21 | 3.21 |
| Total length of scaffold (Gb) | 2.95 | 2.95 |
| Complete BUSCOs (%) | 98.57 | 98.57 |
| Longest scaffold (Mb) | 369.28 | 398.10 |
| LTR Assembly Index | 20.79 | 24.65 |
| Anchor rate (%) | 99.67 | 99.67 |
| **Genome annotation** |  |  |
| Repetitive sequence (%) | 64.28 | 64.56 |
| Number of protein-coding genes | 58,394 | 57,850 |
| Complete BUSCOs (%) | 97.80 | 97.90 |

The HiFi reads showed a 100% mapping rate, and the DNBSEQ transcriptomic reads averaged 86.72%, with both datasets exhibiting uniform chromosomal coverage (Supplementary Table 6; Supplementary Figure 7A, B). Mercury evaluation produced a consensus quality value of 42.5, corresponding to a low error rate of 5.6e-5. Additionally, approximately 64% of *K*-mers were recovered per haplotype, with the combined assembly reaching 96.79% (Supplementary Figure 6C; Supplementary Table 7). The BUSCO analysis revealed a combined completeness of 99.10%, with each haplotype (Hap1 and Hap2) reaching 98.57% (Supplementary Table 8). These results demonstrate a high degree of continuity and completeness of the *O. longilobus* genome. Moreover, telomeric motifs (TTTAGGG)_n_ were identified at both ends of 17 pseudochromosomes (94.4% completeness), validating the haplotype-phased assembly (Supplementary Figures 2H and 7C, D; Supplementary Table 9). Notably, no telomeric signal was detected on one end of chromosome 5.1 (Chr5.1), suggesting potential structural truncation or rearrangement (Supplementary Figure 7C). Four of the 18 pseudochromosomes exhibited detectable size heterogeneity in their centromeric regions, reflecting the variability in centromere structure across the chromosomes (Supplementary Figure 7C, D; Supplementary Table 10). The data indicate that the *O. longilobus* genome assembly achieves near Telomere-to-Telomere (T2T) standards (Supplementary Table 11). Overall, this high-quality, haplotype-resolved assembly establishes *O. longilobus* as a key genomic reference, enabling precise comparative genomics and adaptive evolution studies.

### Genome annotation of *O. longilobus*

Transposable elements (TEs) comprised 64.28% (Hap1) and 64.56% (Hap2) of the genome, predominantly consisting of long terminal repeat (LTR) retrotransposons which accounted for 57.98% and 58.12% of each haplotype, respectively. Notably, *Copia* elements were more abundant (33.44% in Hap1, 37.56% in Hap2) than *Gypsy* elements (21.80% in Hap1, 24.20% in Hap2) (Supplementary Figure 12A; Supplementary Table 13). The LTR Assembly Index (LAI) scores for Hap1 and Hap2 were 20.79 and 24.65, respectively, both exceeding the gold-reference-quality threshold (Table 1; Supplementary Table 14). Noncoding RNAs (ncRNAs) perform a wide range of essential regulatory functions at both transcriptional and post–transcriptional levels. Therefore, we annotated and quantified microRNAs, transfer RNAs, ribosomal RNAs, small nuclear RNAs in the *O. longilobus* genome (Supplementary Table 15). Given that 5S rRNA is a reliable cytogenetic marker for species ploidy levels (He et al., 2022), we further validated its chromosomal localization through FISH analysis. Two distinct hybridization signals were observed on the chromosomes (Supplementary Figure 2G), consistent with the characteristics of a diploid genome. The presence of these rRNA sequences reinforces the completeness and reliability of the assembled *O. longilobus* genome.

Protein-coding genes were predicted using an integrated methodology that combines *de novo* prediction, transcriptome evidence, and homology-based annotation (Supplementary Table 16). In total, 58,394 and 57,850 protein-coding genes were identified in Hap1 and Hap2 of the *O. longilobus* genome, respectively (Table 1; Supplementary Table 17). A total of 63,223 duplicated genes were categorized into five distinct types: 4,597 whole-genome duplicates (WGD, 7.27%), 43,074 dispersed duplicates (DSD, 68.13%), 7,194 transposed duplicates (TRD, 11.38%), 3,957 proximal duplicates (PD, 6.26%), and 4,401 tandem duplicates (TD, 6.96%) (Supplementary Tables 18 and 19). Overall, 55,063 (94.30%) and 54,567 (94.32%) protein-coding genes in Hap1 and Hap2 were successfully annotated in at least one public database (Supplementary Table 20). BUSCO assessment revealed that 97.80% and 97.90% of core conserved genes were complete in Hap1 and Hap2, respectively, demonstrating the high quality of gene prediction and annotation (Table 1; Supplementary Table 8). Due to its superior in continuity and completeness, Hap1 was selected for subsequent genome evolutionary and population genetics analysis.

### Comparative genomics and evolutionary analysis

Phylogenetic analysis integrating the *O. longilobus* genome with 18 representative angiosperms and 22 Asteraceae genomes helps to elucidate its evolutionary trajectory and critical node in the radiation of the Asteraceae family (Supplementary Figure 8; Supplementary Tables 22-24). Gene family evolution analysis revealed that 4,100/4,047 families underwent expansion, while 13,287/19,276 gene families contracted within the *O. longilobus* genome (Figure 1C, 2A; Supplementary Table 26). GO enrichment analysis of the rapidly expanded gene families in *O. longilobus* compared to angiosperms revealed significant associations with functions such as “DNA binding”, “DNA integration”, “nucleic acid binding”, “telomere maintenance”, and “DNA helicase activity” (Supplementary Table 25). Significant enrichment was observed in KEGG pathways related to metabolic and phenylpropanoid biosynthesis (Supplementary Table 25). Additionally, the functional characterization of these expanded families in *O. longilobus* compared to other Asteraceae species showed significant enrichment in flavonoid-related metabolic pathways, with a notable emphasis on O-methyltransferase activity (Figure 2B; Supplementary Table 27).

**Figure 2.**
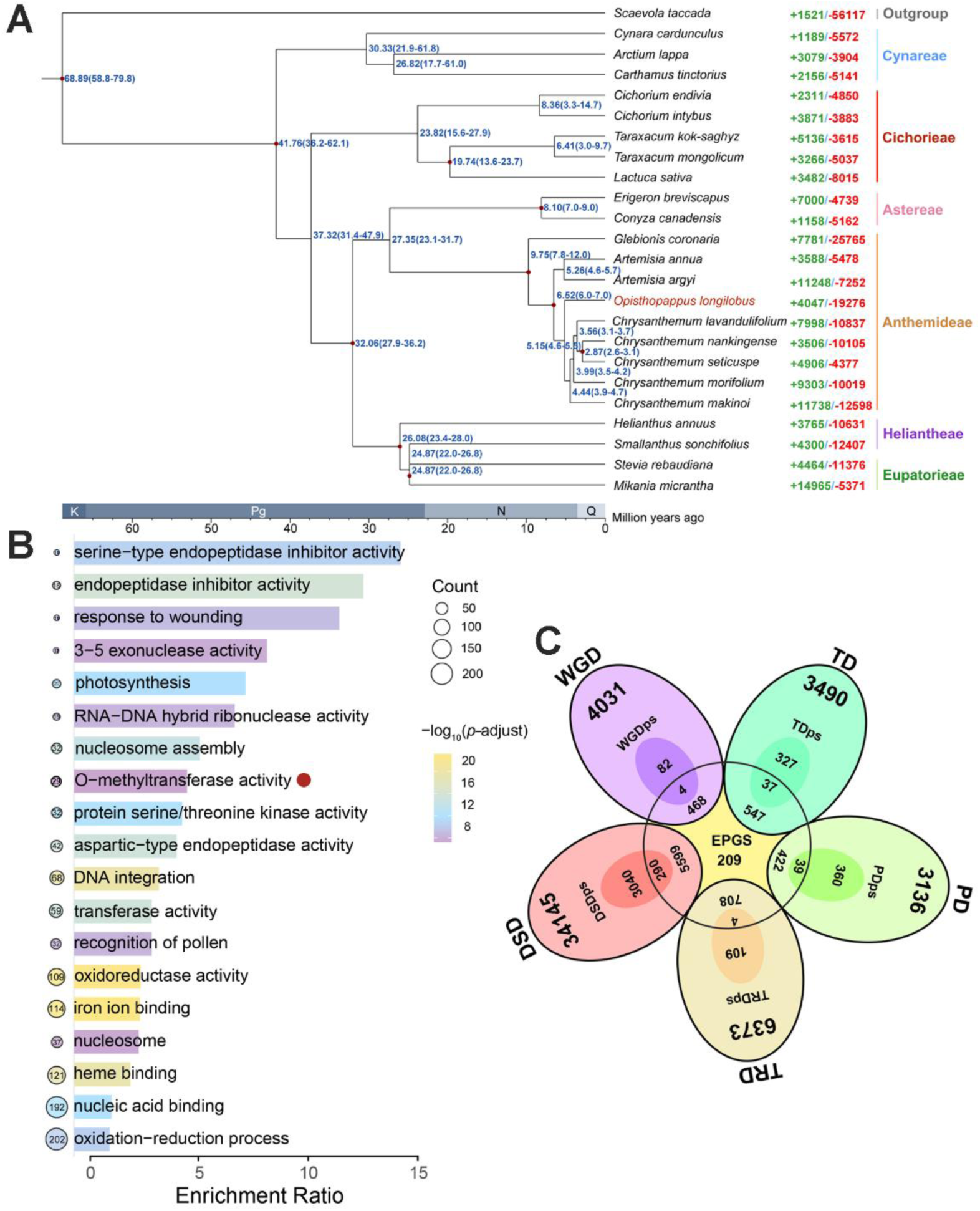
Genome evolution and gene duplication of *O. longilobus*. **A,** The phylogenetic tree illustrates the evolutionary relationship, divergence times, and gene family dynamics of *O. longilobus* and 22 other Asteraceae plants. Gene family expansions (green) and contractions (red) are shown on the right. Divergence times (blue) with 95% confidence intervals were estimated using PAML and calibrated with TIMETREE fossil data (http://www.timetree.org/). K, Cretaceous; Pg, Palaeogene; N, Neogene; Q, Quaternary. **B,** GO enrichment analysis on the expanded gene families (EPGs). **C,** Venn diagram shows the number of shared and specific gene duplications between the EPGs and five duplication types, i.e., dispersed duplications (DSD), proximal duplications (PD), tandem duplications (TD), transposed duplications (TRD), and whole-genome duplications (WGD). Duplicate genes under positive selection were assigned the suffix “ps”.

Phylogenetic analyses confirmed *O. longilobus* as the sister to the *Artemisia*-*Chrysanthemum* clade. The topology distinctly separated *Opisthopappus* from these closely related genera, with strong bootstrap support for its monophyly, thereby reaffirming its recognized generic classification (Figures 1C, 2A; Supplementary Figure 9). The divergence time between the *Opisthopappus* and *Chrysanthemum* genera was estimated at approximately 5.15-5.18 million years ago (Mya), while the divergence between *O. longilobus* and *Artemisia* species was estimated at around 6.52-6.65 Mya (Figures 1C, 2A; Supplementary Table 28). The chloroplast genome-based Asteraceae phylogenetic tree also supported the relationships inferred from nuclear genomes (Supplementary Figure 10).

Whole-genome duplication/triplication (WGD/WGT) and the proliferation of repetitive elements act as fundamental mechanisms driving plant genome evolution (Vanneste et al., 2014). Dot plot analyses based on WGDI and JCVI indicated the presence of extensive triplicated collinear blocks, with synonymous substitution rate (*Ks*) values peaking around 1.2 (Figure 1D; Supplementary Figure 11A-C), suggesting *O. longilobus* also experienced the Asteraceae-specific WGT-1 event with *Chrysanthemum* species (Song et al., 2023). Additionally, the *Ks* peak at approximately 0.1 (Figure 1D) is likely reflects recent tandem (TD) and proximal (PD) gene duplications (Supplementary Figure 11F). A recent LTR burst was also evident in the *O. longilobus* genome, with highly consistent insertion-time distributions between Hap1 and Hap2 and similar patterns for *Copia* and *Gypsy* insertions (∼0.11 Mya). In addition, comparable lineage-specific recent bursts were observed across Asteraceae (Supplementary Figure 12B-D).

Gene duplication is widely acknowledged as a significant driver of evolutionary innovation (Qiao et al., 2019). Analysis of *Ka*/*Ks* ratios showed that TD and PD gene pairs had higher values than other duplication types (Supplementary Figure 11E), indicating rapid sequence divergence and strong positive selection among these duplications. Furthermore, a comparison of duplication types in expanded gene families (EPGs) and 262 genes under positive selection revealed that TRD and DSD duplicates comprised a large proportion of both categories in *O. longilobus* (Figure 2C; Supplementary Tables 26, 29). GO enrichment analyses indicated functional divergence among different duplication modes, with TDs and TRDs showing significant enrichment in the term “response to stimulus” (Supplementary Figure 13), implying their selective retention for adaptive evolution. Water stress is a significant environmental challenge for *Opisthopappus* (Yang et al., 2020). By integrating transcriptomic data with evolutionary insights from EPGs, positively selected genes (PSGs), and positively selected duplicated genes (DupPSGs), we identified a set of core drought-responsive genes and formulated an initial mechanistic framework of environmental adaptation for *O. longilobus* (Supplementary Figure 14; Supplementary Table 30).

### Genomic landscapes of population structure, diversity, and differential

We conducted whole-genome resequencing (35.16×) for 66 *O. taihangensis* accessions and 49 *O. longilobus* accessions, spanning the entirety of this genus’s distribution in the Taihang Mountain region of China (Figure 3A, Supplementary Figure 15, and Supplementary Tables 31-33). Phylogenetic relationships were inferred using a subset of 3,808,756 LD-pruned SNPs. ADMIXTURE analysis indicated that at K = 2, individuals of *O. taihangensis* (OT) and *O. longilobus* (OL) were distinctly assigned to two separate clusters. When K was increased to 3, twelve *O. longilobus* individuals collected from Shanxi Province (OL_SX) were differentiated from those sampled in Hebei Province (OL_HB), diverging into two distinct lineages. Cross-validation error analysis identified the optimal number of clusters as four, resulting in the division of *O. taihangensis* into Northern and Southern lineages (designated as OT_N and OT_S, respectively) (Figure 3b; Supplementary Figure 16). The Maximum-likelihood (ML) trees reconstructed with or without *A. annua* as the outgroup both supported the four geographic clades, with the former suggesting *O. longilobus* as the potential ancestral species (Figure 3B and Supplementary Figure 17A). PCA further corroborated these results, with PC1 (52.56%) effectively distinguishing between OT and OL, while PC2 (14.46%) and PC3 (11.20%) further separated the two genetic lineages within OT and OL, respectively (Figure 3C and Supplementary Figure 17B, C). To comprehend the phylogenetic relationships and geographic distribution of *Opisthopappus* species, we performed kinship and isolation by distance (IBD) analyses. The results revealed that individuals collected from proximate locations exhibited higher kinship coefficients (Supplementary Figure 17D), coupled with a significant positive correlation between geographic and genetic distances (Mantel’s *r* = 0.75, *P* < 0.001) (Supplementary Figure 32C).

**Figure 3.**
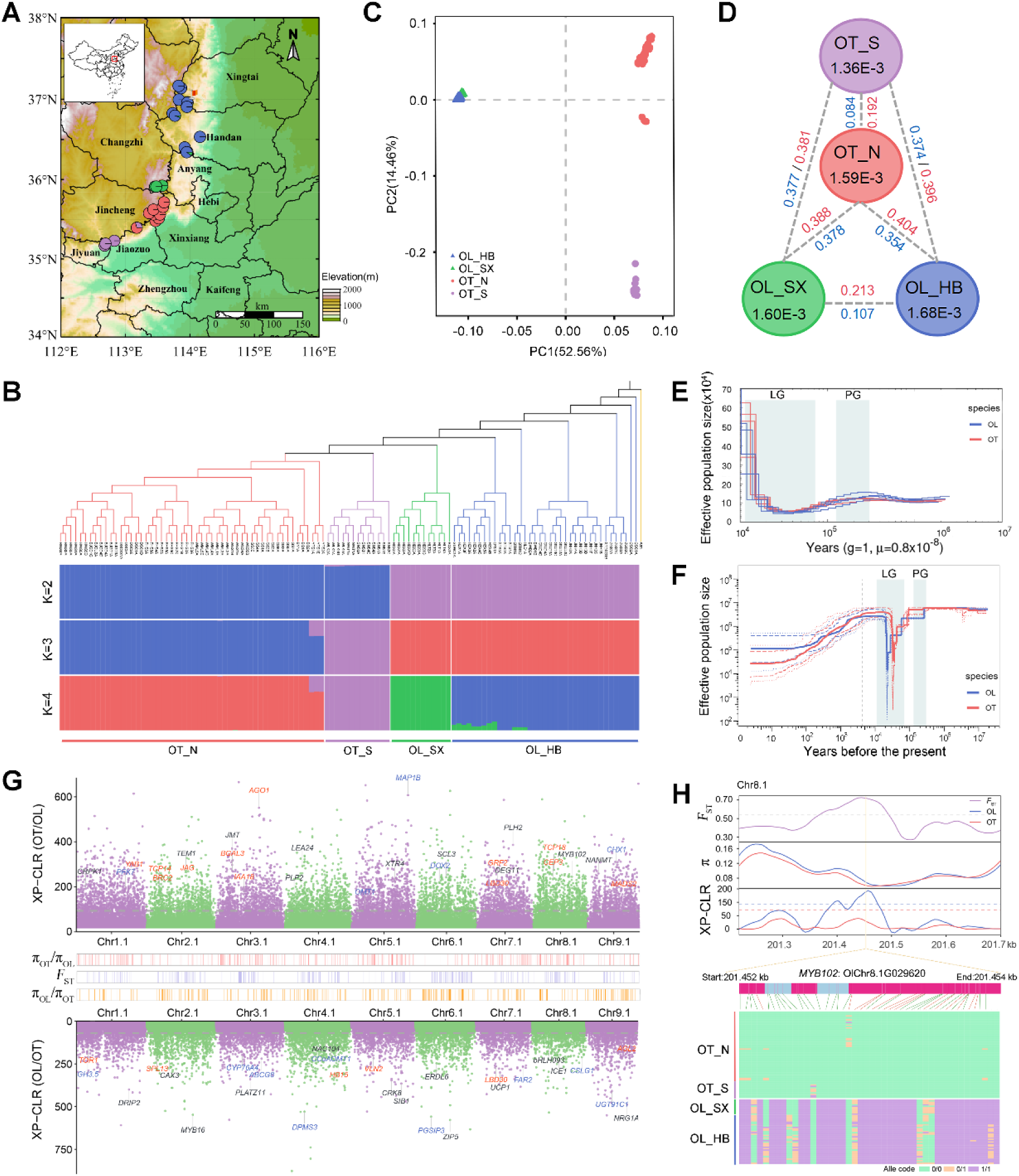
Population structure, demographic history, and genetic divergence analyses of *Opisthopappus* species. **A,** Geographic distribution of 12 sampled *O. longilobus* populations and 16 sampled *O. taihangensis* populations, where colors of the pie charts represent ancestral components inferred by ADMIXTURE at K = 4. The location of the individual used for genome assembly is indicated by a red flag. **B,** Upper panel: The ML phylogeny of the 115 *Opisthopappus* accessions with *Artemisiae annua* (Ann) as the outgroup. Branch colors indicate different groups based on the population structure. Lower panel: Population structure of *Opisthopappus* with K from 2 to 4. **C,** PCA plot of the first two eigenvectors, with different colors correspond to the four subgroups. **D,** Genetic diversity and population differentiation among four subgroups. The value in each circle represents the level of nucleotide diversity (*π*) for each group. The value in black on each line indicates pairwise fixation index (*F*ST), while the value in red indicates pairwise nucleotide divergence (*D*XY). **E,** Demographic history of *O. longilobus* and *O. taihangensis* inferred by PSMC. The bounds of the 95% and 75% confidence intervals for the estimated effective population size are indicated by dashed and dotted lines, respectively. **F,** Demographic history of *O. longilobus* and *O. taihangensis* inferred by Stairway Plot v2. In (**E** and **F)**, the last glacial (LG, 11-70 kya) and penultimate glaciation (PG, 130-300 kya) are shaded in light-green. The beginning of Meghalaya is highlighted by a gray dashed vertical line (4.2 kya). **G,** Genome-wide distribution of selective sweeps identified for *O. longilobus* and *O. taihangensis*. Candidate genes annotated by red, blue, and black characters are associated with development, metabolism, and stress, respectively. **H,** Selection signatures and genotype heatmap (bottom) of *MYB102*. 14 nonsynonymous variants located in the genic region of *MYB102* are highlighted by red vertical lines.

We characterized the genetic parameters across species, lineages, and populations (Supplementary Table 34). Compared to *O. taihangensis*, *O. longilobus* exhibited greater nucleotide diversity (*π*), heterozygosity, and Tajima’s *D*, along with lower inbreeding coefficients (*F*_IS_ and *F*_ROH_) (Figure 3D and Supplementary Figure 18). The genetic diversity within the four intrageneric lineages (*π* = 1.36e-3∼1.68e-3) was found to be lower than that of their respective species (*π* = 1.74e-3 for OL and 1.62e-3 for OT), with the OT_S lineage showing the lowest nucleotide diversity, heterozygosity, and Tajima’s *D*, and the highest inbreeding (Supplementary Figure 19). Considerable genetic variation was also observed among different populations (Supplementary Figure 20). Notably, the observed positive Tajima’s *D* values indicated potential demographic bottlenecks and balancing selection within the *Opisthopappus* species. It is important to highlight that the nucleotide diversity of the two *Opisthopappus* species was lower than that of two widespread Asteraceae species and even that reported for endangered plant species (Supplementary Figure 21), underscoring the urgent need for conservation efforts for *Opisthopappus* plants.

A high pairwise fixation index (*F*_ST_ = 0.335) and nucleotide divergence (*D*_XY_ = 0.398) were observed between *O. longilobus* and *O. taihangensis*, indicating significant genomic divergence (Supplementary Figure 22A). The *F*_ST_ values (0.084-0.107) and *D*_XY_ values (0.192-0.213) of intra-specific lineages were obviously lower than those of inter-specific lineages (*F*_ST_ = 0.354-0.378 and *D*_XY_ = 0.381-0.404) (Figure 3D). Similar patterns of genetic differentiation were evident across populations (Supplementary Figure 23 and Supplementary Table 35). Additionally, differing patterns of linkage disequilibrium (LD) decay revealed distinctions between species and lineages (Supplementary Figure 17D). These findings were further supported by detecting stronger signals of gene flow and introgression within specific lineages compared to those between species (Supplementary Figures 24 and 25; Supplementary Tables 36 and 37). Given the relatively weak divergence of intra-specific lineages, we compared the demographic histories of the two *Opisthopappus* species. Both the PSMC (Figure 3E) and Stairway plot (Figure 3F) approaches indicated that the two species maintained stable and similar effective population sizes (*Ne*) until the onset of the penultimate glaciation (PG, 130-300 kya), aligning with the timing of divergence for *O. longilobus* and *O. taihangensis* as inferred by SMC++ (Supplementary Figure 22B). Thereafter, *O. longilobus* experienced a drastic decline from PG to the middle last glacial (LG), reaching its lowest level before undergoing a population expansion around the last glacial maximum (LGM, 19-26.5 kya). In contrast, *O. taihangensis* showed a delayed decline and an earlier recovery. The recent demographic trajectory reconstructed via Stairway plot revealed a continuous population decline since the Meghalaya period (4.2 kya), with *O. taihangensis* undergoing a more dramatic drop and maintaining a smaller *Ne* (Figure 3F).

### Genome-wide scans for selective sweep between *O. taihangensis* and *O. longilobus*

To clarify the genomic regions underlying intergenic differences, we excluded four admixed individuals with Q < 0.85 from the selective sweep analysis. A total of 793 potential sweep regions (PSRs) were identified in *O. longilobus* (OT/OL), which spanned 94.26 Mb and encompassed 1,672 genes. In comparison, we found 961 highly divergent genomic regions for *O. taihangensis* (OL/OT), covering 116.5 Mb (3.95%) of the reference genome and including 1,961 genes (Figure 3G and Supplementary Table 38). The PSRs in *O. longilobus* were evenly distributed across nine pseudochromosomes, ranging from 2.06% in Chr9.1 to 4.47% in Chr8.1. In contrast, PSRs of *O. taihangensis* on Chr6.1, Chr2.1, and Chr4.1 accounted for 53.12% of the total selective regions, reflecting a disproportionate chromosomal distribution (Supplementary Figure 26A). Notably, the selective sweep signals within both comparative groups were predominantly found in intergenic regions, especially in promoter areas and regions 10-500 kb away from gene bodies (Supplementary Figure 26B, C), suggesting that natural selection may have preferentially targeted non-genic regulatory elements during the evolution of *Opisthopappus*.

GO analysis revealed that the selective genes in *O. longilobus* were predominantly enriched in biological processes such as “multicellular organism development”, “jasmonic acid metabolic process”, “root system development”, “response to light stimulus”, “vegetative to reproductive phase transition of meristem”, and “response to oxidative stress”. In contrast, the selective genes in *O. taihangensis* displayed significant GO terms including “mono carboxylic acid metabolic process”, “positive regulation of metabolic process”, “cell cycle process”, “regulation of gene expression”, and “fatty acid biosynthetic process” (Supplementary Figure 27; Supplementary Table 39). These findings suggest a pattern of independent adaptive evolution between the two species. Remarkably, several functionally characterized genes involved in development, stress responses, and metabolism were identified within PSRs (Figure 3G). For instance, *MYB102* (OlChr8.1G029620; Chr8.1: 201.40-201.56 Mb) showed distinct selection signals in *O. longilobus* (Figure 3H) and was found to be an expanded gene relative to other Asteraceae species (Supplementary Table. 26). There were 44 SNPs located within the genic region of *MYB102*, with *O. longilobus* and *O. taihangensis* individuals carrying distinctly different haplotypes (Figure 3H). Expression analysis indicated that *MYB102* was induced under both waterlogging and drought stress conditions in *O. longilobus* (Supplementary Tables 30 and 40). Orthologs of this gene in Arabidopsis and rice are involved in diverse biotic and abiotic stress responses (Zhu et al., 2018; Piao et al., 2019). These results suggest a crucial role for *MYB102* in the adaptive evolution of *O. longilobus*, potentially contributing to species diversification. Additionally, several stress-responsive genes were highlighted, including *CRPK1*, *JMJ*, *PLP2*, *bHLH93*, *NAC104*, and *ICE1* (Figure 3G).

Leaf morphology, as a key functional trait, reflects adaptive responses shaped by environmental pressures (Fritz et al., 2018). The most prominent difference between the two species lies in their leaf shape (Supplementary Figure 28A). As expected, several organ development genes with known or putative functions in auxin and brassinosteroid signaling, cell division/expansion, polarity growth were identified in selective sweeps particularly for *O. longilobus.* Among these, *IAA18* (OlChr3.1G023990), which plays a role in leaf and root development through the auxin pathway (Uehara et al., 2008), exhibited similar haplotype patterns as *MYB102* (Supplementary Figure 28B, C). Plants have developed complex metabolic systems that generate a wide range of specialized metabolites to adapt to their local environments (Weng et al., 2021). Notably, within the metabolism-related selective genes, we identified three tandem OMT duplicates (OlChr5.1G020250, OlChr5.1G020270, and OlChr5.1G020320) that displayed weak sweep signals in *O. taihangensis*, suggesting the potential significance of metabolite changes during species differentiation (Supplementary Figure 29). Overall, our results shed light on the selective regions associated with *Opisthopappus* species and offer valuable resources for the investigation of genes contributing to genetic differentiation.

### Identification of genomic variants and genes associated with local climate adaptation

A total of 15,470 SNPs and 1,554 InDels were identified for 19 climatic factors and altitude through LFMM, with 5,838 variants co-associated with three or more variables (Supplementary Tables 41 and 42). The Gradient forest (GF) analysis identified three temperature-related variables (Bio4, Bio7, and Bio2) and precipitation seasonality (Bio15) as key determinants of *Opisthopappus* distribution (Supplementary Figure 30A). To address the importance and correlations of environmental factors, we conducted RDA using seven selected variables (Bio1, Bio3, Bio4, Bio12, Bio13, Bio15, and altitude) to mitigate multicollinearity (Supplementary Figure 31A). This analysis identified 4,266 SNPs and 354 InDels overlapping with LFMM results. Among these core adaptive variants, nearly half exhibited multi-trait linkage, with precipitation-related associations (75.52%) being more prevalent than temperature-related ones (46.47%) (Supplementary Figure 32A and Supplementary Table 43). Moreover, these adaptive genetic variations were largely attributable to the seven variables, reflecting clear precipitation-driven changes in spatial distribution (Supplementary Figures 30B, C, and 32B).

Although significant IBD and isolation by environment (IBE) were detected across all variants, partial Mantel tests controlling for geography demonstrated a significant IBE only in adaptive variants, thus indicating that environmental factors predominantly shape adaptive genetic variation (Supplementary Figure 32C, D). To characterize these candidate adaptive variants, we estimated and compared the *F*_ST_ between species/lineages based on adaptive and neutral variants. Remarkably, *F*_ST_ for adaptive variants was significantly lower between the two *Opisthopappus* species but higher between intra-specific lineages. This finding aligns with the significant climatic differences between lineages (Supplementary Figure 31B), providing evidence that local adaptation is driving their genomic differentiation (Supplementary Figure 32F-H). Additionally, PCA analysis based on core adaptive variants revealed a distinct population structure, with stronger differentiation between lineages and reduced dependence on geographical clustering (Supplementary Figure 17E). The enrichment of core adaptive variants in genic regions and UTRs (Supplementary Table 33) suggests that both protein-coding and regulatory alterations are key drivers of environmental adaptation in *Opisthopappus*.

A total of 4,437 core genes were identified with a distribution skewed toward chromosome ends (Figure 4A, Supplementary Figure 33A, and Supplementary Table 44). Functional analysis implicated these candidate genes in key pathways for climatic adaptation, involved in biological processes such “cell wall organization”, “regulation of hormone levels”, “response to desiccation”, “methylation”, “regulation of metabolic processes”, “response to high light intensity”, “transcription factor activity”, and “terpene synthase activity” (Supplementary Figure 33B and Supplementary Table 45). Among them, the *JOX1* gene (OlChr9.1G050340) stood out, being associated with three temperature variables (Bio2, Bio3, and Bio7; Figure 4B). It encodes a jasmonate-induced oxygenase that modulates JA signaling to balance growth and defense (Zhu et al., 2021). To examine allele frequency patterns, a non-synonymous adaptive SNP in *JOX1* was selected as an illustrative example. The A allele was almost fixed in OL_HB populations located at the northernmost extent of their range in the Taihang Mountains, which experience large diurnal and annual temperature fluctuations, while the T allele was more prevalent in OT_N populations with low isothermality (Figure 4D and Supplementary Figure 31B). We also identified a set of precipitation-associated genes, including *CRK3*, *NF-YA2*, *BBX8*, *CYP82G1*, *RGA1*, *WRKY4*, and *MYB73* (Figure 4A).

**Figure 4.**
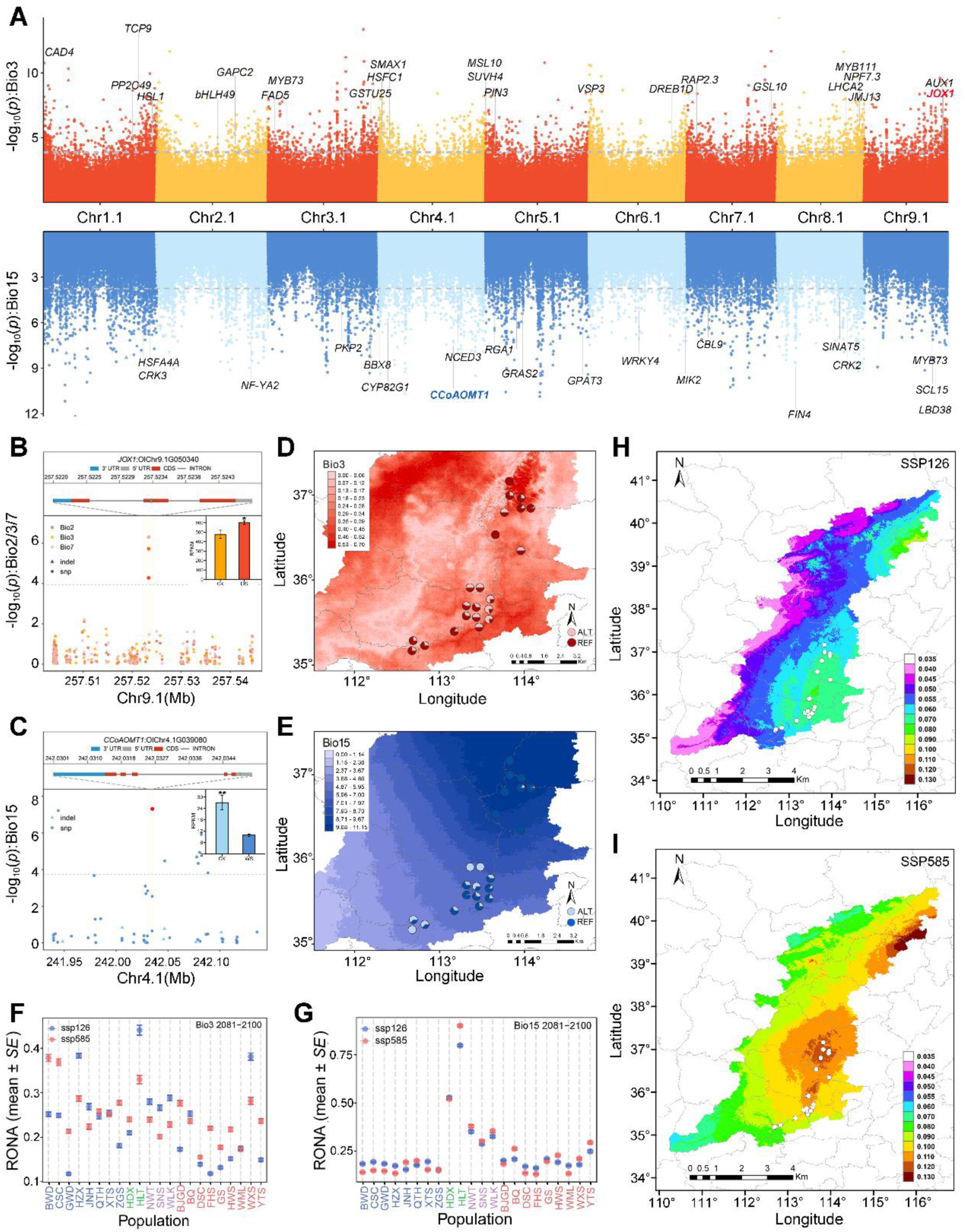
Local adaptation and genomic vulnerability to future climate change in *Opisthopappus* genus. **A,** Genome-wild scans identify variants associated with Bio3 (Isothermality) (red, upper panel) and Bio15 (Precipitation seasonality) (blue, lower panel). SNPs and InDels are represented by points and triangles, respectively. Dashed horizontal lines are the significance thresholds with FDR correction. Candidate adaptive genes are labeled in the plot. **B, C,** Local Manhattan plots highlighting the preventative adaptive genes *JOX1* (**B**) and *CCoAOMT1* (**C**). Upper panels show their gene structure with proposed functional adaptive variants marked by green vertical lines. Inset: Bar plots showing the expression patterns of *JOX1* and *CCoAOMT1* under drought (DS) and waterlogging (WS) treatments, respectively. Two-tailed Student’s *t*-test was used to determine the significance (* *P* < 0.05; ** *P* < 0.01). **D, E,** Allele frequencies of candidate adaptive SNP (OlChr9.1 257523353; nonsynonymous, c.T724A: p.C242S) and InDel (OlChr4.1 242034948; 5’ UTR c.-106_-105insC) across the 22 *Opisthopappus* populations, corresponding to the sites shown in upper panels of **B** and **C**, respectively. Areas with darker color are predicted to experience more substantial climate change from current (1970-2000) to future (2081-2100) under SSP585 scenarios. **F, G,** Comparison of RONA values across the 22 populations under SSP126 and SSP585 in 2081-2100. Error bars represent the standard error (*SE*) of the average RONAs calculated from four climate models for Bio3 (n = 1,168 variants) and Bio15 (n = 1,824 variants), respectively. **H, I,** Predicted genetic offsets to future climate change under SSP126 and SSP585 in 2081-2100 across the Taihang Mountain range (n = 200,824 grids). Circles on the map indicate the 22 sampled populations.

A notable example is the *CCoAOMT1* gene (OlChr4.1G039080), which encodes caffeoyl CoA 3-O-methyltransferase, a lignin-related enzyme involved in abiotic and biotic stress responses (Yang et al., 2017). This gene was linked to precipitation seasonality (Bio15) and exhibited distinct selection in *O. taihangensis* (Figure 4C and Supplementary Figure 34C-F). A significant 1 bp deletion in the 5’ UTR of *CCoAOMT1* was identified: the A allele was prevalent in OL_HB populations characterized by high precipitation seasonality, while the AG allele was more common in populations with constrained intra-annual precipitation variability (Figure 4E and Supplementary Figure 31B). Interestingly, the expression of *JOX1* and *CCoAOMT1* was induced in *O. longilobus* under both waterlogging and drought stresses, suggesting their critical roles in mediating adaptation to the variable hydrological conditions of cliff environments (Supplementary Figure 34A, B). Several genes, including *TPS1*, *PRX*, and *DREB1C*, were highlighted for their potential pleiotropic roles in adaptation to temperature and precipitation changes, supported by strong linear correlations between allele frequency and phenotypic variation (Supplementary Figures 34A, 35, and 36).

### Genomic vulnerability of *Opisthopappus* species to future climate changes

We employed two methods, the single-locus-based RONA (risk of non-adaptedness) and the multi-locus-based GF, to quantify the genetic vulnerability of *Opisthopappus* populations across different climate change scenarios. The results revealed substantial variation in RONA estimates across both environmental variables and populations. On average, RONA analysis projected an increasing trend over three consecutive 20-year periods (2041-2100), with a more pronounced increase under the high-emission scenario (SSP585) than under the low-emission scenario (SSP126) (Figure 4F, G; Supplementary Figure 37). Populations in regions facing more drastic environmental shifts were expected to exhibit higher RONA values. Specifically, we found an ambiguous tendency that OL_HB populations exhibited higher vulnerability to precipitation fluctuations, whereas OT_N populations were more sensitivity to temperature changes. The OT_S populations had substantially higher RONA values for precipitation-related variables than for temperature variables. In particular, the HLT population within the OL_SX group was predicted to be highly maladapted to expected climate changes across most scenarios (Supplementary Figures 38, 39; Supplementary Table 46).

GF is an advanced genomic offset prediction framework that enables to model the associations between the composite effects of many putatively adaptive locus to multiple climate variables simultaneously (Bernatchez et al., 2024). Consistent with RONA, the spatial genomic offset estimates increased with the severity of greenhouse gas emissions and the temporal shifts projected from 2041 to 2100 (Figure 4H, I; Supplementary Figure 40A-D). A comparative analysis of genetic offsets across the geographic range of *Opisthopappus* revealed uneven environmental selection pressures. Specifically, *O. longilobus* populations were more vulnerable to climate change than *O. taihangensis* populations, especially under the high-emission SSP585 scenario from 2081 to 2100 (Supplementary Figure 40E, F; Supplementary Table 47).

### Establishing bridge hybrids for investigating the biosynthetic pathway of linarin

Among *Opisthopappus* and its closely related genera accessions, *O. longilobus* stood out with the highest linarin content (30.66 mg/g), whereas cultivated chrysanthemums showed minimal accumulation, with ‘Jinling Zipao’ being the lowest (Supplementary Tables 49-51; Supplementary Figure 41B, D). Targeted metabolomic analysis of *O. longilobus* and ‘Jinling Zipao’ indicated linarin was a key differential metabolite (Supplementary Figure 41C, D; Supplementary Table 50). To elucidate the biosynthetic pathway of linarin, we successfully obtained an intergeneric offspring F_1_(1-3) between ‘Jinling Zipao’ and *O. longilobus* (Supplementary Figure 41; Supplementary Table 51). Subsequent metabolic flux analysis enabled us to establish a linear pathway: apigenin → 4’-O-methylation → 7-O-glucosylation → linarin, which was supported by integrating NDP scores with correlation analysis (Figure 5A, F; Supplementary Figure 41F; Supplementary Table 50). The findings revealed that linarin (metabolite 4, m/z 593.2 [M+H]+, m/z 285.0, and m/z 447.1) was exclusively present in *O. longilobus*, with its close related metabolite of acacetin-7-O-glucoside (metabolite 3, m/z 447.1 [M+H]+, m/z 285.1), which lacks glycosylation, suggesting it is a critical pathway intermediate. Further analysis demonstrated strong correlations among acacetin-7-O-glucoside, acacetin (metabolite 2, m/z 285.1 [M+H]+, m/z 270.1), and apigenin (metabolite 1, m/z 271.1 [M+H]+, m/z 153.0) (Figure 5A; Supplementary Figure 42).

**Figure 5.**
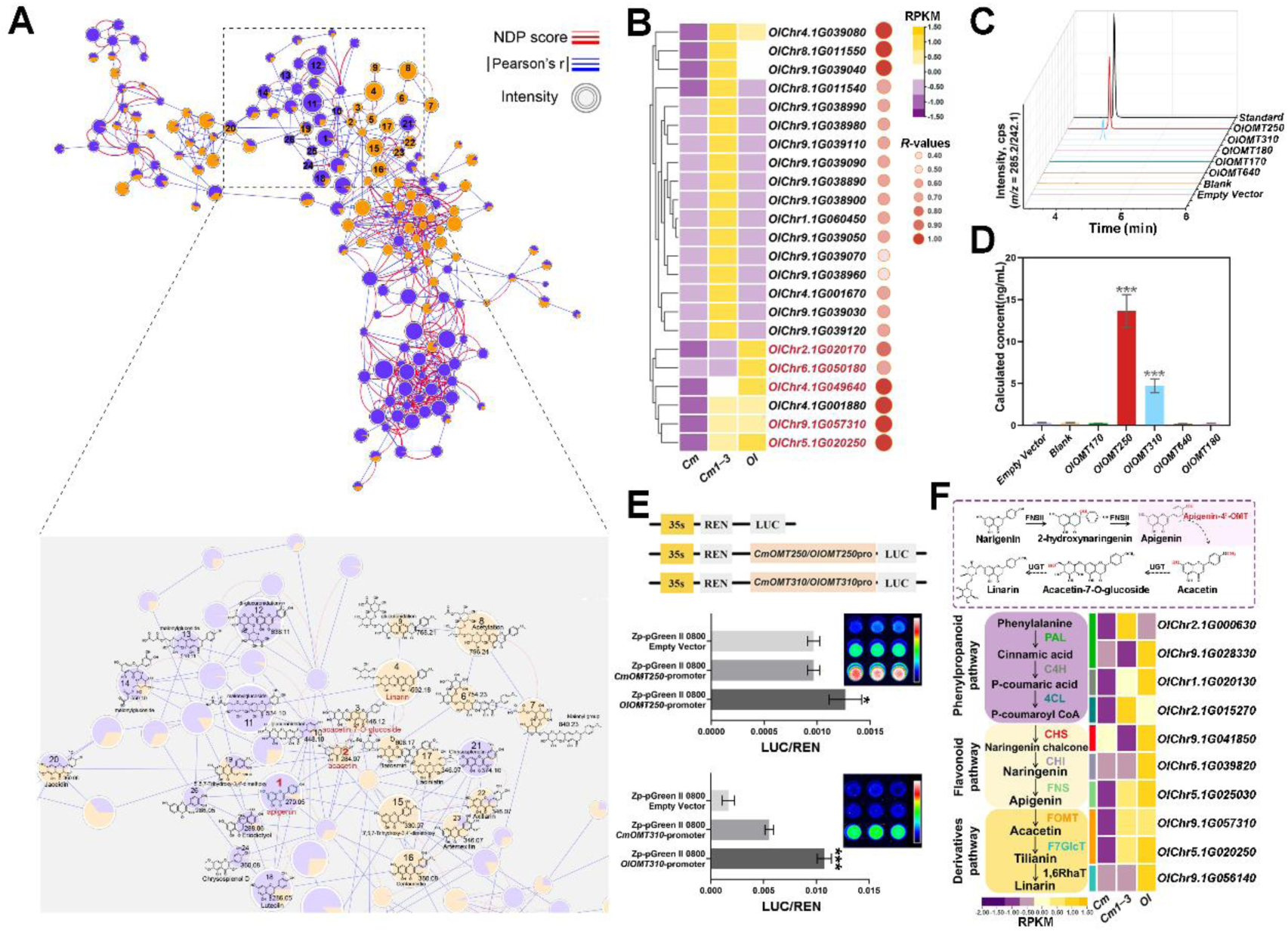
Identification and verification of key OMT genes involved in flavonoid biosynthesis of *O. longilobus*. **A,** Spectral similarity was evaluated using the Normalized Dot Product (NDP, threshold = 0.5) and Neutral Loss (NL). The resulting similarity matrix was visualized as a molecular network in Cytoscape, where edge thickness represents the NDP (red) or NL (blue) scores, node colors indicate the identified compounds. Purple and yellow nodes represent metabolites identified in cultivated chrysanthemum ‘Jinling Zipao’ and *O. longilobus*, respectively. The node size reflects the relative abundance of each metabolite. **B,** Heatmap of RPKM expression levels of OMT genes in *O. longilobus* (Ol), *C. morifolium* cv. ‘Jinling Zipao’ (Cm), and F1 hybrid (Cm1-3), excluding genes with FPKM < 1. The correlation coefficients (*R*-values) represent associations between gene expression and metabolite accumulation. Genes highlighted in red were selected for experimental validation. **C,** Functional characterization of the representative OMT*s* using apigenin as the substrate. Data were from three independent assays. **D,** Statistical analysis of acacetin content. Results are shown as means ± *SD* (n = 3 biological replicates). **E,** Promoter activity of *OMT250* and *OMT310* genes in *O. longilobus* and *C. morifolium* cv. ‘Jinling Zipao’. **F,** Expression patterns of structural genes involved in flavonoid biosynthesis. phenylalanine ammonia-lyase (PAL), cinnamate-acid-4-hydroxylase C4H (C4H), 4-coumarate CoA ligase (4CL), chalcone synthase (CHS), chalcone isomerase CHI (CHI), flavone synthase II (FNSII), flavonoid O-methyltransferase (FOMT), flavonoid 7-O-glucosyltransferase (F7GlcT), and 1,6-rhamnosyltransferase (1,6RhaT).

This pathway was fully active in *O. longilobus* and partially retained in the F_1_ hybrid, whose metabolic profiles closely resembled the male parent (Supplementary Table 50). In ‘Jinling Zipao’, we detected neither acacetin nor its downstream products but instead identified 3’,5,7-trihydroxy-3,4’-dimethoxyflavone (metabolite 19) (Figure 5A). This finding suggests that methoxytransferase activity is not entirely abolished but is specifically deficient in the 4’-O-methyltransferase required to convert apigenin to acacetin, resulting in a pathway bottleneck at the apigenin stage (Supplementary Figure 41F). A comprehensive genome-wide search identified 98 OMT genes, of which 29 were specifically expanded in *O. longilobus* (Supplementary Figure 43A; Supplementary Tables 26 and 52). Among these, five OMT genes exhibited expression levels significantly correlated with linarin accumulation (Figure 5B; Supplementary Figures 43B and 44). By integrating transcriptome-metabolome correlations, we reconstructed the upstream structural genes and the downstream flavonoid modification steps involved in linarin biosynthesis (Figure 5F). Prokaryotic expression assays and transgenic experiment confirmed that *OlOMT250* (OlChr5.1G020250) and *OlOMT310* (OlChr9.1G057310) catalyzed the conversion of apigenin to acacetin (Figure 5C, D; Supplementary Figure 47), thereby validating their roles as functional 4’-O-methyltransferases.

Homolog analysis and *in vitro* enzyme assays indicated the differences in linarin accumulation are not attributable to variations in OMT protein sequences (Supplementary Figures 45A, B and 46). Instead, marked divergence in the *OMT250*/*OMT310* regulatory sequences was observed between *O. longilobus* and ‘Jinling Zipao’, with the promoters from *O. longilobus* exhibiting higher transcriptional activity (Figure 5E; Supplementary Figure 45C, D), suggesting that promoter sequence variations drive the differential accumulation of acacetin and linarin. Docking analysis revealed that both ligands were positioned within the catalytic pocket, forming stable hydrogen bonds between the 4’-hydroxyl group of apigenin and S-adenosylmethionine. The shorter bond length observed in OMT250 (3.3 Å) likely accounts for its enhanced catalytic activity (Supplementary Figure 48A, B). Additionally, molecular dynamics simulations showed a reduced radius of gyration, an increase in hydrogen bonds, and a decrease in solvent-accessible surface area in the ligand-bound complexes, provide further evidence for the selective substrate interaction at the 4’ position during catalysis (Supplementary Figure 48C-H).

## Discussion

As an endangered species, *Opisthopappus* inhabits specialized cliff habitats, facing threats that necessitate urgent conservation research. In this study, we present a conservation genomics framework by integrating the first haplotype-resolved genome assembly of *O. longilobus* with population and conservation genomic analyses. This integration provides valuable evolutionary insights and delivers a critical roadmap for securing the species’ long-term persistence under projected climate change.

### A genomic perspective on the phylogenetic history and adaptive landscape of Opisthopappus

The taxonomic placement of *Opisthopappus* remains controversial. Historically, it has often been classified as a member of *Chrysanthemum*, supported by morphological similarity, intergeneric hybridization, and evidence from low-copy nuclear genes and nrDNA/ITS data (Shih, 1979; Zheng et al., 2013; Shen et al., 2021). Our phylogenetic analysis of nuclear and chloroplast genomes collectively confirmed *O. longilobus* as an independent lineage, revealing a closer nuclear relationship with the *Artemisia*-*Chrysanthemum* clade (Figures 1C, 2A; Supplementary Figure 10), which is consistent with findings from the genomic study of *O. taihangensis* (Wang et al., 2025b). This provides a clearer and more robust phylogenetic framework, enhancing our understanding of the relationships within Asteraceae.

Genome polyploidization and LTR retrotransposon proliferation drive plant genome evolution and can enhance adaptation to environmental change (Jiao et al., 2011; Galindo-González et al., 2017). *O. longilobus* underwent pronounced genome remodeling in the Miocene, characterized by extensive TD/PD duplications, and experienced a pulse of LTR-RT expansion during a late Pleistocene interglacial (∼110 ka). The Miocene TD/PD burst coincided with the prolonged uplift and intensified exhumation of the Taihang Mountains, as well as a long-term shift toward cooler and drier climates (An et al., 2006; Ge et al., 2012; Cao et al., 2015; Zhang and Zhang, 2020). This consistency suggests that TD/PD-driven dosage alterations and regulatory rewiring may have facilitated adaptation (Supplementary Figure 11F). In contrast, LTR-RT activation likely reflects a shorter-term response to Quaternary climate fluctuations.

From the last interglacial to the last glacial maximum, as the climate transitioned from relatively warm-humid to colder condition, the potential distribution range of *O. longilobus* generally contracted (Liu et al., 2023); thus, this episodic LTR expansion may be associated with adaptive responses under rapid environmental change (Supplementary Figure 12B-D).

### Climate-driven divergence and conservation strategies of *Opisthopappus* plants

The high-quality reference genome for *O. longilobus* and in-depth resequencing data together support comprehensive evolutionary genomic analyses in *Opisthopappus*. Multiple lines of evidence suggest that the two *Opisthopappus* species exhibit significant genetic differentiation, and *O. taihangensis* was diverged from *O. longilobus* approximately 139.5 thousand years ago (Figure 3; Supplementary Figures 18, 19, 22B and 23). By integrating demographic history and paleoclimatic data, we speculate that the penultimate glaciation (130-300 kya) and concurrent tectonic activity in the Taihang Mountains likely facilitated the speciation of these two *Opisthopappus* species by promoting geographic isolation and adaptive divergence (Figure 3E). Our field surveys together with existing literature (Yang, 2021; Wang et al., 2024) indicate that the southern Taihang Mountains have experienced more severe human-induced destruction, which may be a critical factor in the recent rapid decline of *O. taihangensis* compared to *O. longilobus* (Figure 3F).

Understanding the genetic variation underlying plant adaptation is critical for facilitating the climate resilience of key species, an imperative intensified by global change (Lasky et al., 2022). Based on the core adaptive SNPs/InDels identified here, our genetic offset analysis revealed that while *O. longilobus* exhibits higher genetic diversity (Figure 3D), it faces a heightened risk of local extinction due to future climate change, particularly under more severe carbon emission scenarios (Figure 4 H, I, and Supplementary Figure 40), emphasizing the need for prioritizing conservation efforts. Additionally, we also identified populations such as WLK (Wulongkou, Henan Province) and BJGD (Baijinggudao, Shanxi Province) with enhanced resilience to climate fluctuations, which may serve as candidates for targeted breeding programs.

### Conservation and divergence of apigenin 4**’**-O-methyltransferases

*Opisthopappus* is characterized by exceptionally high accumulation of medicinal flavonoids, particularly linarin, which consequently faces sustained anthropogenic disturbance. Systematic elucidation of the linarin biosynthetic pathway offers a molecular foundation for reducing dependence on *Opisthopappus* populations. Interestingly, the apigenin 4’-O-methyltransferase of *Opisthopappus*, despite sharing less than 40% sequence identity with homologs from phylogenetically distant taxa (e.g., soybean, liverwort, safflower, and *Chrysanthemum*), retains a functionally conserved catalytic pocket. (Kim et al., 2005; Umezawa et al., 2013; Liu et al., 2017; Hou et al., 2024b). In this pocket, hydrophobic residues (Phe, Trp, Ile, Leu) stabilize the apigenin positioning, whereas polar residues (Asn, His) mediate the 4’-OH group, thereby supporting efficient methylation (Supplementary Figure 43C and 48A, B, I and J).

We introgressed the high-linarin trait from *O. longilobus* into cultivated chrysanthemum via distant hybridization and developed stable lines that retained elevated linarin content along with efficient vegetative propagation (Supplementary Figure 41A and 47C, D). This strategy simultaneously improves flavonoid profiles of cultivated chrysanthemum and alleviates pressure on wild *Opisthopappus* populations. To date, the functional relationship between flavonoids and environmental adaptation in *Opisthopappus* remains poorly understood. Notably, the 190 core adaptive genes and 629 LFMM-based adaptive genes that overlap with Asteraceae-specific expanded genes in *O. longilobus* were both enriched in the molecular function term related to O-methyltransferase activity (Supplementary Figure 33C; Supplementary Tables 48 and 54), underscoring their dual role in local environmental adaptation and linarin accumulation. However, further validation experiments are still required.

In summary, the high-quality *O. longilobus* genome highlights its distinct evolutionary position within Asteraceae and demonstrates that its adaptive evolution is likely driven by genomic expansion via TD/PD duplication and TE proliferation. The integrated population and ecological genomic analysis reveal that the pronounced genetic divergence between *O. longilobus* and *O. taihangensis* is shaped by both geographic isolation and environmental heterogeneity. The adaptive genes and vulnerable populations uncovered here are key genetic assets for guiding future conservation and breeding efforts. Moreover, we verify the promoter sequence diversity of *OMT250* and *OMT310* are responsible for the differential accumulation of linarin in *C. morifolium* and *O. longilobus*. This study establishes a scientific foundation for the conservation and utilization of cliff-dwelling genus *Opisthopappus*, offering a key paradigm for the climate-resilient protection of other endangered plants.

## Methods

### Plant materials and genome sequencing

For whole-genome sequencing and assembly, we selected a diploid (2n = 2x = 18) plant of *O. longilobus* collected from Xingtai City, Hebei Province, China as the material (37.29°N, 114.51°E; altitude 800 m; Supplementary Figure 1), which was preserved at the Chrysanthemum Germplasm Resource Preservation Center of Nanjing Agricultural University (accession ID: NAU158). Aseptic plantlets were regenerated on Murashige and Skoog (MS) medium, and young leaves were harvested for genomic DNA extraction using a magnetic bead-based method. Detailed information of DNBSEQ-2000 short-read sequencing (129.87×), PacBio Sequel II long-read sequencing (82.95 ×), and Hi-C library construction and sequencing (136.06 ×) are outlined in Supplementary Note 1.

### Haplotype-resolved genome assembly

Genome assembly of *O. longilobus* was performed using the aforementioned sequencing dataset, with all sequencing reads subjected to microbial contamination screening and removal of organellar sequences before assembly. The HiFi and Hi-C reads were jointly assembled using Hifiasm v0.19.5 (Cheng et al., 2021), with the Hi-C assisted phasing mode activated (parameters: -i -h1 hic_reads1.fq.gz -h2 hic_reads2.fq.gz). We employed the built-in Purge-dups function of Hifiasm to handle heterozygous regions and eliminate redundant haplotypic. This process resulted in the production of one primary genome and two haplotype draft contigs. Hi-C reads were aligned to the primary contig assembly using BWA v.0.7.12 (Li and Durbin, 2009), followed by retention of only uniquely mapped paired-end reads for subsequent analysis. The contigs were clustered, ordered, and oriented using Juice v1.5.6 (Durand et al., 2016b) and 3D-DNA v180922 (parameters: -m haploid -r 0) (Dudchenko et al., 2017), after which they were integrated to generate pseudochromosomes. Then the assemblies were manually refined using Juicebox Assembly Tools v2.15.07 (Durand et al., 2016a) to correct for potential mis-joins and orientation errors introduced by automated scaffolding. We anchored approximately 99.67% of each haplotype assembly onto 18 pseudomolecules following Hi-C–guided scaffolding and manual curation, producing two chromosome-level haplotype assemblies (Hap1 and Hap2). The quality of the *O. longilobus* genome assembly was systematically evaluated through multiple approaches: assessing continuity with the LTR Assembly Index (LAI), verifying accuracy by mapping PacBio long reads and RNA-seq reads back to the assembled genome, examining completeness with BUSCO v5.2.2 (Waterhouse et al., 2018), and conforming haplotype phasing accuracy using Merqury v1.3 (Rhie et al., 2020) alongside Hi-C contact map-guided curation in Juicebox. Genome features were summarized using a 1 Mb non-overlapping sliding window and visualized using Circos software v0.69 (Krzywinski et al., 2009). A more detailed description of the genome assembly methods can be found in Supplementary Note 2.

### Repeat and gene annotation

Repetitive elements were annotated by an integrated approach that merged *de novo* predictions with homology-based searches using RepeatMasker v4.0.7 (Tarailo-Graovac and Chen, 2009), while tandem repeats were detected using Tandem Repeats Finder v4.09.1 (Benson, 1999). Gene prediction was performed using a combined strategy incorporating ab initio predictions, homology-based evidence, and transcriptome data. Specifically, for homology-based annotation, protein sequences from six Asteraceae species were aligned to the *O. longilobus* genome using GeMoMa v1.9 (Keilwagen et al., 2019). In terms of transcriptome-based gene prediction, RNA-seq data from eight tissues (buds, flowers, leaves, stems, and roots) of *O. longilobus* collected at different developmental stages were mapped to the reference genome using HISAT2 v2.1.0 (Kim et al., 2015) and GMAP v2020-06-02 (Wu and Watanabe, 2005). Following integration of protein homology, transcript, and ab initio prediction evidence via GeMoMa, annotation completeness was evaluated using BUSCO. In addition, rRNAs were detected by BLAST v2.2.26 (Altschul et al., 1990) searches against ncRNA datasets from *A. thaliana* and *O. sativa*, whereas miRNAs and snRNAs were identified with INFERNAL v1.1.2 (Nawrocki and Eddy, 2013) and tRNAs were predicted using tRNAscan-SE v1.3.1 (Lowe and Eddy, 1997). For a detailed overview of the methodologies used for genome annotation, please refer to Supplementary Note 3.

### Genome evolution

We first performed the angiosperm phylogenetic reconstruction using orthologous genes from 18 representative plant species with well assembled genome sequences, including monocots *O. sativa* and *Z. mays*; basal angiosperms represented by *A. trichopoda*; core eudicots of *C. quinoa*, *A. coerulea*, *V. vinifera*, *S. lycopersicum*, *D. carota*, and *C. canephora*; as well as nine Asteraceae species from the order Asterales: *H. annuus*, *L. sativa*, *C. cardunculus*, *A. annua*, *A. argyi*, *C. lavandulifolium*, *C. nankingense*, *C. seticuspe*, and *C. morifolium*. To further build a comprehensive phylogenetic tree of Asteraceae, we included additional 14 species including *G. coronaria*, *E. canadensis*, *C. makinoi*, *C. intybus*, *C. endivia*, *A. lappa*, *S. rebaudiana*, *S. sonchifolius*, *T. mongolicum*, *T. kok-saghyz*, *C. tinctorius*, *M. micrantha*, *E. breviscapus*, and *S. taccada* (Supplementary Table 23).

The detailed description of gene family and genome evolution analyses is available in Supplementary Note 4. Briefly, we identified single-copy orthologous genes from the aforementioned plant genomes by OrthoFinder v2.5.4 (Emms and Kelly, 2015), and constructed two maximum-likelihood species trees using MAFFT (Katoh and Standley, 2013), FastTree v2.1 (Price et al., 2010), IQ-Tree v2.2.6 (Minh et al., 2020), and RAxML v8.2.12 (Stamatakis, 2014) programs with 1000 bootstrap replicates. The resulting tree topologies were compared and a representative tree was selected for downstream analyses. We estimated the divergence times using the MCMCtree program within the PAML v4.5 (Yang, 1997). The CAFÉ software v4.2.1 (De Bie et al., 2006) was utilized to analyze gene family evolution based on the phylogenetic tree and orthologous groups. Genes from specific expanded gene families underwent GO and KEGG enrichment analyses. Additionally, positively selected genes were identified using CodeML program in PAML v4.10.7.

The JCVI pipeline was subjected to infer syntenic self-alignment of the assembled *O. longilobus* genome. To detect ancient whole-genome duplication (WGD) events, WGD analysis was conducted using WGDI v0.6.5 (Sun et al., 2022) among *O. longilobus*, *C. morifolium*, and *C. nankingense*. For enhanced accuracy, DupGen_finder (Qiao et al., 2019) was employed to identify gene pairs resulting from WGD events. The values of synonymous substitutions per synonymous site (*Ks*) between these gene pairs were calculated to estimate the divergence times. Additionally, the four-fold synonymous third-codon transversion rate (4DTv) was analyze to date the WGD events using a public script (https://github.com/JinfengChen/Scripts/blob/ master/FFgenome/03.evolution/distance_kaks_4dtv/bin/calculate_4DTV_correction.p l) (Supplementary Note 4).

### Whole-genome resequencing and variant calling

A total of 226 individuals were collected from 28 natural populations throughout the entire distribution range of the *Opisthopappus* genus in the Taihang Mountains, China (Supplementary Table 31). To prevent the risk of sampling cloned individuals and to avoid habitat destruction, individuals from each population were spaced at least 100 meters apart, with only a few samples taken from some populations. Consequently, 66 representative individuals of *O. taihangensis* (OT) and 49 of *O. longilobus* (OL) were selected for genome resequencing on the DNBSEQ-T7 platform (Supplementary Table 32). Following quality control procedures (Supplementary Note 5), high-quality reads were aligned to the *O. longilobus* haplotype 1 genome using BWA-MEM v0.7.17 (Li, 2014) with default parameters. We utilized SAMtools v1.9 (Li et al., 2009a) to sort BAM files and generate index files. PCR duplications were marked by MarkDuplicates function of GATK v4.2.6.1 (McKenna et al., 2010).

Variant calling was executed using a joint calling strategy implemented through GATK’s HaplotypeCaller, CombineGVCFs, GenotypeGVCFs, and SelectVariants modules. The raw SNPs and InDels were subsequently separated and filtered according to the following parameters: “QD < 2.0 || FS > 60.0 || MQ < 40.0 || MQRankSum < -12.5 || ReadPosRankSum < -8.0” for SNPs and “QD < 2.0 || FS > 200.0 || SOR > 10.0 || MQRankSum < -12.5 || ReadPosRankSum < -8.0” for InDels. To reduce the impact of sequencing and mapping biases on the variants, we conducted hard filtration using VCFtools v0.1.17 (Danecek et al., 2011) with the parameters “--minDP 5 --maxDP 300 --minQ 30 --maf 0.05 --max-missing 0.85 --min-alleles 2 --max-alleles 2”. As a result, a total of 27,009,474 SNPs and 3,010,432 InDels were retained for further analysis, and functional annotation of the genetic variants was performed using ANNOVAR software (Wang et al., 2010).

### Phylogenetic inference and population genetic analysis

We initially employed PLINK v1.9 (Chang et al., 2015) with the parameters “indep-pairwise 50 10 0.2” to filter out excessive linkage sites, resulting in 3,808,756 independent SNPs for our phylogenetic and population structure analyses. ADMIXTURE v1.3.0 (Alexander and Lange, 2011) was utilized to infer population structure by testing values of K (ancestor number) ranging from 2 to 10, with the optimal K determined based on the lowest cross-validation (*CV*) error value. To visualize the structural information, we used the R package pophelper v2.3.0 (Francis, 2017). PCA was conducted using PLINK v1.9. Using FastTree v2.1.11 (Price et al., 2010), we investigated the phylogeny of *Opisthopappus* by constructing two ML trees—with and without *A. annua* as an outgroup. Kinship analysis was performed by GCTA v1.93.0 (Yang et al., 2011).

We calculated nucleotide diversity (*π*) and Tajima’s *D* statistics for each group using VCFtools within a 100 kb window and a sliding step of 20 kb. The individual inbreeding coefficient (*F*_IS_) was estimated using GCTA, while heterozygosity was assessed with VCFtools. Additionally, we evaluated the level of inbreeding using the frequency of runs of homozygosity (FROH) to assess the inbreeding level, which was defined as the ratio of the total length of ROH (>100 kb) to the effective length of the genome (Supplementary Note 5). These population genetic parameters were derived from the high-quality SNP set, without prior linkage disequilibrium (LD) filtering.

### Population divergence and gene flow analysis

To investigate the divergence and genomic differences within and between species, we first calculated the fixation index (*F*_ST_) and nucleotide divergence (*D*_XY_) statistics utilizing VCFtools and PIXY v1.2.5 (Korunes and Samuk, 2021) across 100 kb non-overlapping windows. To estimate the divergence times between *O. taihangensis* and *O. longilobus*, we employed the split function in SMC++ (Terhorst et al., 2017).

Treemix v1.13 (Pickrell and Pritchard, 2012) was employed to infer potential gene flow among *Opisthopappus* populations, relying on all LD-pruned SNPs. Each of the assumed 0-15 migration events (m) was run for five iterations, with resampling blocks of 500 SNPs per iteration (-bootstrap -se -k 500 -noss). The optimal number of migration edges was determined and visualized using the R package OptM (Fitak, 2021). To avoid bootstrapped TreeMix runs having a standard deviation of 0, we initially used a subset comprising 80% of LD-pruned SNPs for the TreeMix and OptM runs (Lofgren et al., 2022). Subsequently, we re-ran TreeMix at the optimal value of m = 6, incorporating all pruned SNPs as input.

Dsuite v0.3 (Malinsky et al., 2021) was utilized to conduct the ABBA-BABA test for detecting introgression among populations. The *D*-statistic and *f*4 ratio were estimated using the Dtrios module, with *A. annua* serving as the outgroup. To derive the species tree, we employed TreeMix analysis while assuming zero migration events. The significance of each test was assessed using 20 jackknife resampling runs, and only those cases with a *Z*-score greater than 3 and an FDR-adjusted *P*-value of less than 1E-3 were deemed significant gene flow events. The *D*-statistic and *f*4 ratio are particularly effective at identifying recent introgression (Bjorner et al., 2024). Consequently, we further utilized the Fbranch program to infer deeper introgression events for each phylogenetic branch on the species tree, with the results visualized using the “dtools.py” script.

### Genome scanning for selective signals

To identify potential selective signals between *O. taihangensis* and *O. longilobus*, we employed three methodologies: *π* ratio, *F*_ST_, and XP-CLR, based on 27,009,474 high-confidence filtered SNPs. *π* ratios (*π*_OL_/*π*_OT_ and *π*_OT_/*π*_OL_) and *F*_ST_ values were calculated using VCFtools with a 100-kb window and a step size of 20 kb along the *O. longilobus* haplotype1 genome. The cross-population composite likelihood ratio (XP-CLR) test was conducted with the Python version of XP-CLR v1.1.2 (https://github.com/hardingnj/xpclr), utilizing the parameters “--rrate 3.64e-9 --ld 0.95 --maxsnps 200 --size 100000 --step 20000”. To enhance the accuracy, we defined regions that ranked in the top 5% of the highest values from any two of the three methods as putative selective sweeps.

### Demographic history inference

We utilized PSMC v0.6.5 (Li and Durbin, 2011) and Stairway Plot v2.1 (Liu and Fu, 2020) to examine the changes in effective population size (*N*e) of *Opisthopappus* over time. For the PSMC analysis, we randomly selected four individuals with “pure” genetic backgrounds (Q > 0.9999 in structure analysis) and high sequencing depth from both *O. taihangensis* and *O. longilobus* populations. The bam files were processed using the bcftools mpileup, bcftools call, and fq2psmcfa pipeline before conducting the PSMC analysis with parameters set to “-N25 -t15 -r5 -p 4+25*2+4+6”. The Stairway Plot method, which incorporates genetic information from all samples, is effective in inferring recent demographic dynamics. For this analysis, we first generated a folded (ancestor allele unknown) site frequency spectrum (SFS) from the VCF file containing all SNPs using easySFS (https://github.com/isaacovercast/easySFS). In both methods, we assumed a generation time of one year and a mutation rate of 8.25e-9 per site per year (Song et al., 2018).

### Genotype-environment associations analysis

A total of 19 bioclimatic variables (Bio1-Bio19) at a resolution of 30 seconds (∼1 km) for each sampling location were downloaded from the WorldClim v2.1 database (Supplementary Table 41; Supplementary Note 6). Two distinct approaches were employed for the genome-wide identification of environment-associated genetic variants. Only high-quality pruned variants with a minor allele frequency (MAF) greater than 0.1 and a genotype missing rate of less than 0.05 were retained for the genotype-environment association (GEA) analysis, resulting in a final dataset comprising 1,353,182 SNPs and 132,918 InDels.

Initially, we conducted the GEA analysis for the 19 bioclimatic variables as well as altitude using a univariate latent-factor linear mixed model (LFMM) through the R package LEA v3.20.0 (Frichot and François, 2015). Four latent factors, accounting for population structure, were defined in the LFMM analysis (K = 4), according to the optimal number of ancestry clusters inferred by ADMIXTURE. For each environmental variable, the Gibbs sampler algorithm was executed five times, each for a duration of 5000 cycles, following a burn-in period of 5000 cycles. *P*-values from all five runs were subsequently averaged for each variant and adjusted for multiple tests using a Benjamini-Hochberg false discovery rate (BH-FDR) correction with a significance threshold of 5%.

We proceeded to conduct a redundancy analysis (RDA) to identify adaptive variants, which is recognized as an effective multivariate GEA method known for its low false-positive rates (Sang et al., 2022). We selected six significant environmental variables (Bio1, Bio3, Bio4, Bio12, Bio13, Bio15, and Altitude; Supplementary Note 6) that demonstrated pairwise correlation coefficients |*r*| < 0.6 before performing RDA using the R package vegan v2.7.1 (Oksanen et al., 2013). Outliers that fell within the tails of the 3 ± *SD* cutoff were identified as candidate environmental associations. Genetic variants that overlapped between both approaches were classified as “core adaptive variants” and then subjected to functional annotation. Local adaptive genes were extracted from the 15-kb genomic regions surrounding each core adaptive variant, based on the LD decay distance estimated by PopLDdecay software (Zhang et al., 2019).

### Genomic offset assessment under future climate chang

We collected current environmental data from 1970 to 2000 and projected conditions for three future periods (2041-2060, 2061-2080, and 2081-2100) across the Taihang Mountain region. This was done under two shared socioeconomic pathways (SSPs) emission scenarios, SSP126 and SSP585, utilizing four global climate models: ACCESS-CM2, BCC-CSM2, CMCC-ESM2, and GISS-E2-1-G, as detailed in Supplementary Note 6. To assess genomic offset, a measure of climate vulnerability, we employed the risk of non-adaptedness (RONA) and gradient forest (GF) approaches. To ensure adequate statistical power and avoid bias, we excluded populations containing only a single individual for the genomic offset analysis.

RONA measures the theoretical average change in allele frequency needed to align with future climatic scenarios. This metric is derived by regressing the allele frequencies of selected loci against chosen environmental variables, with the resulting regression coefficients informing predictions of expected allele frequencies for those loci under future conditions (Karunarathne et al., 2024). Following the methodology outlined by Sang et al. (2022), we calculated RONA values based on core adaptive variants for each locus, population, and environmental variable using R v4.3.6. A higher RONA value indicates an increased risk of extinction under projected climate change scenarios.

GF method was implemented using the R package gradient Forest v0.1.37 (Ellis et al., 2012), serving as a complementary approach to RONA. This method predicts the genomic offset of locally adapted populations under future climate scenarios by capturing the nonlinear relationships between current climatic variables and allele frequencies. We first calculated allele frequencies for 22 populations using 4,620 core adaptive variants, and subsequently built a GF model (ntree = 500, nbin = 01, corr.threshold = 0.5) to estimate the genetic offset using 19 environmental variables. The genetic offset was defined as the Euclidean distance between the genomic compositions of current and projected future climates, and we visualized the spatial distribution of local maladaptation using ArcGIS Desktop v10.7.

### Identification of candidate genes in linarin biosynthetic pathway

UPLC was employed to quantify linarin levels across wild and cultivated chrysanthemum accessions, revealing markedly higher accumulation in *O. longilobus* and the lowest levels in the cultivated chrysanthemum ‘Jinling Zipao’. These contrasting materials were subsequently selected to generate intergeneric hybrids for downstream biosynthetic pathway analysis. Flavonoid-targeted metabolomics using UPLC-ESI-QQQ-MS/MS was applied to identify linarin-related flavonoids, including potential intermediates and branch metabolites. UPLC–TripleTOF 5600+–based untargeted metabolomics was performed, with metabolites clustered by structural similarity using the NDP algorithm (NDP) and prioritized by neutral loss (NL) scoring for pathway reconstruction. UPLC-ESI-TOF-MS/MS fragmentation patterns were inspected to further confirm metabolite identities.

O-methyltransferase (OMT) proteins in *O. longilobus* were identified using a combination of homology-based BLASTP searches and HMMER v3.2.1 analyses with the conserved Pfam domains (PF00891, PF08100, and PF01596). Gene expression patterns across *O. longilobus*, ‘Jinling Zipao’, and their hybrid were analyzed, from which genes exhibiting expression levels significantly correlated with linarin content (*r* > 0.6) were retained as candidates for functional analysis. A more detailed protocol for this can be found in Supplementary Note 7.

### Enzyme assays

OMT enzyme assays were performed in 100 μL reactions. Each reaction consisted of 200 mM Tris-HCl buffer (pH 7.5), 0.5 mM S-adenosyl-L-methionine (SAM), 4 mM dithiothreitol (DTT), and 0.2 mM apigenin (dissolved in DMSO), along with 4 μg of purified recombinant protein. The final volume was adjusted to 100 μL with double-distilled water (ddH₂O). The mixture was incubated at 37°C for 30 min with gentle shaking, and the reactions were terminated by adding an equal volume of quenching solution (acetonitrile containing 10% methanol and 10% DMSO). Heat-denatured protein expressed from the empty pCold II vector served as a negative control. Following centrifugation at 12,000 rpm for 10 min, the supernatant was analyzed using LC-MS to detect acacetin products.

### Transgenic validation of key candidate OMT genes

The sequences of *OlOMT310* and *OlOMT250* were cloned into the pORE-R4-35AA vector to generate overexpression constructs. Due to the lack of a stable transformation system for the chrysanthemum cultivar ‘Jinling Zipao’, overexpression was conducted in another cultivar ‘Nannong Fencui’, which also exhibits low linarin content (0.03 mg/g). For gene silencing, gene-specific sense and antisense fragments of *OlOMT310* and *OlOMT250* were inserted into the pCVA vector. These vectors were introduced into *Agrobacterium tumefaciens* strain GV3101 (Tang et al., 2010). Stable overexpression lines were obtained by leaf disc transformation (Geng et al., 2025), whereas transient RNA interference in *O. longilobus* was achieved by vacuum penetration using mixed *A. tumefaciens* suspensions carrying pCVA or pCVA-amiR-*OlOMT250*/*OlOMT310* together with pCVB. Leaves from the rooted seedlings were collected for DNA extraction, and positive seedlings were identified through PCR analysis.

The control and transgenic materials were incubated at 37°C overnight, followed by drying at 60°C to constant weight. Dried tissues were ground into powder, and 0.1 g of each sample was extracted with methanol for metabolite analysis. Metabolic products generated in these plant materials were subsequently analyzed using targeted quantification on a Waters ACQUITY UPLC H-Class system coupled to a Triple Quad 6500+ mass spectrometer (Supplementary Note 8).

### Promoter sequence analysis and dual luciferase transactivation assays

Pairwise alignments of the *OMT250* and *OMT310* promoter sequences between *O. longilobus* and *C. morifolium* cv. ‘Jinling Zipao’ were generated using N-ESPript v0.9. The variants located in the promoter region (2 kb upstream of the translational start codon) of *OMT250*/*310* were amplified and cloned into the pGreenII 0800-LUC vector, which harbors both the firefly luciferase (LUC) and Renilla luciferase (REN) reporter genes. Plasmid constructs were introduced into ‘Jinling Zipao’ protoplasts through PEG-Ca²⁺-mediated transfection, with the empty pGreenII 0800-LUC vector serving as a negative control. After 18 h, LUC and REN activities were assessed using the CCD imaging system (Tanon 5200) in conjunction with a Dual-Luciferase Reporter Assay Kit (Vazyme). LUC activity was normalized against REN activity. The primers utilized for vector construction are listed in Supplementary Table 55.

## Data availability

All sequencing data utilized for genome assembly and annotation, population genetics and transcriptome, along with details regarding the assembled haplotype-resolved genome of *O. longilobus*, have been deposited at the National Genomics Data Center (NGDC; https://ngdc.cncb.ac.cn/). These raw sequencing data can be accessed in the Genome Sequence Archive (GSA; project accession CRA048684), while the haplotype-resolved chromosome-level genome assembly is available in the Genome Warehouse (GWH; accession GWHERQKxxxx). Source data are provided alongside this paper.

## Author contributions

F.C. and J.J. designed and conceived the research. J.J., W.F., Z.W., S.C., and A.S. architected the sequencing strategy. J.J., W.C., and H.W. collected the wild germplasm resources. R.J., Y.W., Y.Z., and X.S. maintained and prepared the sequencing samples. R.J., A.S., and Z.Y. analyzed the genome evolution. J.S. and F.Z. performed the population genomics analyses. M.W., Z.W., and R.J. performed metabolomics raw data acquisition and analysis. R.J., Y.W., J.H., and L. M. conducted experiments. R.J. and S.J. wrote the draft manuscript. F.C., J.J., J.S., J.X., W.F., Z.W., A.S., and H.W. revised the manuscript. All authors approved the final manuscript.

## Acknowledgments

We thank Dr. Yuehua Ma (Central laboratory of College of Horticulture, Nanjing Agricultural University) and Dr. Shaoyan Lin (State Key Laboratory of Crop Genetics and Germplasm Innovation, Nanjing Agricultural University) for assistance in using UPLC-MS/MS, and Bioinformatics Center of Nanjing Agricultural University. We thank Dr. Yunpeng Zhao (College of Life Sciences, Zhejiang University) for suggestions. We thank Dr. Yingui Cao (China University of Geosciences, Beijing) for kindly providing the boundary map of the Taihang Mountain.

