## Supplemental Figures for "Evolutionary genomics of local adaptation, climate vulnerability, and linarin metabolism in *Opisthopappus longilobus*"

**Supplementary Figures**

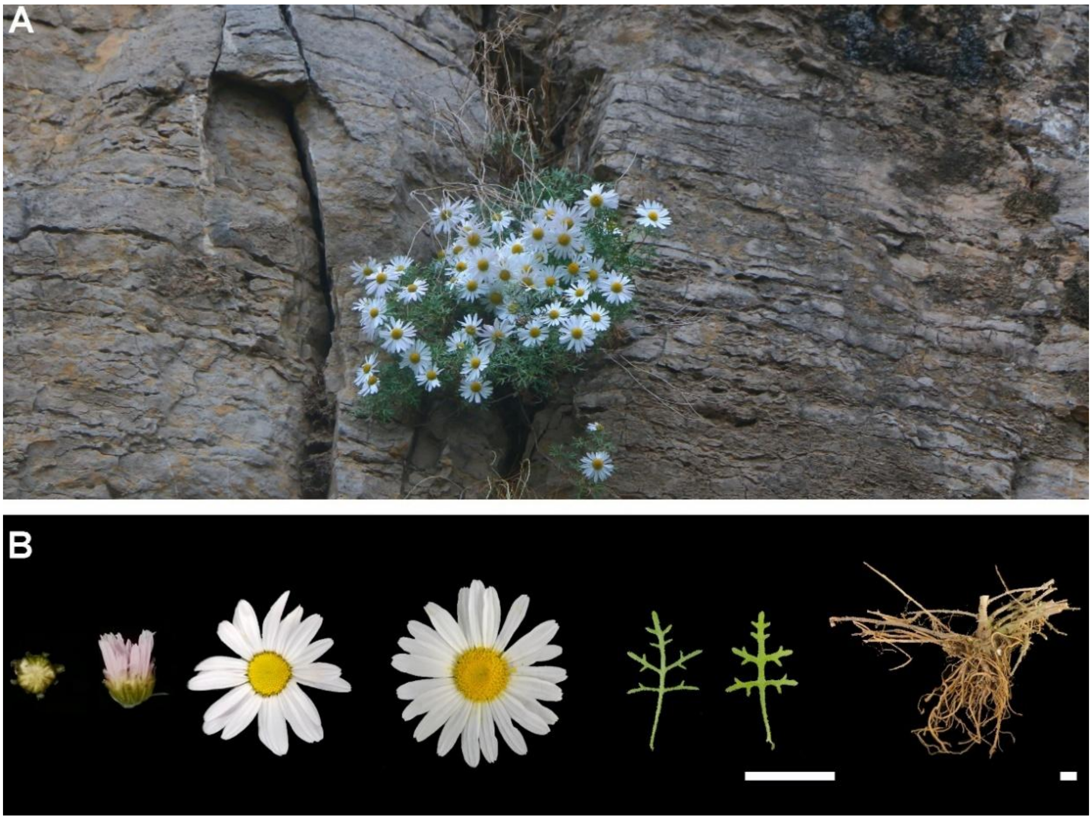


**Supplementary Figure 1. An Overview of the natural habitat (A) and phenotypic characteristics (B) of *O. longilobus*.** Buds, flowers and leaves share a common scale bar, whereas roots are shown with a separate one. Scale bar = 1 cm.

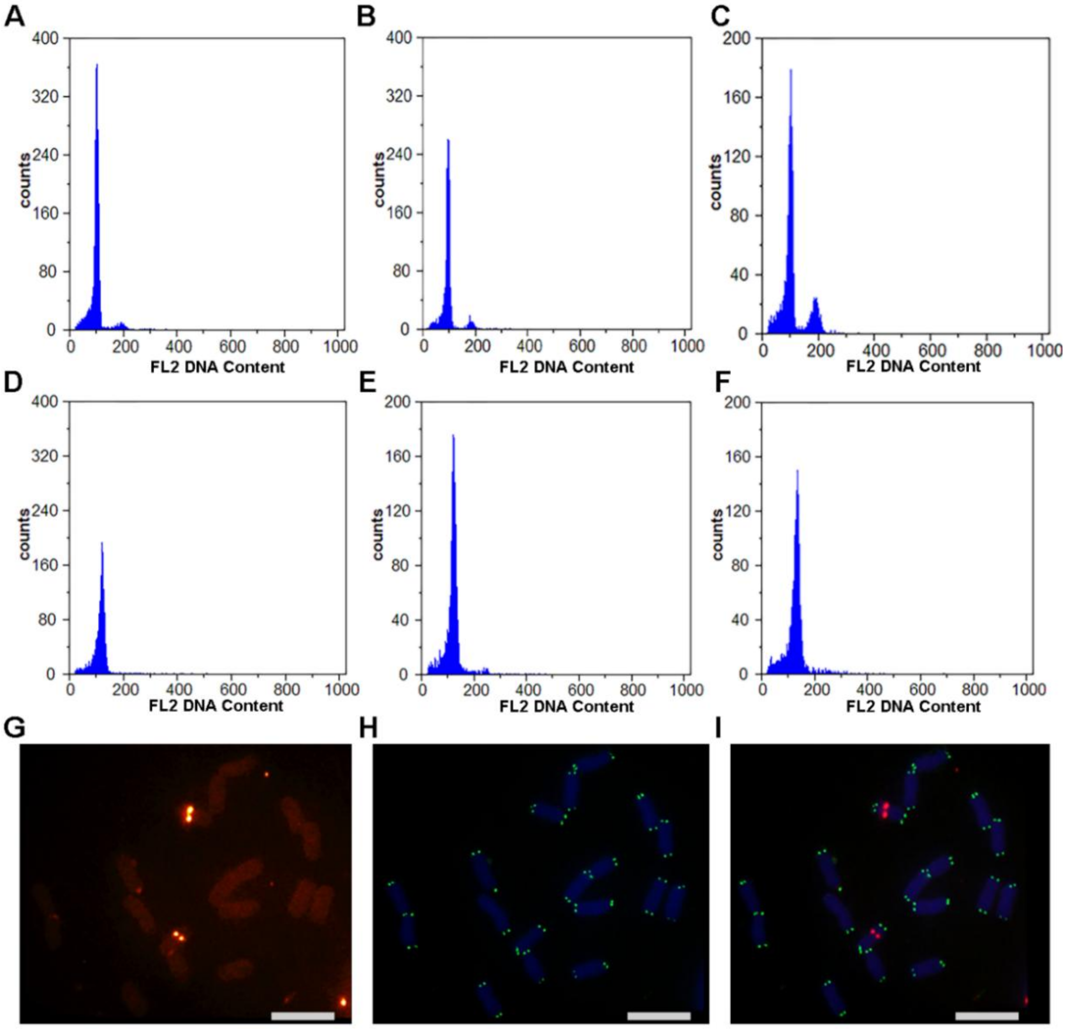


**Supplementary Figure 2. Flow cytometric genome size estimation and Oligo-FISH mapping of 5S rDNA and telomere signals for *O. longilobus*. A-C,** The diploid *Z. mays* with a genome size of 2655.3 Mb was used a reference in cell flow cytometry analyses. **D-F,** *O. longilobus* was estimated to be a diploid with a genome size of 3360.36 Mb. **g,** Visualization of 5S rDNA sites using 5S rDNA probes. **H,** Visualization of telomere sites using telomere probes. **I,** Merged image showing the chromosomal localization of telomere and 5S rDNA sites. Scale bars = 10 µm.

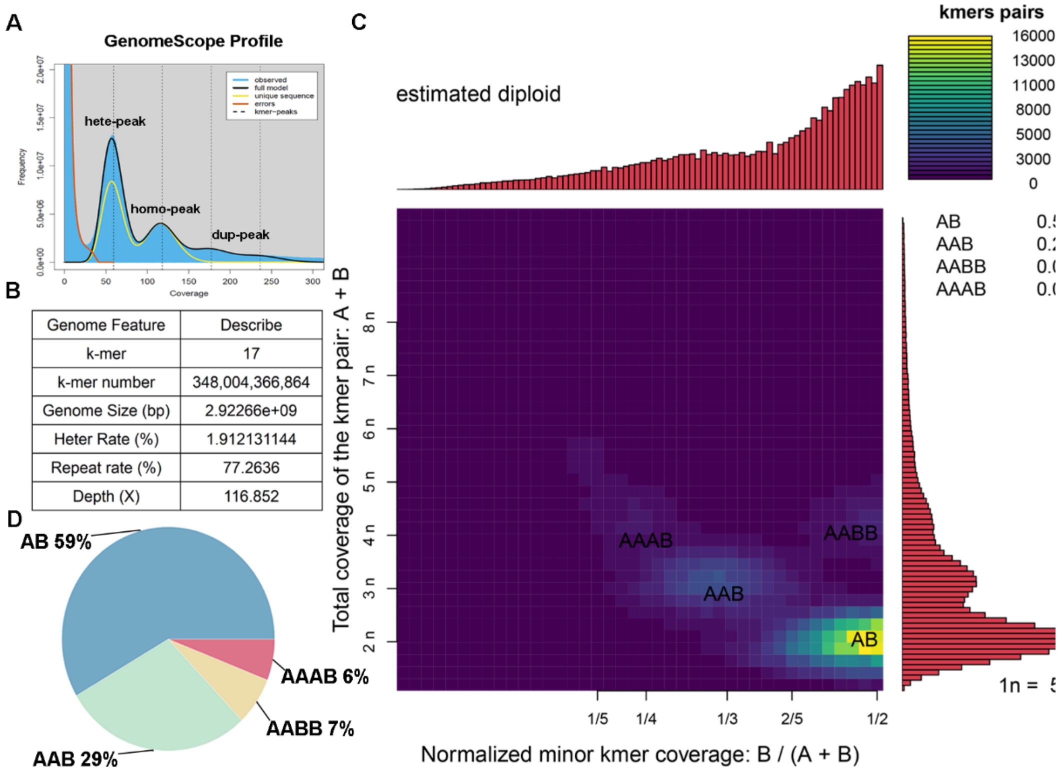


**Supplementary Figure 3. Smudgeplot analysis and *K*-mer depth distribution in the genome of *O. longilobus*. A,** Diploid GenomeScope plot. X axis shows *K*-mer = 17 depth. Y axis shows *K*-mer frequency. The genome size was measured as 2.92 Gb. **B,** Genomic features inferred using Jellyfish and GCE analyses. **C,** Diploid Smudgeplot. The Y-axis represents the total coverage of homologous *K*-mer pairs (CovA+CovB) from allelic or duplicated loci in the *O. longilobus* genome, and the X-axis shows the relative coverage of minor *K*-mers (CovB/(CovA+CovB)). The color bar indicates the number of *K*-mer pairs from distinct genomic structures (AB, AAB, AABB, and AAAB). **D,** The proportion of different type *K*-mer pairs in the *O. longilobus* genome.

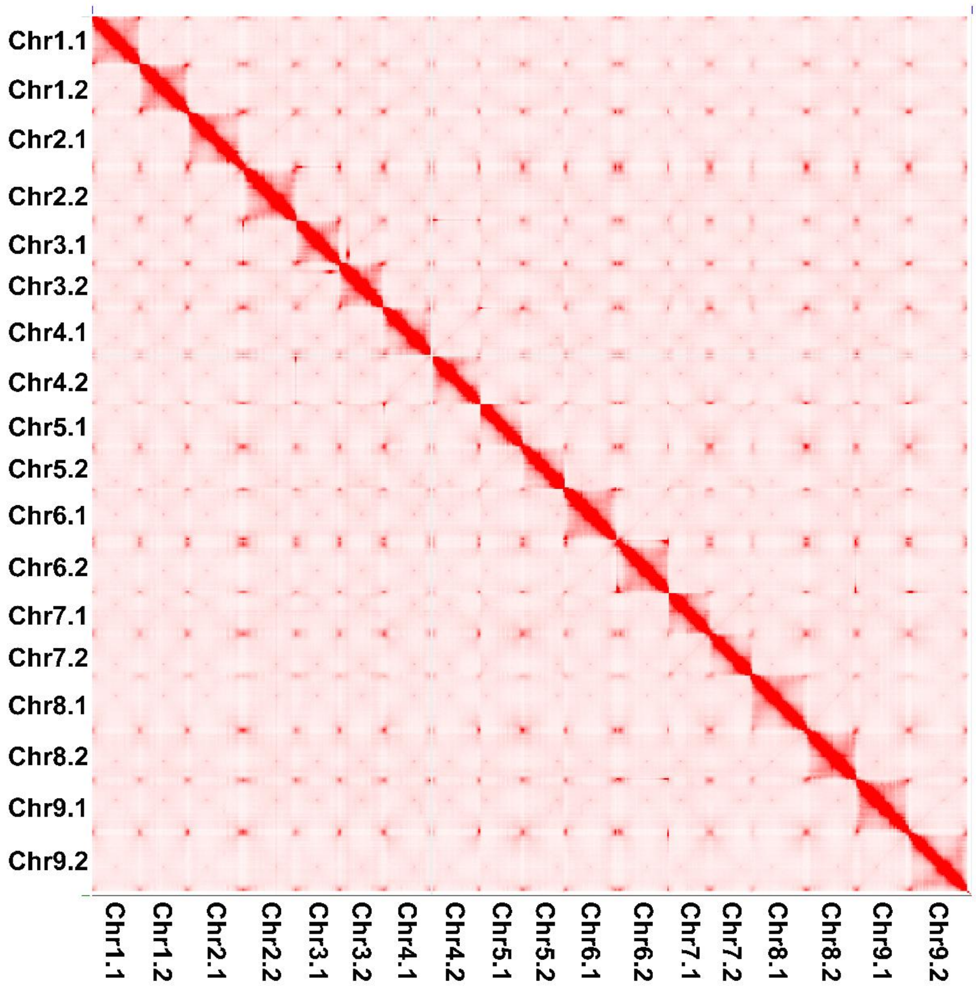


**Supplementary Figure 4. Hi-C interaction signals of *O. longilobus*.** The densely red areas represent stronger signals. There was a total of 2n = 18 chromosomes in the figure, and the chromosomes that show interacting signals (thin red lines) are considered homologous.

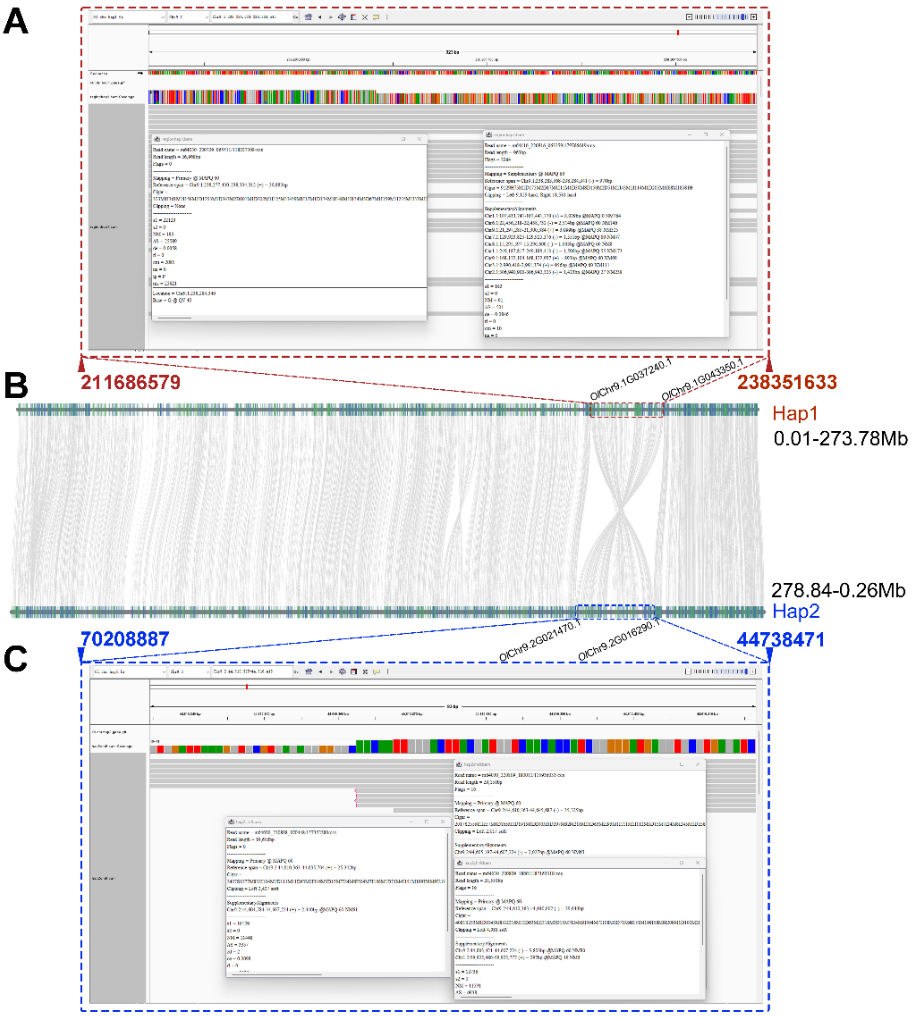


**Supplementary Figure 5. Analysis of an inversion identified on chromosome 9 of *O. longilobus*. A,** IGV visualization of the inversion region in haplotype 1 (Hap1) genome (Chr9.1:211,686,579-238,351,633). **B,** Collinearity analysis of homologous chromosomes using JCVI. **C,** IGV visualization of the inversion region in haplotype 2 (Hap2) genome (Chr9.2:44,738,471-70,208,887).

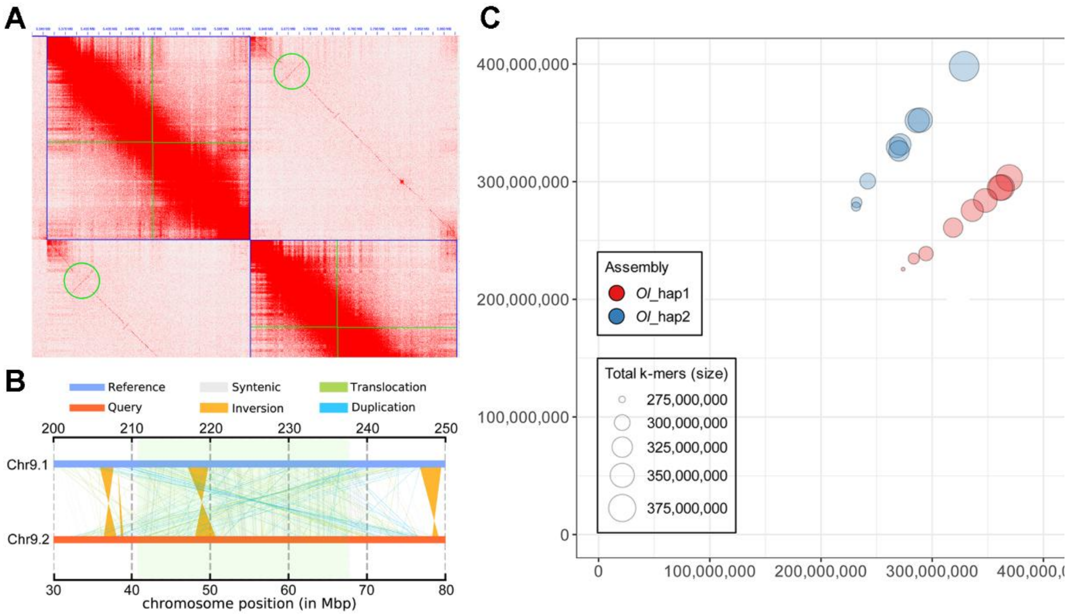


**Supplementary Figure 6. Inversion identified on chromosome 9 and haplotype-specific *K*-mer bubble plot in the *O. longilobus* genome. A,** Verification of the inversion detected between two haplotypes of *O. longilobus* by conflict signals of Hi-C interaction, which was marked by green circles. **B,** Summary of structural and sequence variations detected by SyRI. **C,** The bubble plot displays *K*-mers that are specific to either Hap1 or Hap2 of haplotype-resolved *O. longilobus* genome. Bubble size is proportional to the length of the corresponding contigs.

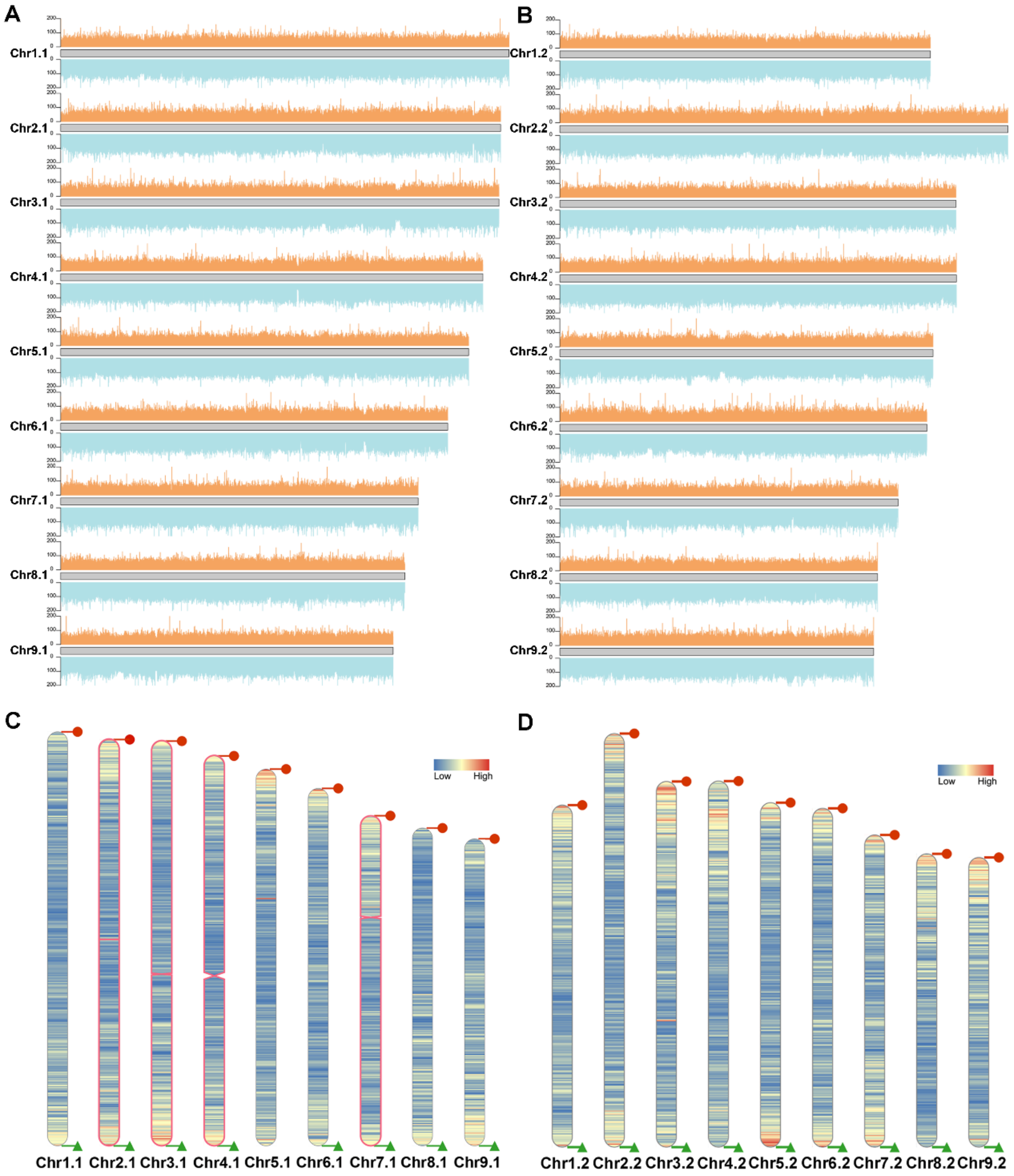


**Supplementary Figure 7. The landscape of read depth and telomere/centromere distribution in the *O. longilobus* haplotype-resolved genome. A, B,** The depth distribution plots for Hap1 and Hap2, respectively. The upper orange bars represent the genome coverage of PacBio long reads, while the lower blue bars correspond to the genome coverage of Illumina short reads. These profiles are used to assess assembly completeness and heterozygosity. **C, D,** Telomeric and centromeric feature mapping in Hap1 and Hap2, respectively.

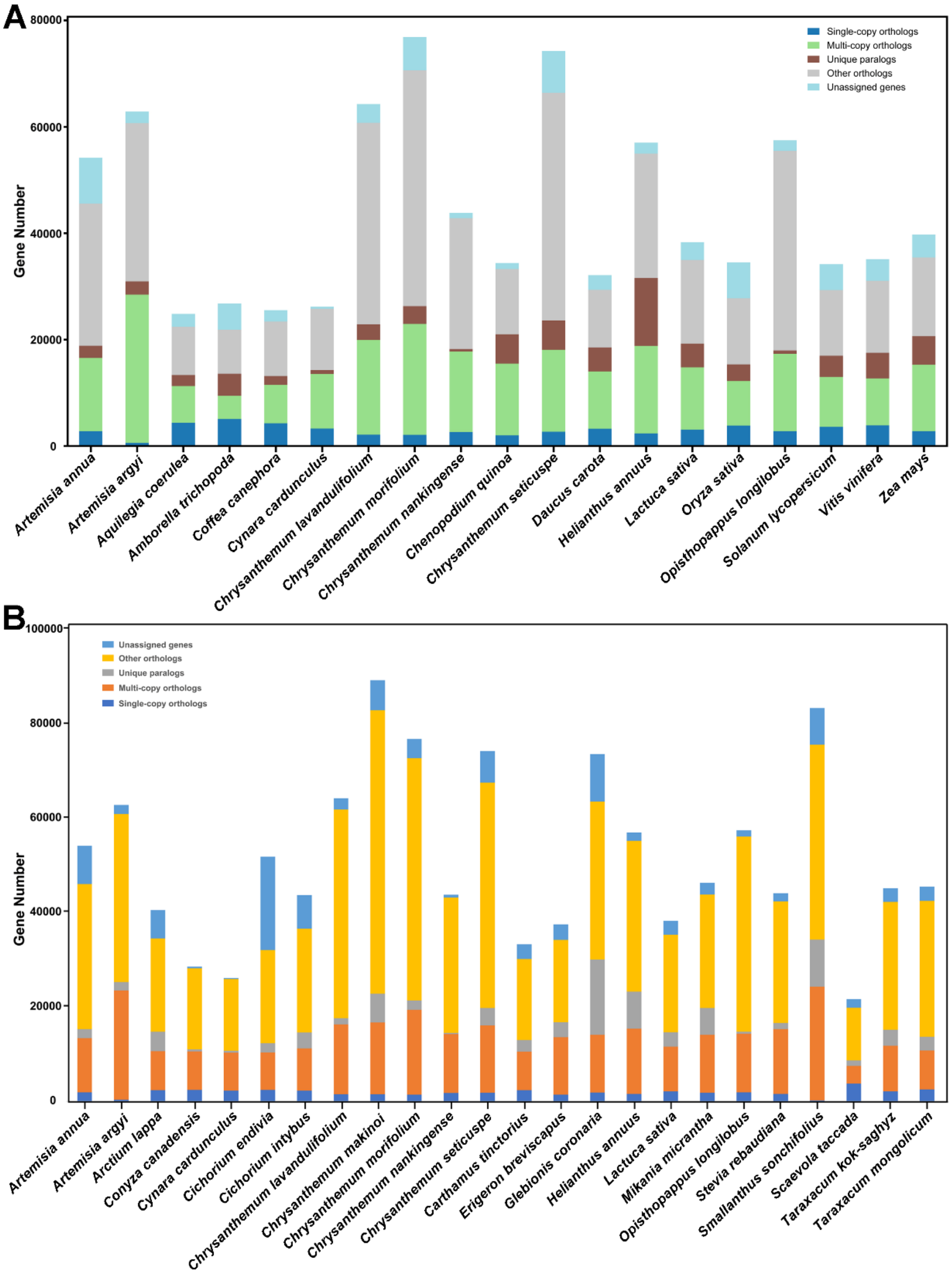


**Supplementary Figure 8. Gene family statistics used in comparative genomics analysis. A,** Gene family clustering across *O. longilobus* and 18 representative plant species**. B,** Gene family clustering statistics within 23 Asteraceae species and the outgroup *Scaevola taccada*. Source data are provided in Supplementary Tables 23 and 24.

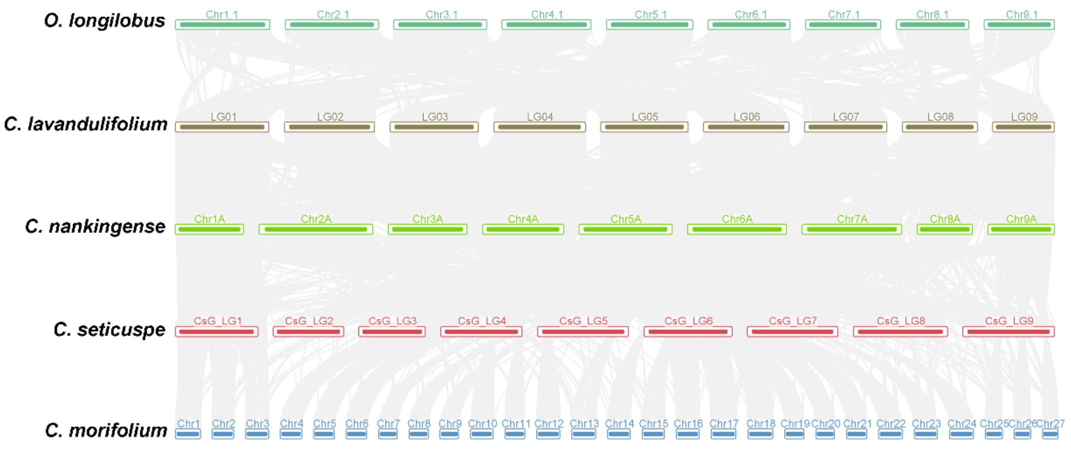


**Supplementary Figure 9. Syntenic diagrams between *O. longilobus* and four *Chrysanthemum* species.** Syntenic blocks are marked using gray lines.

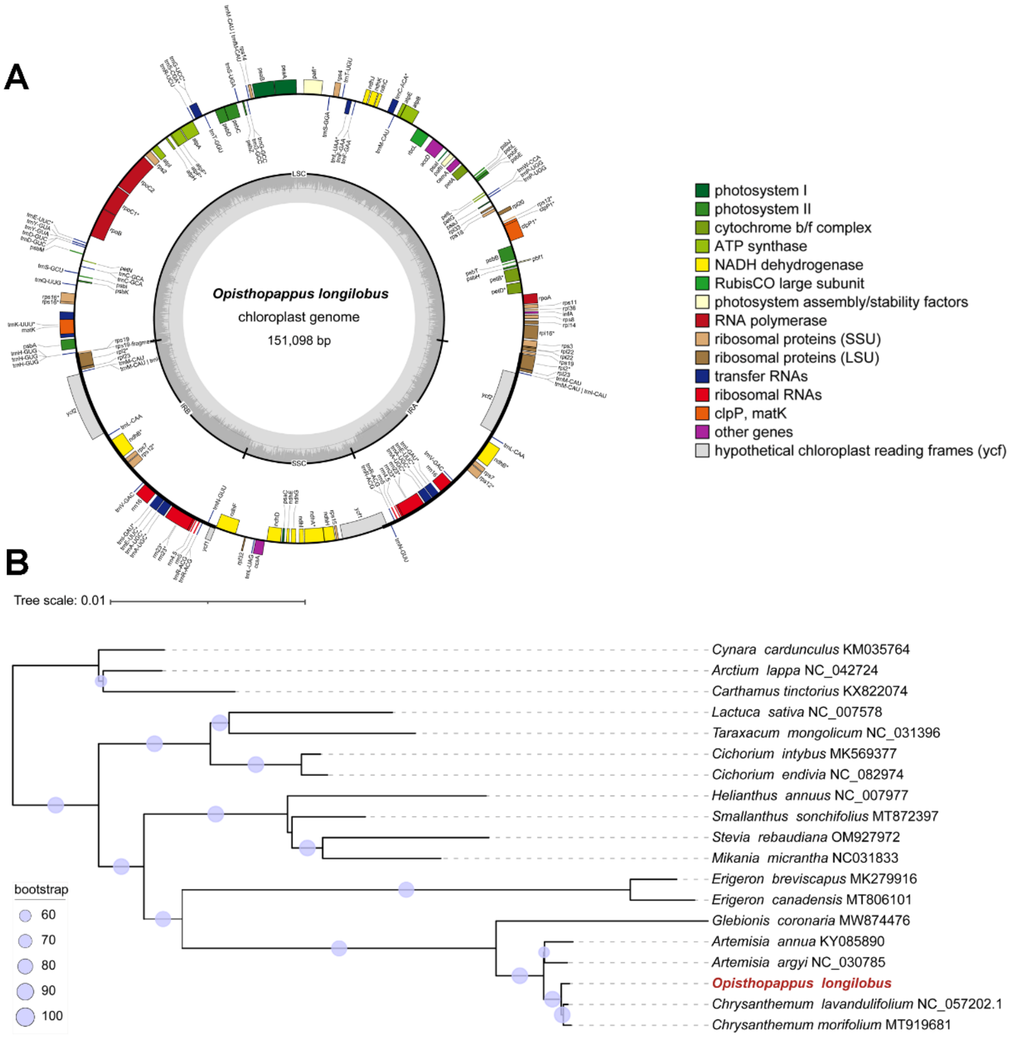


**Supplementary Figure 10. Analysis of chloroplast genome structure and phylogenetic relationships among Asteraceae species. A,** Map of the chloroplast genome of *O. longilobus*. Genes lying outside the outer circle are transcribed in a counter-clockwise direction, while genes inside this circle are transcribed in a clockwise direction. The colored bars indicate known protein-coding genes, transfer RNA genes, and ribosomal RNA genes. The area shaded with dark grey dashed lines in the inner circle denotes GC content, and the light grey area indicates AT content. LSC, large single-copy, SSC, small single-copy. IR, inverted repeat. **B,** Phylogenetic relationships of Asteraceae based on complete chloroplast genomes. Maximum likelihood (ML) phylogenetic tree was reconstructed from complete chloroplast genome sequences of *O. longilobus* and 18 other representative Asteraceae species, with NCBI accession numbers given after each species name. The purple circles above the branches indicate the ML bootstrap support values. The target species *O. longilobus* is highlighted in red.

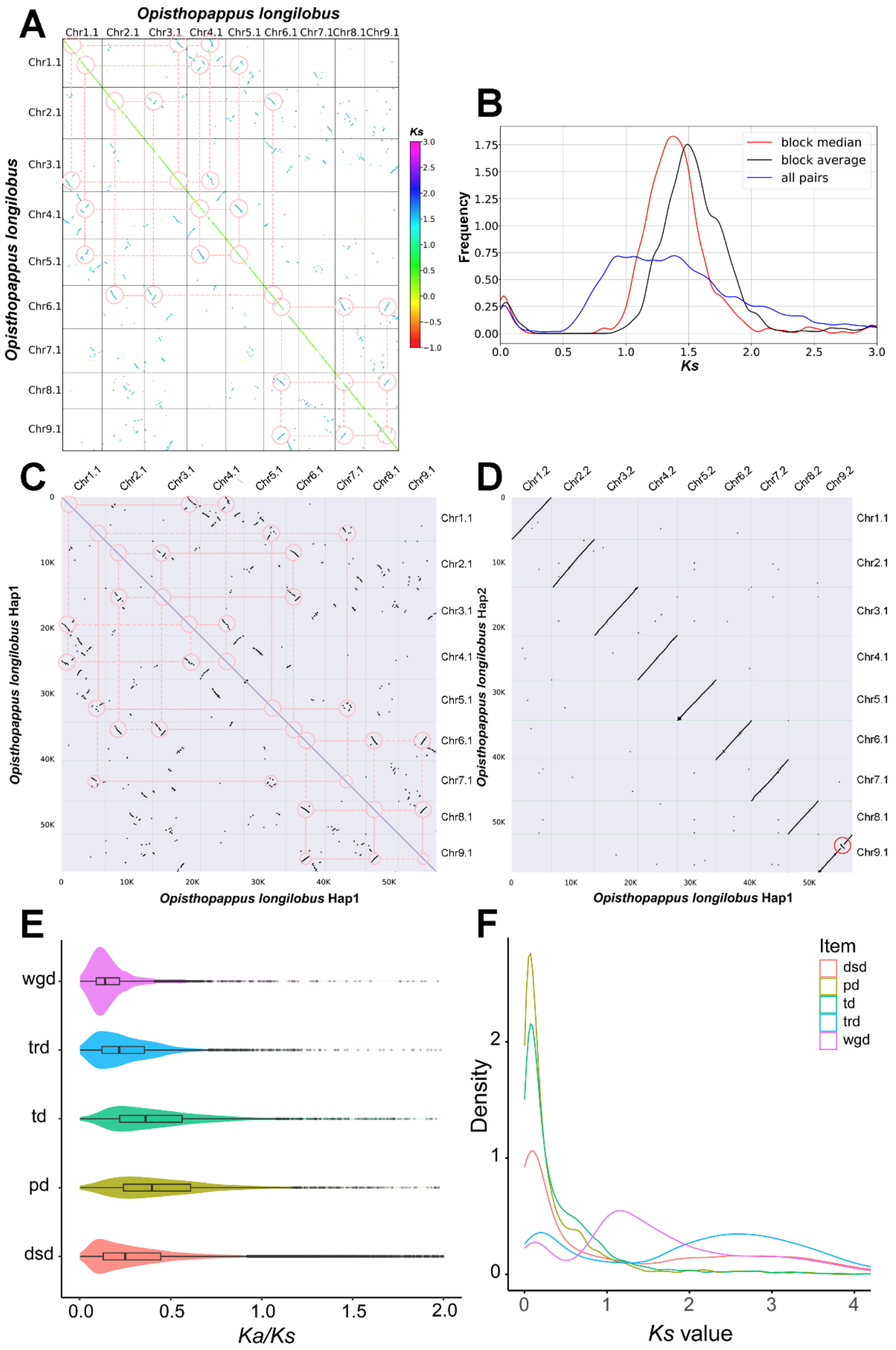


**Supplementary Figure 11. Genome-wide *Ks* heatmap, synteny relationships, and duplication-mode distributions in *O. longilobus*. A,** *Ks* thermogram of *O. longilobus* after removal of tandem repeats. The graph reveals traces of WGT in *O. longilobus*, indicating that it has undergone whole-genome triplication. **B,** Peak plot of *Ks* in *O. longilobus*. **C,** Synteny plot of *O. longilobus* Hap1 genome assembly. **D,** Synteny plot between Hap1 and Hap2. **E,** *Ka/Ks* distribution of different duplication types. **F,** *Ks* distribution of different duplication types.

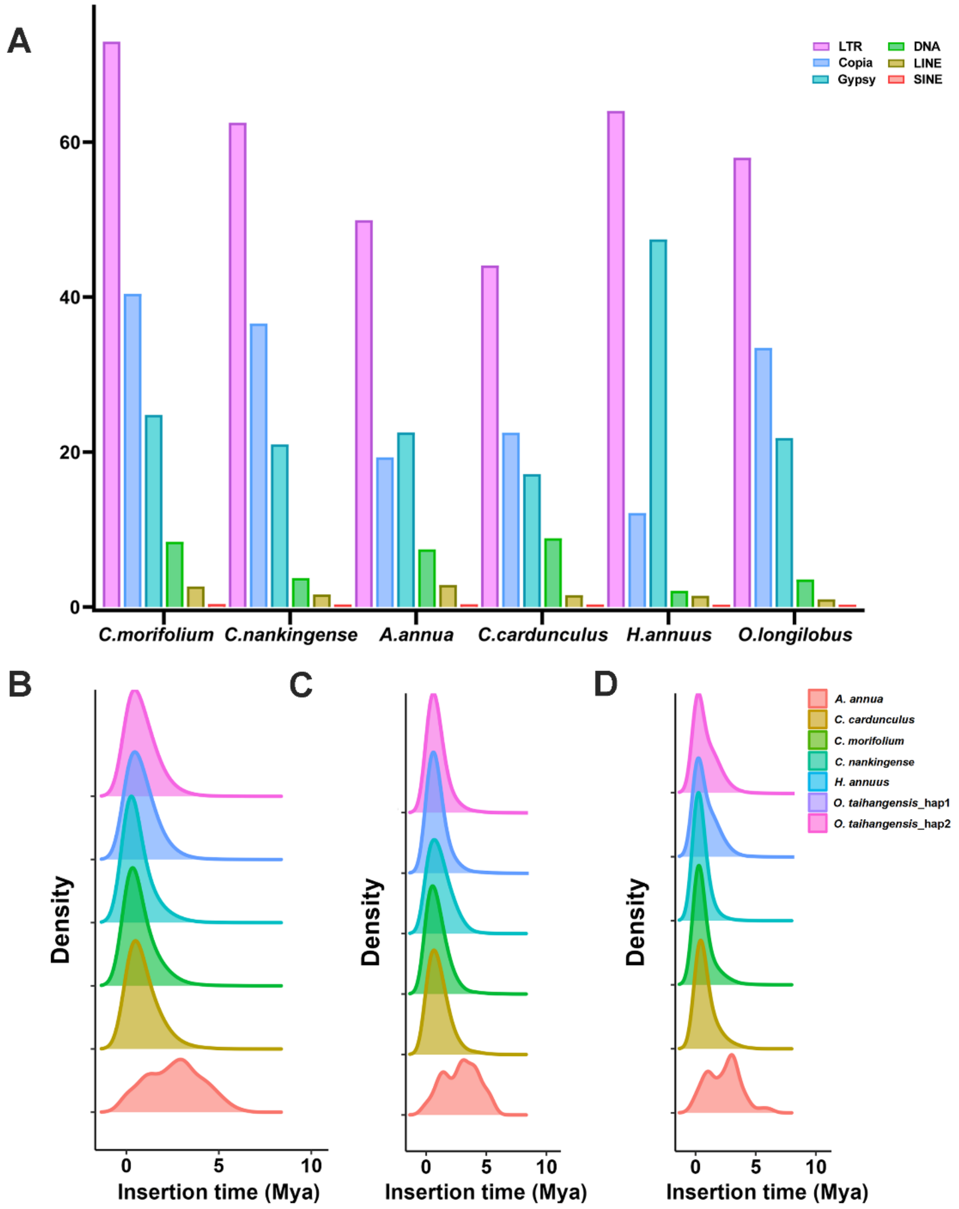


**Supplementary Figure 12. Comparative analysis of repetitive elements and LTR insertion histories across Asteraceae species. A,** Comparison of repetitive elements among *O. longilobus*, *C. morifolium*, *C. nankingense*, *A. annua*, *H. annuus*, and *C. cardunculus*. **B-D,** Distributions of complete LTR (b), *Copia* (c), and *Gypsy* (d) insertion times across different Asteraceae species.

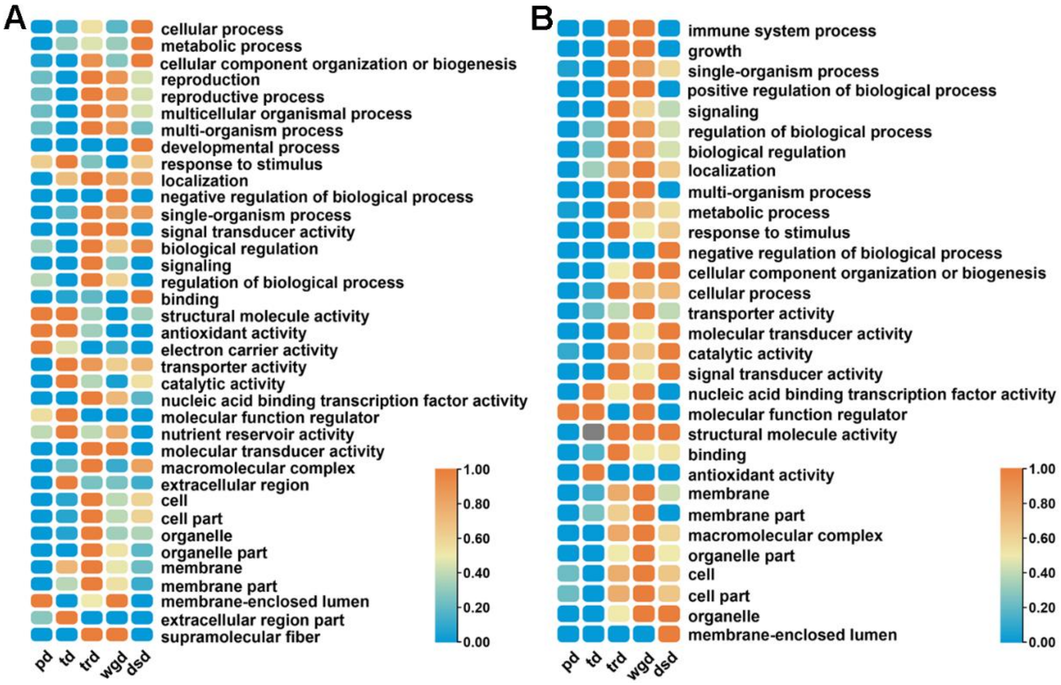


**Supplementary Figure 13. GO category enrichment analysis on the EPGs (A) and PAML (B) of five types of gene duplications.**

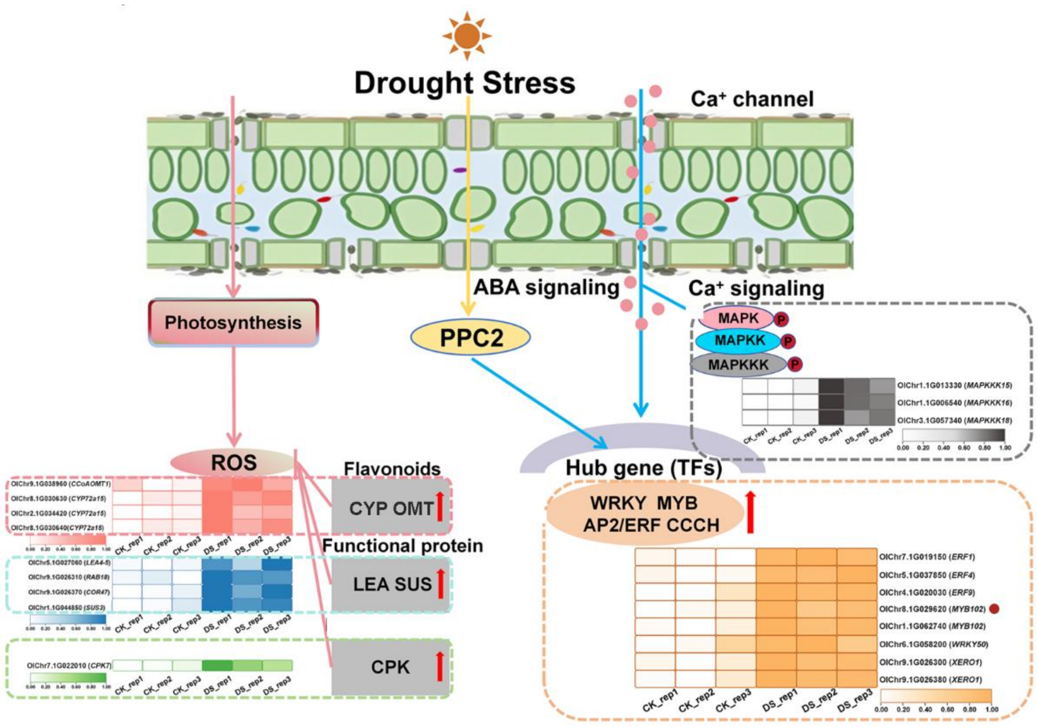


**Supplementary Figure 14. A hypothetical model of the drought tolerance mechanism for *O. longilobus.*** Source data are provided in Supplementary Tables 18, 26, and 29.

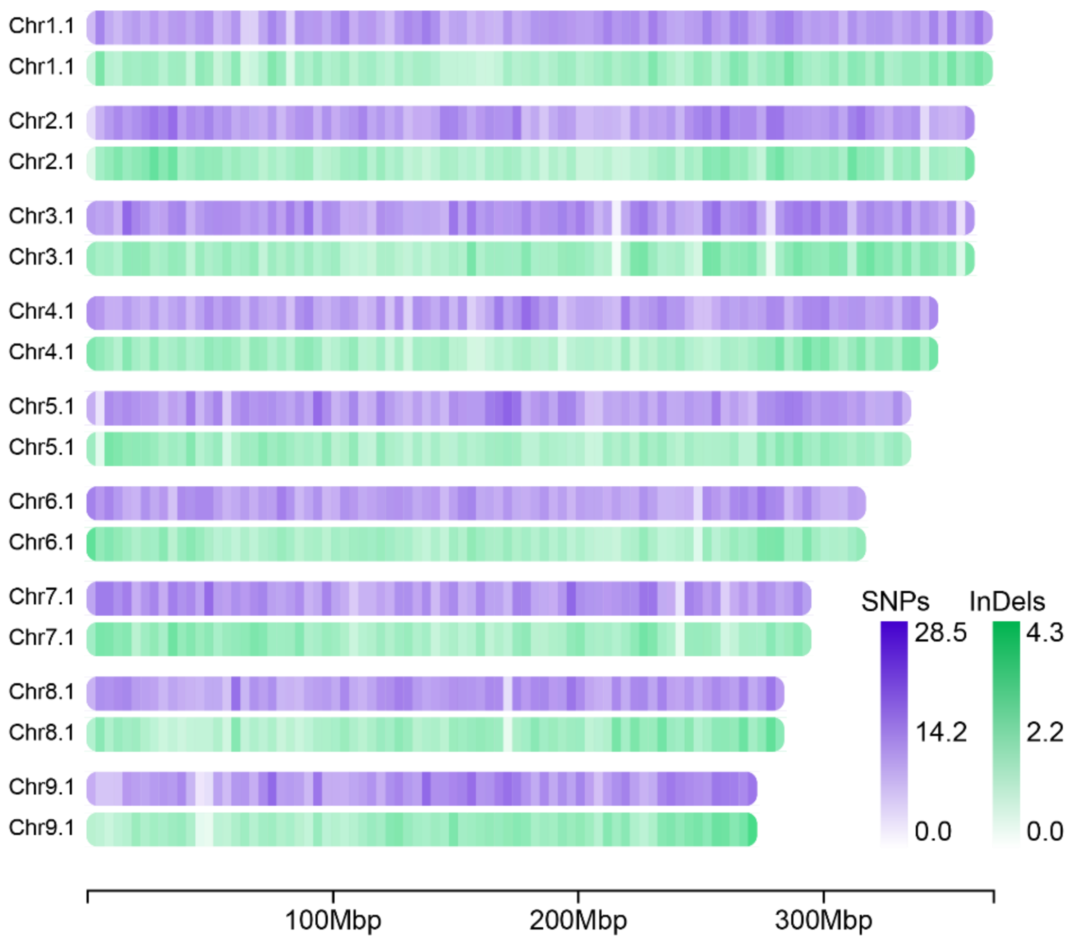


**Supplementary Figure 15. The density distribution of SNPs and InDels across 9 *O. longilobus* pseudochromosomes under 1 Mb windows.**

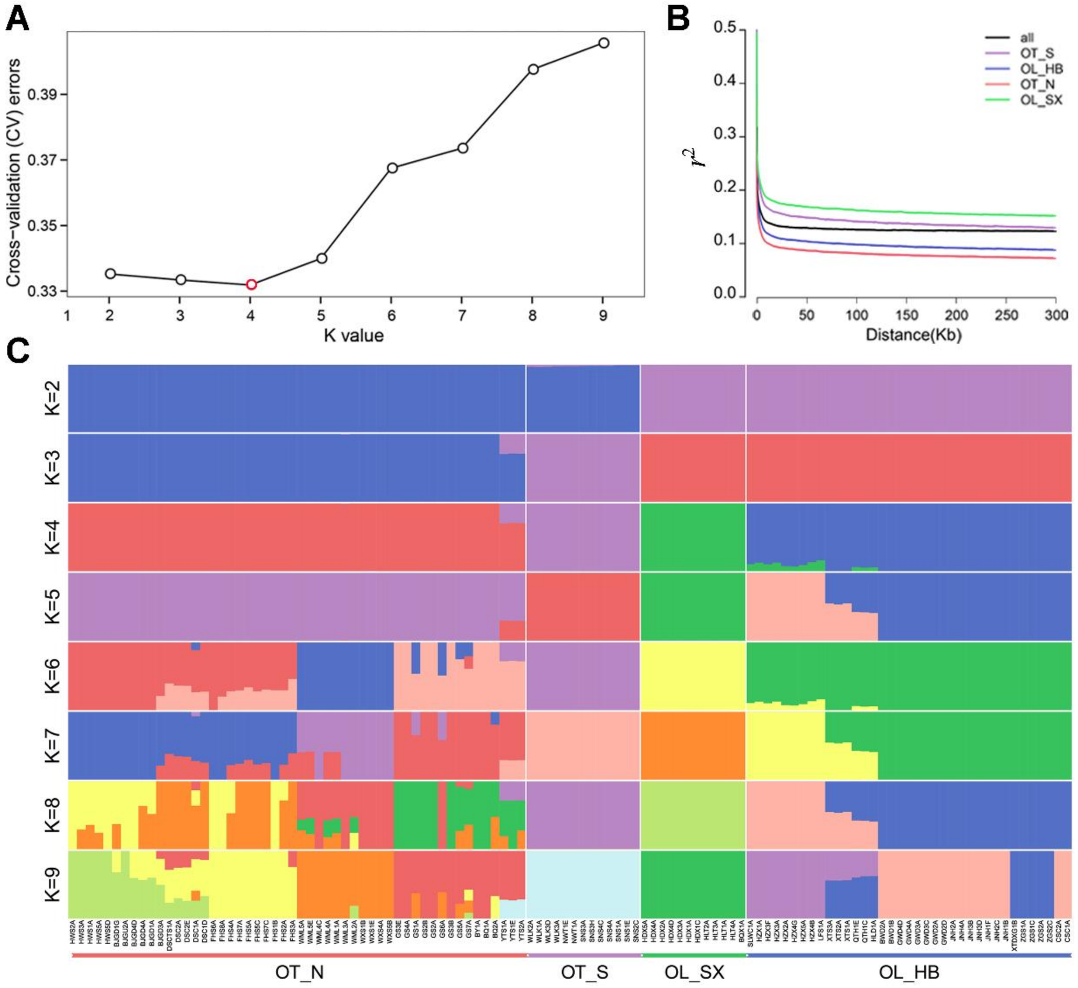


**Supplementary Figure 16. Population structure and LD decay analyses for *Opisthopappus* species. A,** Cross-validation (*CV*) error curve with assumed number of clusters K ranging from 2 to 9. The optimal K = 4 is indicated by a red point. **B,** Individual ancestry coefficients from K = 2 to K = 9 of 115 *Opisthopappus* accessions inferred by ADMIXTURE. Each individual is denoted by a vertical bar, and the Y-axis quantifies the proportion of the individual's genome derived from inferred ancestral lineages. **C,** Linkage disequilibrium (LD) decay patterns in four sub-populations and whole population. The X-axis indicates physical distances between two SNPs. The Y-axis is $r^2$ (squared correlation coefficient) that used to measure LD level. Colors in each population are based on ADMIXTURE analyses when K = 4.

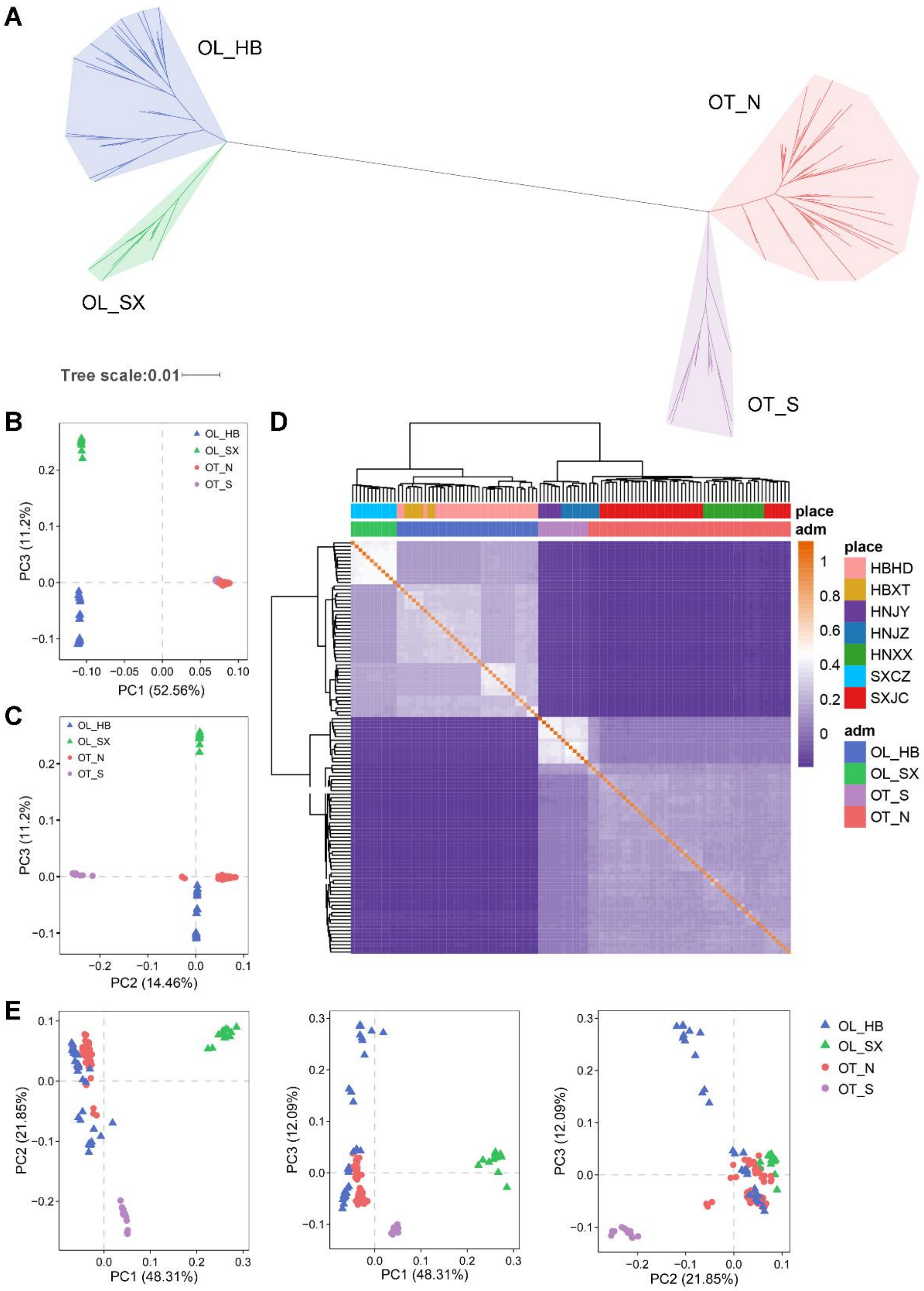


**Supplementary Figure 17. Phylogenetic relationships of 115 resequenced *Opisthopappus* accessions. A,** A unrooted phylogenetic tree of 115 accessions inferred from whole-genome SNPs. **B, C,** PCA plots based on 3,808,756 LD-pruned SNPs. **D,** Heatmap of kinship relationships among resequenced individuals. **E,** PCA analysis based on 4,620 adaptive variants. In **B, C** and **E,** individuals with different color denotes different groups that determined by ADMIXTURE analysis when K = 4. Triangles and dots represent *O. longilobus* and *O. taihangensis* accessions, respectively.

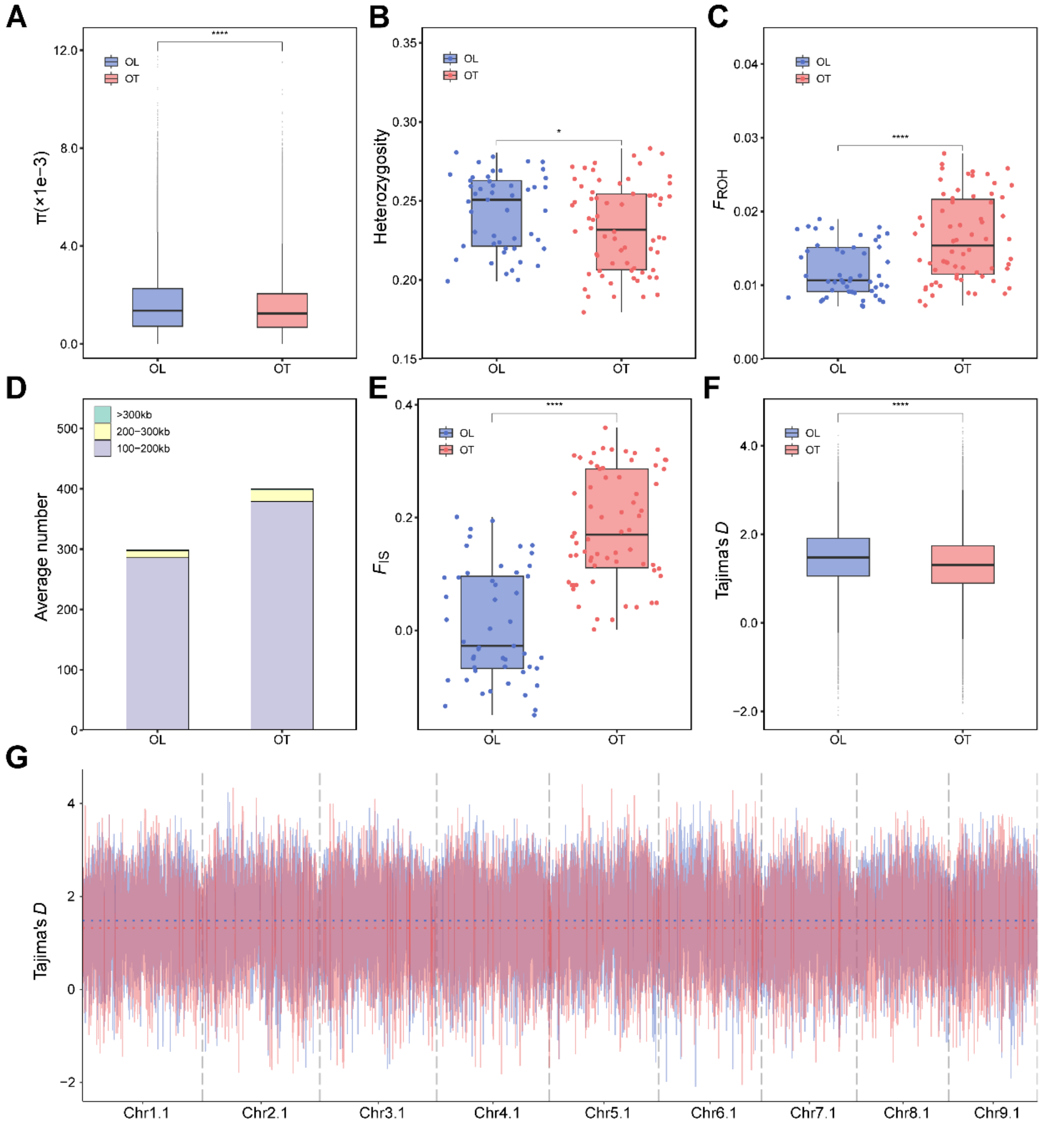


**Supplementary Figure 18. Comparisons of population genetic statistics between *O. longilobus* and *O. taihangensis*. A-C, E** and **F,** The distributions of nucleotide diversity ($\pi$), individual heterozygosity, the genome proportion of long runs of homozygosity ($F_{ROH}$, > 100 kb), inbreeding coefficient ($F_{IS}$), Tajima's *D* values, respectively. The central line, edges and whiskers for each box correspond to the median, the 25th and 75th quartiles, 1.5 times the inter-quartile range (IQR), respectively. *, **** indicate significance at the 5% and 0.01% level, respectively, as calculated by Wilcox test. **D,** Proportion of the genome with ROH longer than 100 kb, 200 kb and 300 kb. **G,** Comparison of Tajima's *D* values between *O. longilobus* (blue line) and *O. taihangensis* (red line) across 9 pseudochromosomes.

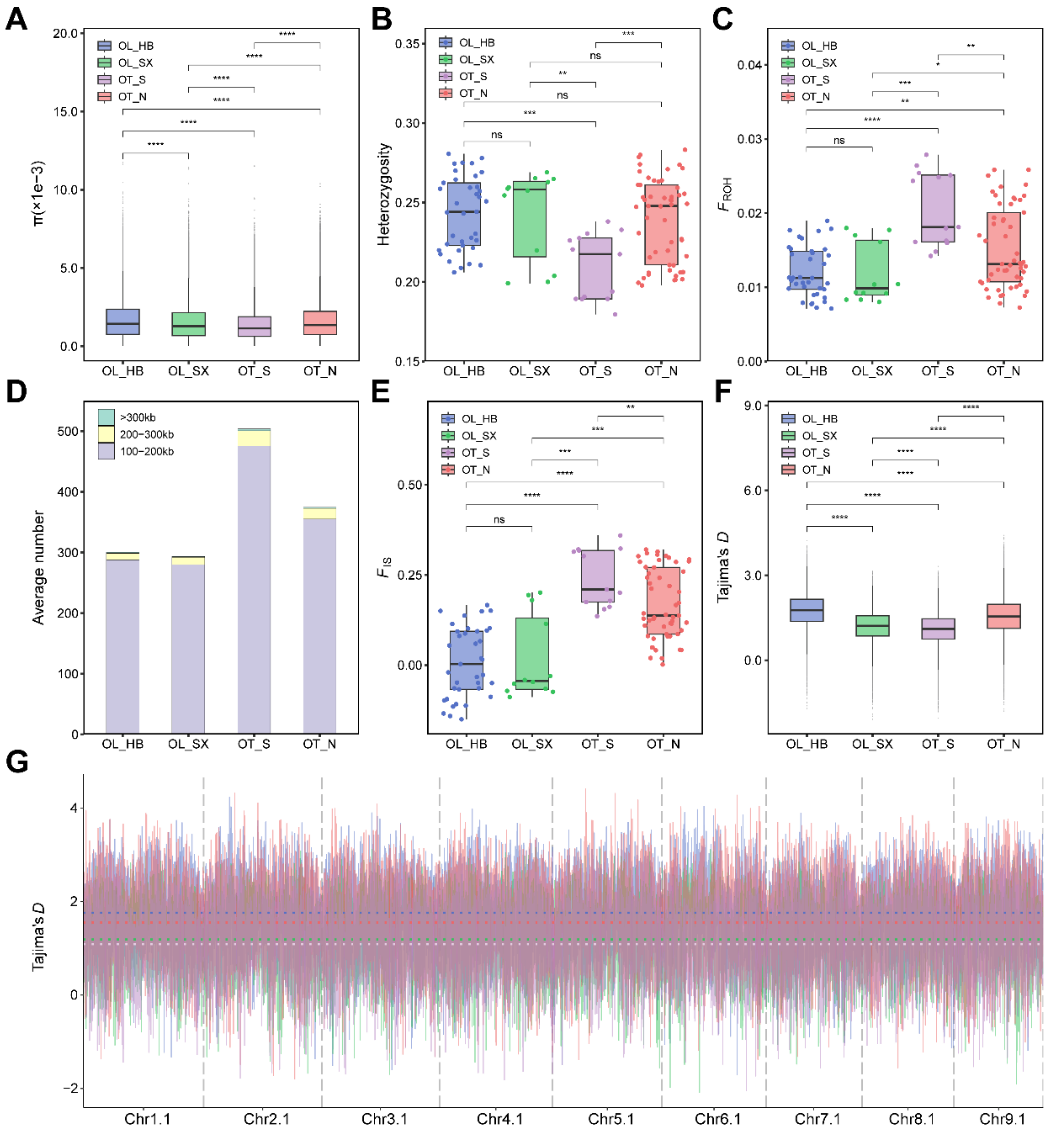


**Supplementary Figure 19. Comparisons of population genetic statistics among four intraspecific lineages of *Opisthopappus*. A-C, E** and **F,** The distributions of nucleotide diversity ($\pi$), individual heterozygosity, the genome proportion of long runs of homozygosity ($F_{\mathrm{ROH}}$, > 100 kb), inbreeding coefficient ($F_{\mathrm{IS}}$), Tajima's $D$ values, respectively. The central line, edges and whiskers for each box correspond to the median, the 25th and 75th quartiles, 1.5 times the inter-quartile range (IQR), respectively. ns, not significance. *, **, *** and **** indicate significance at the 5%, 1%, 0.1%, and 0.01% level, respectively, as calculated by Wilcox test. **D,** Proportion of the genome with ROH longer than 100 kb, 200 kb and 300 kb. **G,** Comparison of Tajima's $D$ values among four intraspecific lineages across 9 pseudochromosomes. Blue, green, red, and purple lines represent the group of OL_HB, OL_SX, OT_N, and OT_S, respectively.

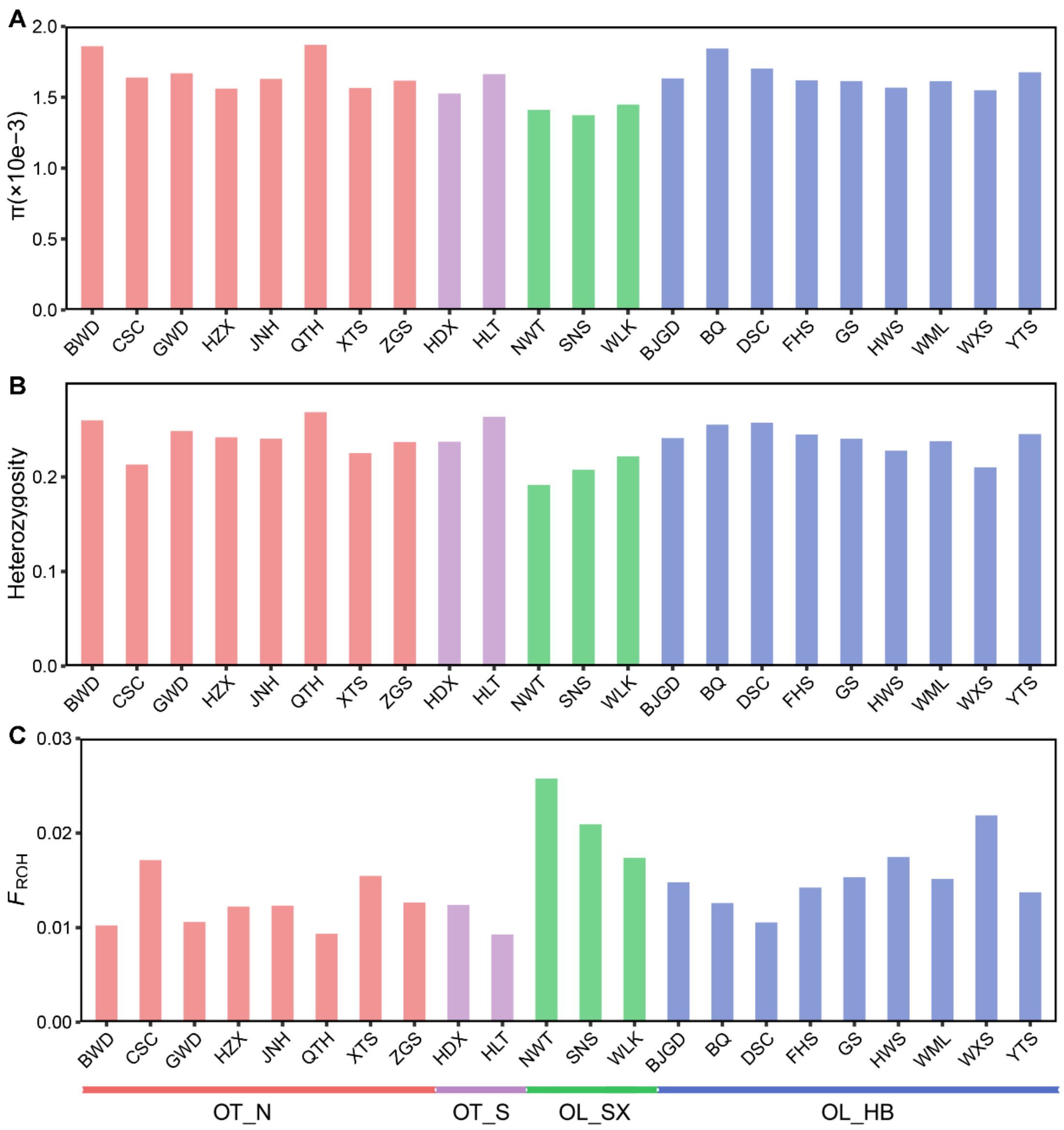


**Supplementary Figure 20. Comparisons of population genetic statistics among 22 populations. A-C,** The distributions of nucleotide diversity ($\pi$), individual heterozygosity, the genome proportion of long runs of homozygosity ($F_{\mathrm{ROH}}$ > 100 kb), respectively. The population abbreviations are described in Supplementary Table 31.

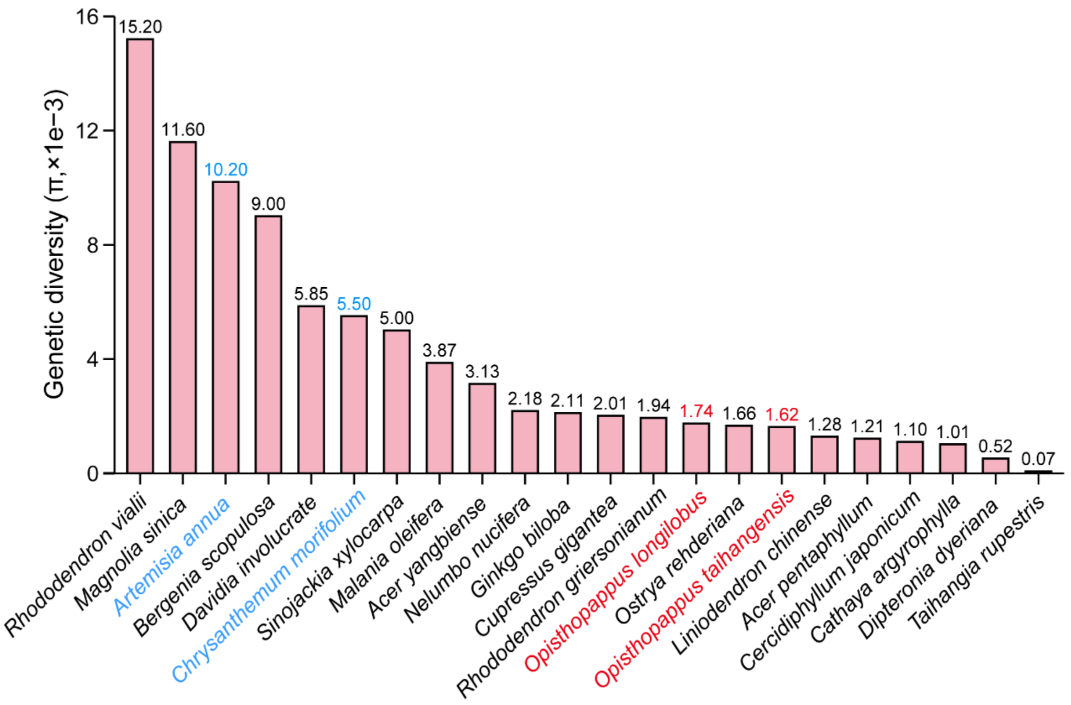


**Supplementary Figure 21. The nucleotide diversity ($\pi$) of 20 endangered species and 2 non-endangered Asteraceae species.** *O. longilobus* and *O. taihangensis* are marked by red fonts. Two representative Asteraceae species with broad distribution, *Chrysanthemum morifolium* and *Artemisia annua*, are highlighted in blue.

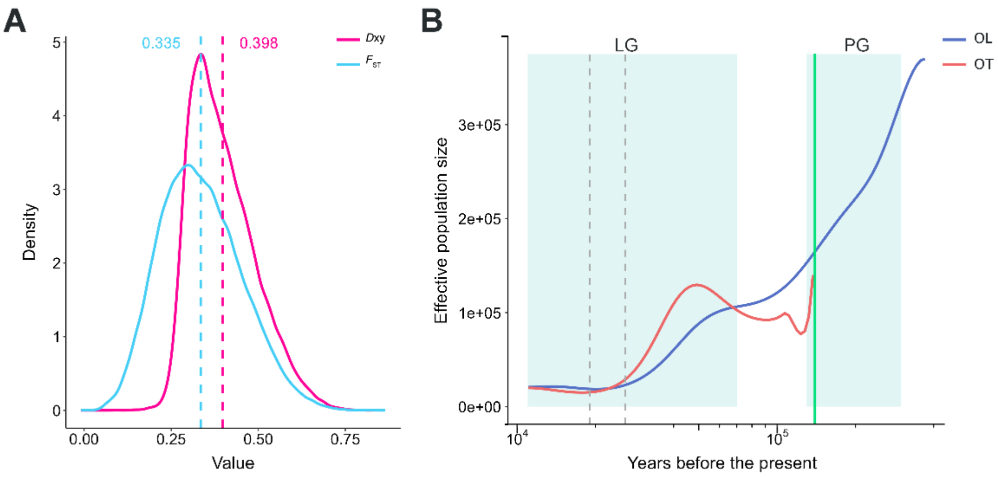


**Supplementary Figure 22. The genetic differentiation of *O. longilobus* and *O. taihangensis*. A,** The density distribution of fixation index ($F_{ST}$) and nucleotide divergence ($D_{XY}$) between *O. longilobus* and *O. taihangensis* over 100 kb windows across the genome. The dashed lines indicate the average estimates. **B,** Demographic changes on recent time scales established for *O. longilobus* and *O. taihangensis* using split command of SMC++ (g = 1, $\mu$ = 8.25e-9). Two periods, the last glacial (LG, 11-70 kya) and penultimate glaciation (PG, 130-300 kya), are shaded in light-green. The last glacial maximum (LGM) is indicated by two gray dashed vertical lines (LGM, 19-26.5 kya). The split time of *O. longilobus* and *O. taihangensis* is shown with a green dashed vertical line (x = 139.5 kya).

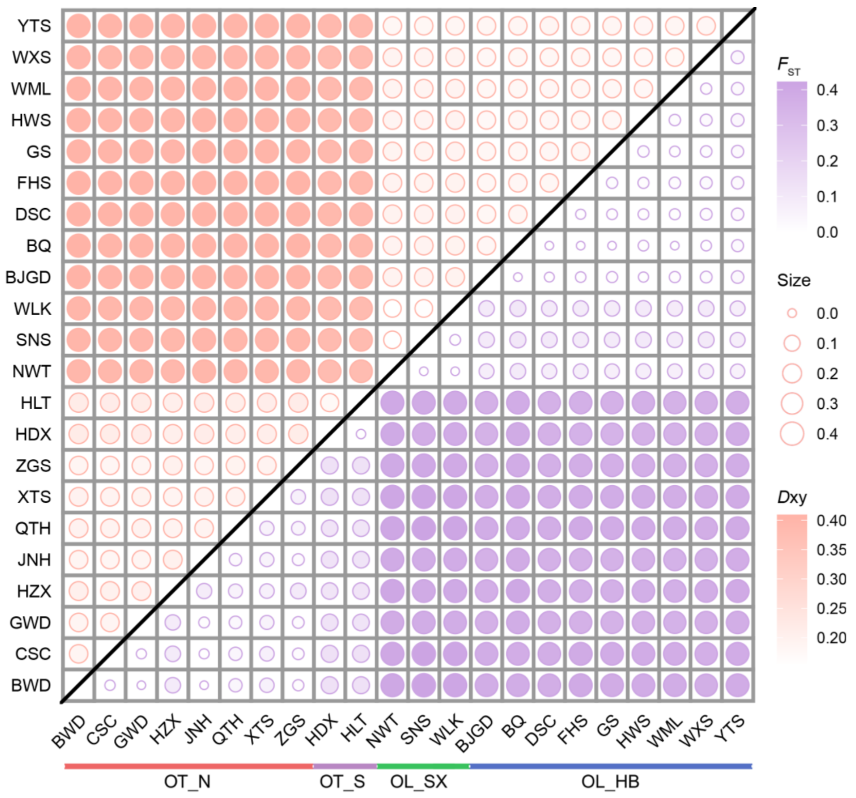


**Supplementary Figure 23. The heatmap of fixation index ($F_{ST}$, below diagonal) and nucleotide divergence ($D_{XY}$, above diagonal) among 22 *Opisthopappus* populations.** Source data are provided in Supplementary Table 35.

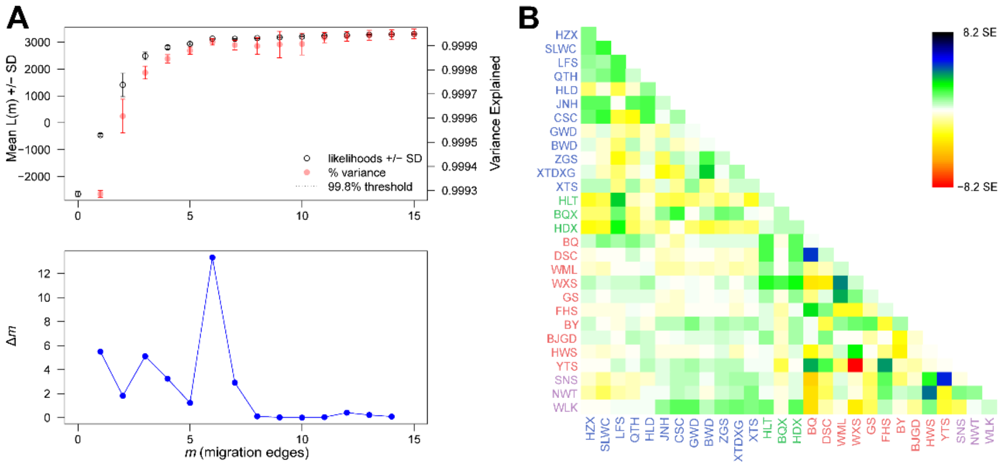


**Supplementary Figure 24. Infer the most possible number of migration edges among different *Opisthopappus* populations. A,** The output produced by OptM. The upper panel represents the mean ± standard deviation (*SD*) for the composite likelihood L($m$) (left axis) and proportion of variance explained (right axis). The lower panel shows the second-order rate of change ($\Delta m$) across values of $m$ = 0-15, indicting the optimal number of migrations is $m$ = 6. **B,** The residual fit from the maximum likelihood tree with six migrations inferred by TreeMix. Residuals above zero suggest the pairwise populations might undergone candidate admixture events.

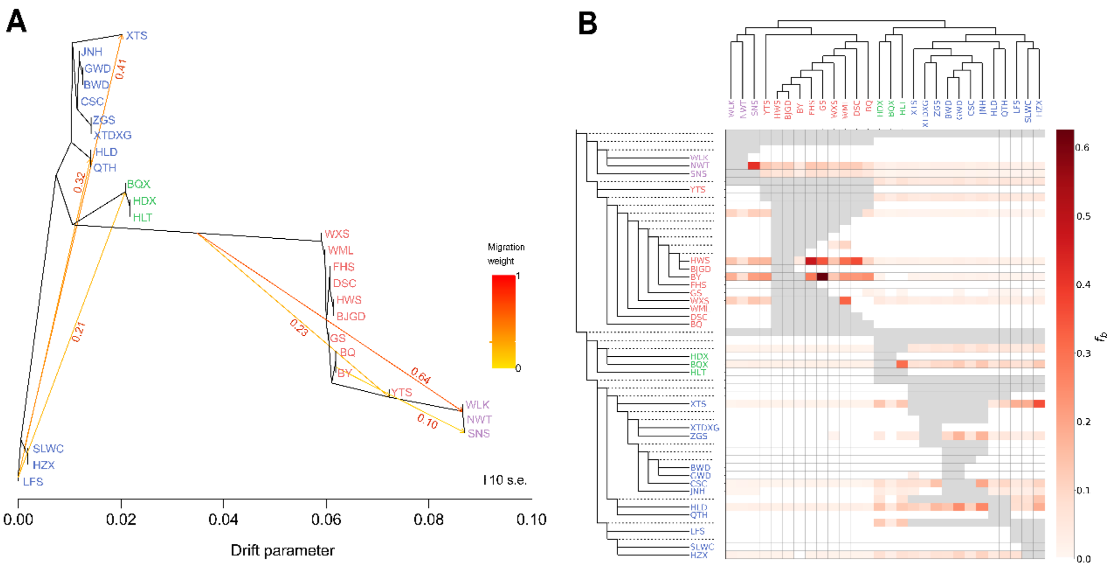

**Supplementary Figure 25. Signatures of gene flow and introgression in *Opisthopappus.*** **A,** Maximum-likelihood phylogenetic tree inferred by TreeMix with six migration events. Arrows show the direction of gene flow. Numbers in red indicate the migration weight, also illustrated by the color of the arrows (scale at bottom left). Scale bars represent a 10-fold average standard error (*SE*) of the accessions in the sample covariance matrix. **B,** Heatmap for pairwise *f*-branch statistics ($f_b$) tested by *D*suite. The colour scale at right indicates $f_b$ values of alleles excess sharing between the branch tree on the Y-axis and the population on the X-axis. Cells in grey indicate comparisons that cannot be made. See Supplementary Table 37 for detailed source data.

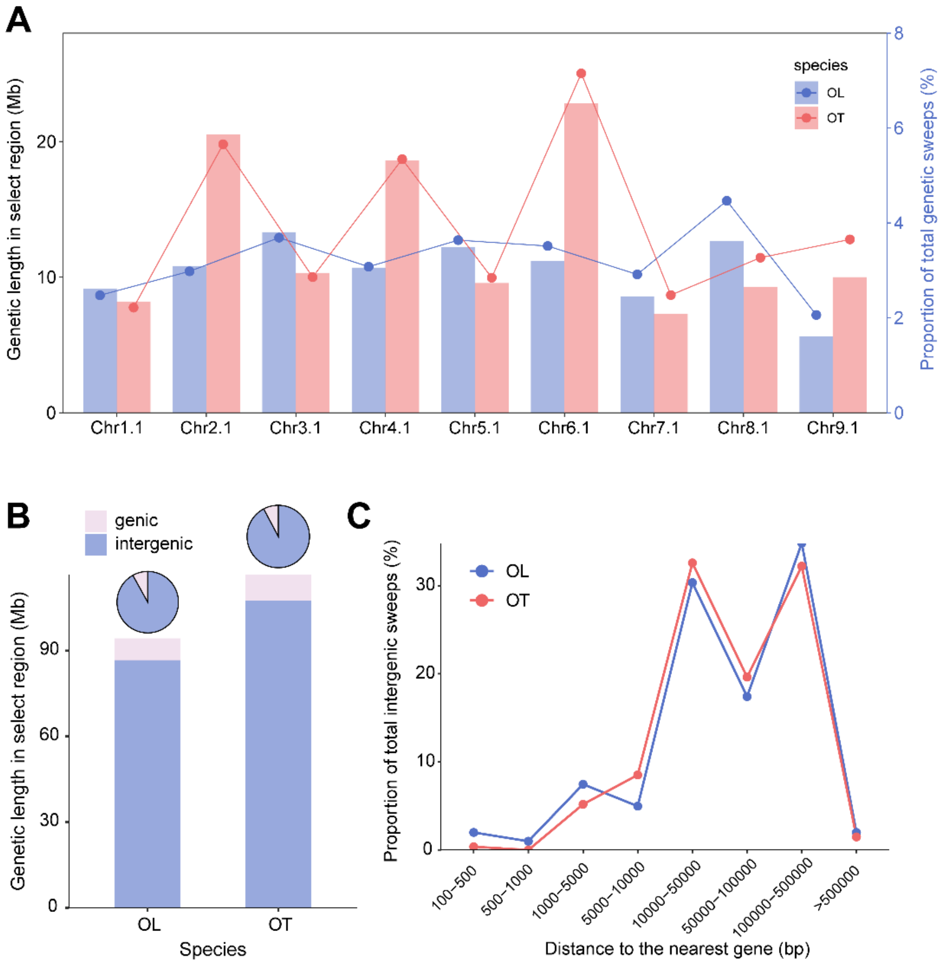


**Supplementary Figure 26. Distribution patterns of selective sweeps for *O. longilobus* and *O. taihangensis*. A,** Distribution of the selective sweeps length (left axis) and proportion (right axis) across 9 pseudochromosomes. **B,** The distribution ratio of selective sweeps located in genic and intergenic regions in two compared groups. **C,** Distribution of the distances from intergenic selective sweeps to the nearest gene.

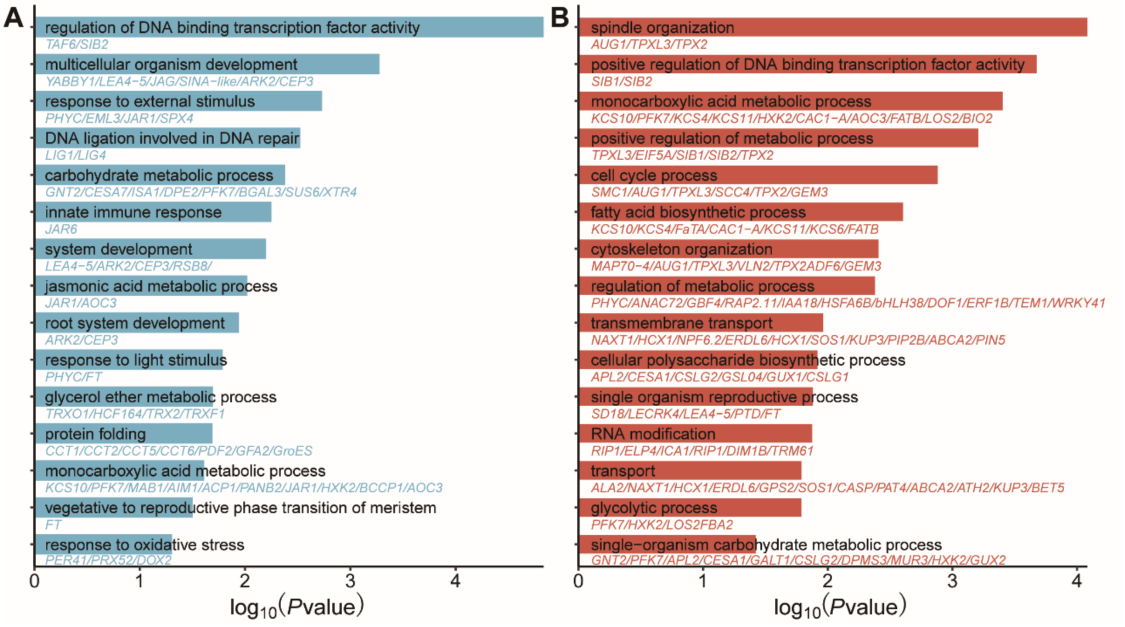


**Supplementary Figure 27. GO functional enrichment analysis of the selective genes for *O. longilobus* (A) and *O. taihangensis* (B).** The representative genes are shown below each of the fifteen GO biological process terms. For all annotated genes are provided as background information. Detailed information is shown in Supplementary Table 39.

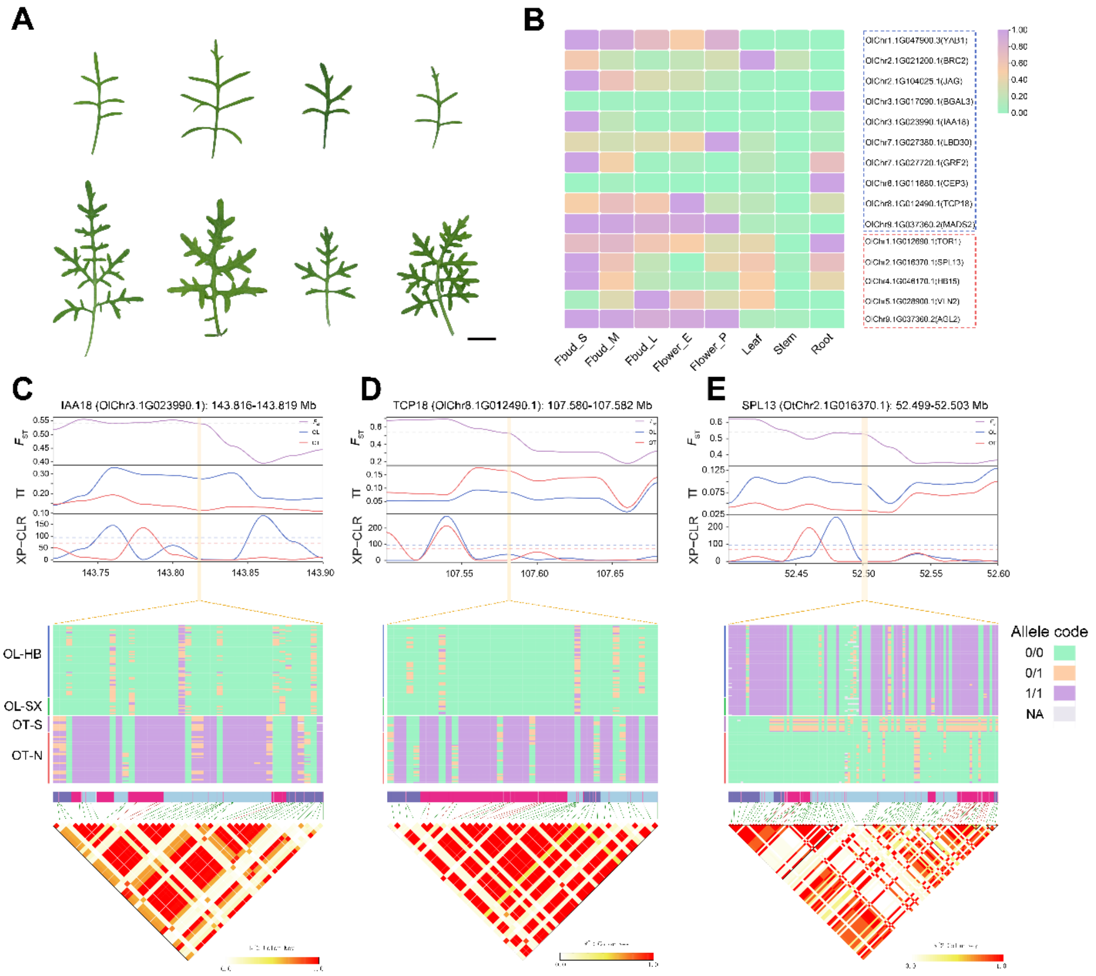


**Supplementary Figure 28. Selection signatures for the organ development genes. A,** Representative images showing leaf morphology of each four *O. longilobus* (top panel) and *O. taihangensis* (bottom panel) accessions. **B,** The tissue expression profiles of the identified selective genes for *O. longilobus* (dashed rectangular box highlighted in blue) and *O. taihangensis* (dashed rectangular box highlighted in red). **C-E,** Local distribution of $F_{ST}$, nucleotide diversity ($\pi$), XP-CLR values in the regions of 143.716-143.919 Mb (Chr3.1), 107.480-107.682 Mb (Chr8.1), and 52.399-52.603 Mb (Chr2.1), where selective genes *IAA18*, *TCP18* and *SPL13* are located (marked by yellow rectangular box), respectively. The central square heatmap shows the haplotypes for each of the three selective genes. The columns represent genotypes of SNPs, while the rows represent accessions. LD heatmap (bottom) shows the linkage degree of SNPs located in the genic region of selective genes. The nonsynonymous SNPs are highlight by red vertical lines.

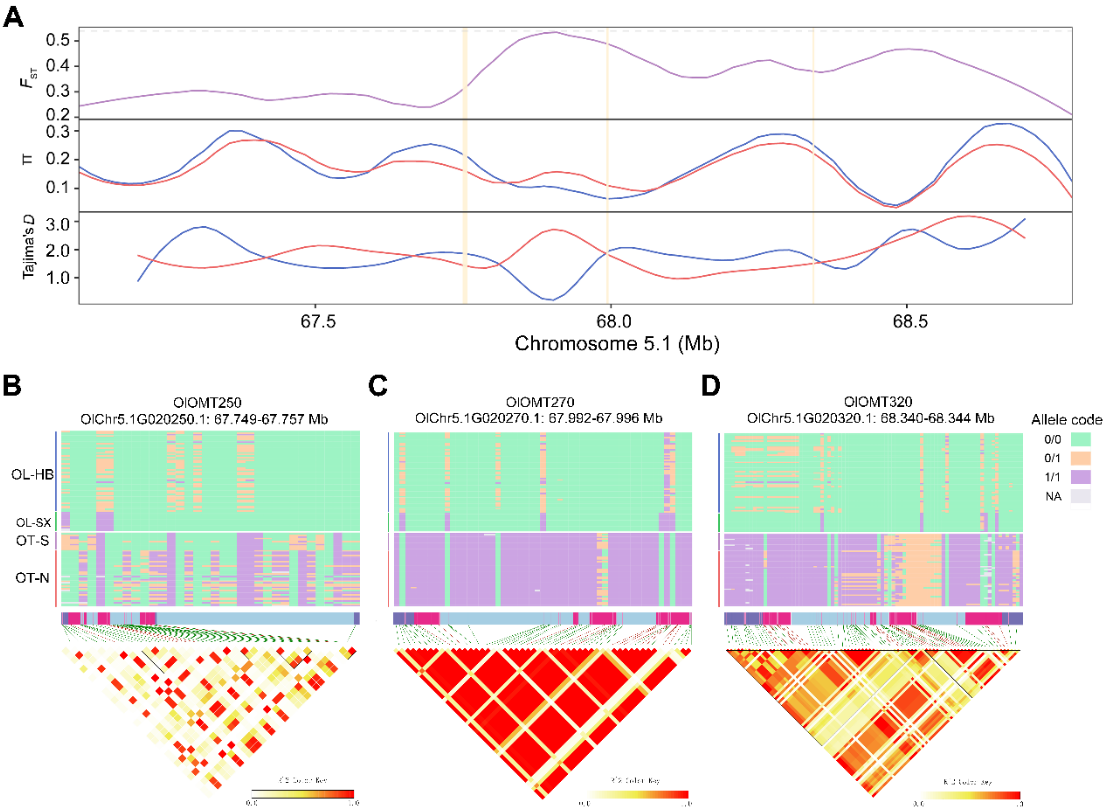


**Supplementary Figure 29. Selection signatures for three tandem repeat OMT genes on chromosome 5.1. A,** Local distribution of $F_{ST}$, nucleotide diversity ($\pi$), Tajima's *D* values in the regions of 67.100-68.794 Mb on chromosome 5.1, where three tandem repeat OMT genes *OlOMT250*, *OlOMT270* and *OlOMT320* are located (marked by yellow rectangular box), respectively. **B-D,** The central square heat map shows the haplotypes for each of the three tandem repeat OMT genes. The columns represent genotypes of SNPs. The rows represent accessions. LD heatmap (bottom) shows the linkage degree of SNPs located in the genic region of OMT genes. The nonsynonymous SNPs are highlight by red vertical lines.

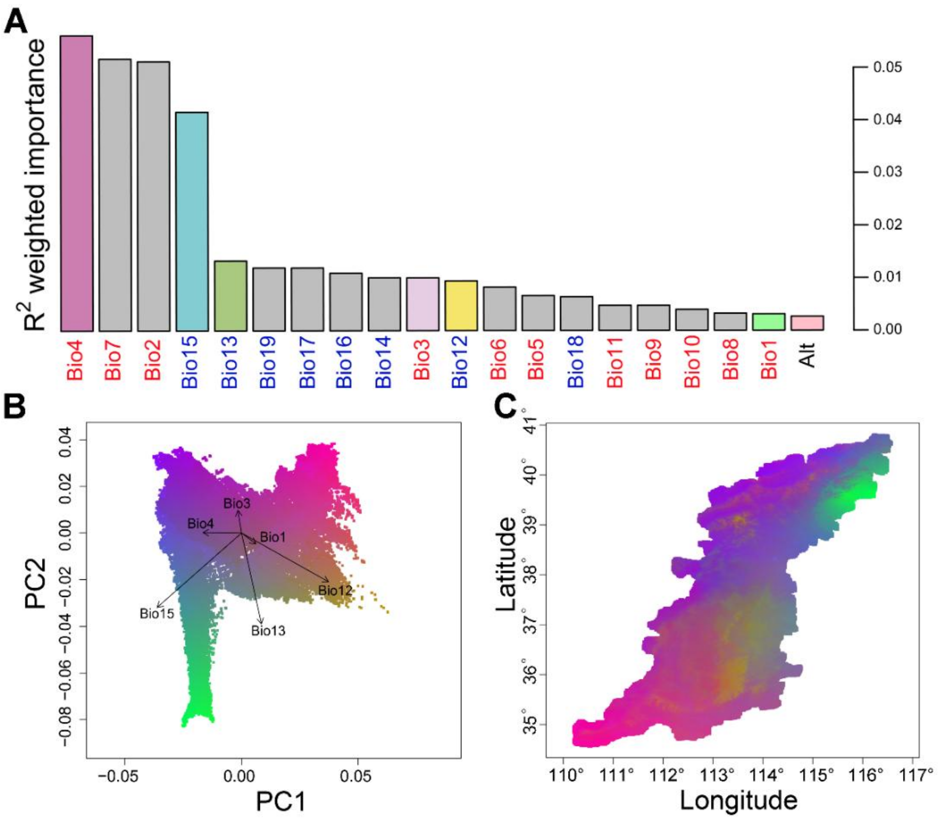


**Supplementary Figure 30. Importance ranking of environment variables and spatial distribution patterns of adaptive genetic variation within *Opisthopappus* genus using the GradientForest (GF) model. A,** $R^2$ weighted importance ranking of 20 environmental variables in GF models based on a dataset of 500,000 randomly selected SNPs after linkage disequilibrium (LD) filtering (plink --thin-count 500,000). The colored bar charts indicate the 7 representative variables selected for RDA analysis. **B,** The PCA analysis of adaptive genetic variation. Arrows indicate the loadings of 7 selected variables colored in **A**. Labeled vectors of the PCs present the direction and magnitude correlation. **C,** The visualization of the composition of adaptive genetic variation across the Taihang Mountain range in a red-green-blue (RGB) color scale. Similar colors denote similar expected genetic composition at adaptive variants.

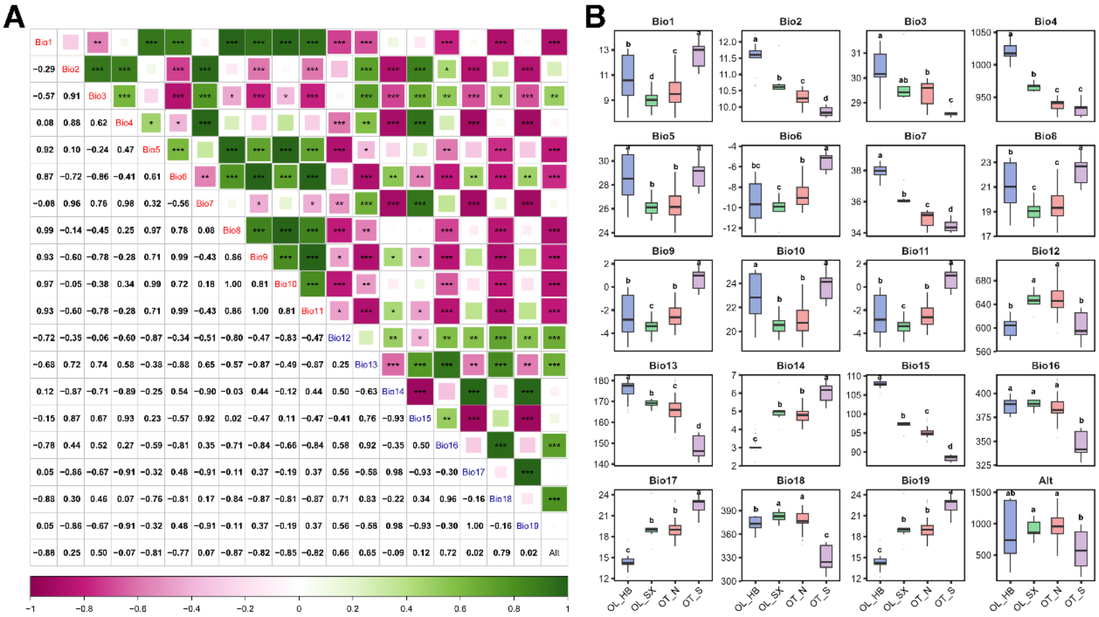


**Supplementary Figure 31. Correlation heatmap and distribution profiles of 20 environmental variables. A,** Pairwise Spearman's correlation coefficients among the 20 environmental variables. Temperature-related variables (Bio1-11) and precipitation-related variables (Bio12-19) shown in the diagonal are highlighted in red and blue, respectively. Alt, altitude. *** $P < 0.001$, ** $P < 0.01$, * $P < 0.05$. **B,** Comparison of 20 environmental phenotypes across four subgroups. The central line, edges and whiskers for each box correspond to the median, the 25th and 75th quartiles, 1.5 times the inter-quartile range (IQR), respectively. Values with different lowercase letters denote statistically significant differences ($P < 0.05$) based on a Student's *t*-test.

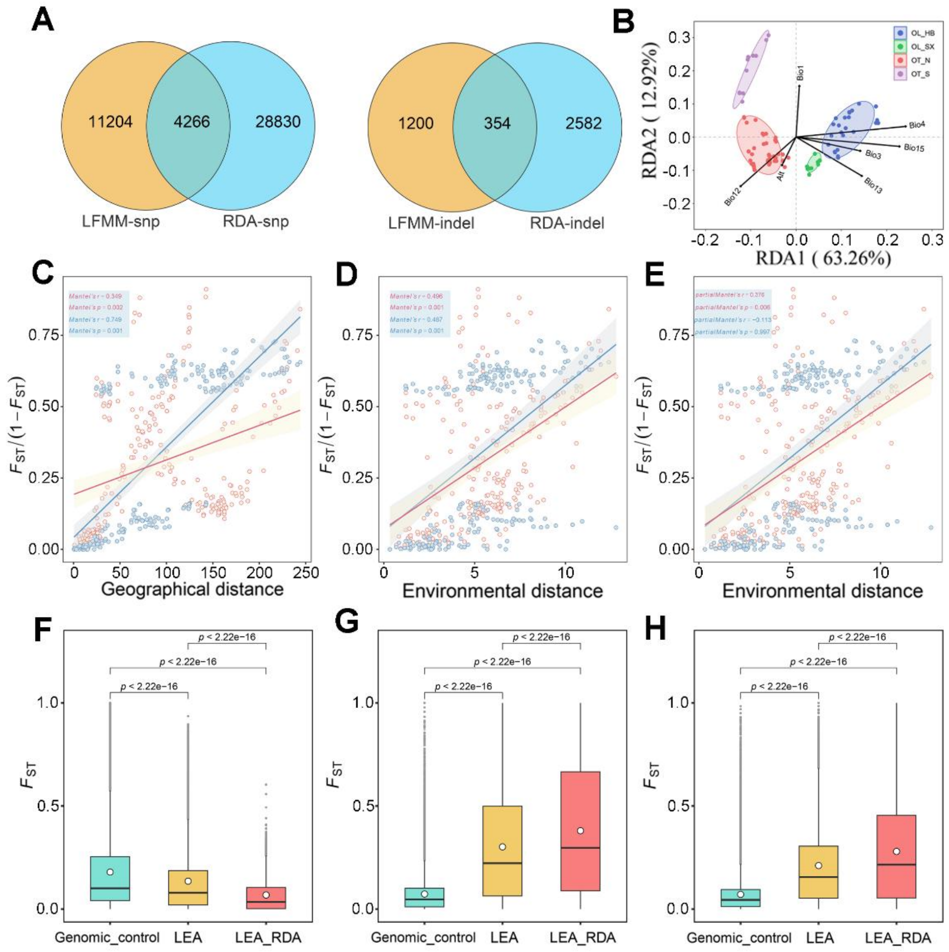


**Supplementary Figure 32. Genome-wide identification and analysis of the loci associated with local environmental adaptation.** **A,** Adaptive SNPs and InDels explored by latent factor linear mixed model (LFMM) and redundancy analysis (RDA). **B,** RDA reveals the relationships between the independent environmental variables and population structure of *Opisthopappus* along the RDA1 and RDA2 axes. Individuals are color-coded based on the four sub-populations determined by ADMIXTURE analysis. **C,** Isolation-by-distance (IBD) analysis (Mantel test, two-sided). **D,** Isolation-by-environment (IBE) analysis (Mantel test, two-sided). **E,** IBE analysis (partial Mantel test, two-sided). In **C-E,** the IBD or IBE analyses using the neutral and adaptive variants are distinguished by blue and red dots/lines, respectively. The shadow of linear regression denotes the 95% confidence interval. **F-H,** Comparison of genetic differentiation ($F_{ST}$) calculated by using 100,000 random variants (Genomic control), 17,024 environmental-associated variants detected by LFMM, and 4,620 core adaptive variants detected by both LFMM and RDA (LFMM & RDA) for group pairs of OL and OT, OL_HB and OL_SX, OT_N and OT_S, respectively. The centre lines of each box plot represents the medians, the top and bottom edges correspond to the 25th and 75th quartiles, and the whiskers extend to 1.5× the interquartile range. The gray points beyond the end of the whiskers are outliers. The average $F_{ST}$ value for each group is indicated as white dot. The Wilcoxon test (two-tailed) was used to determine the significance.

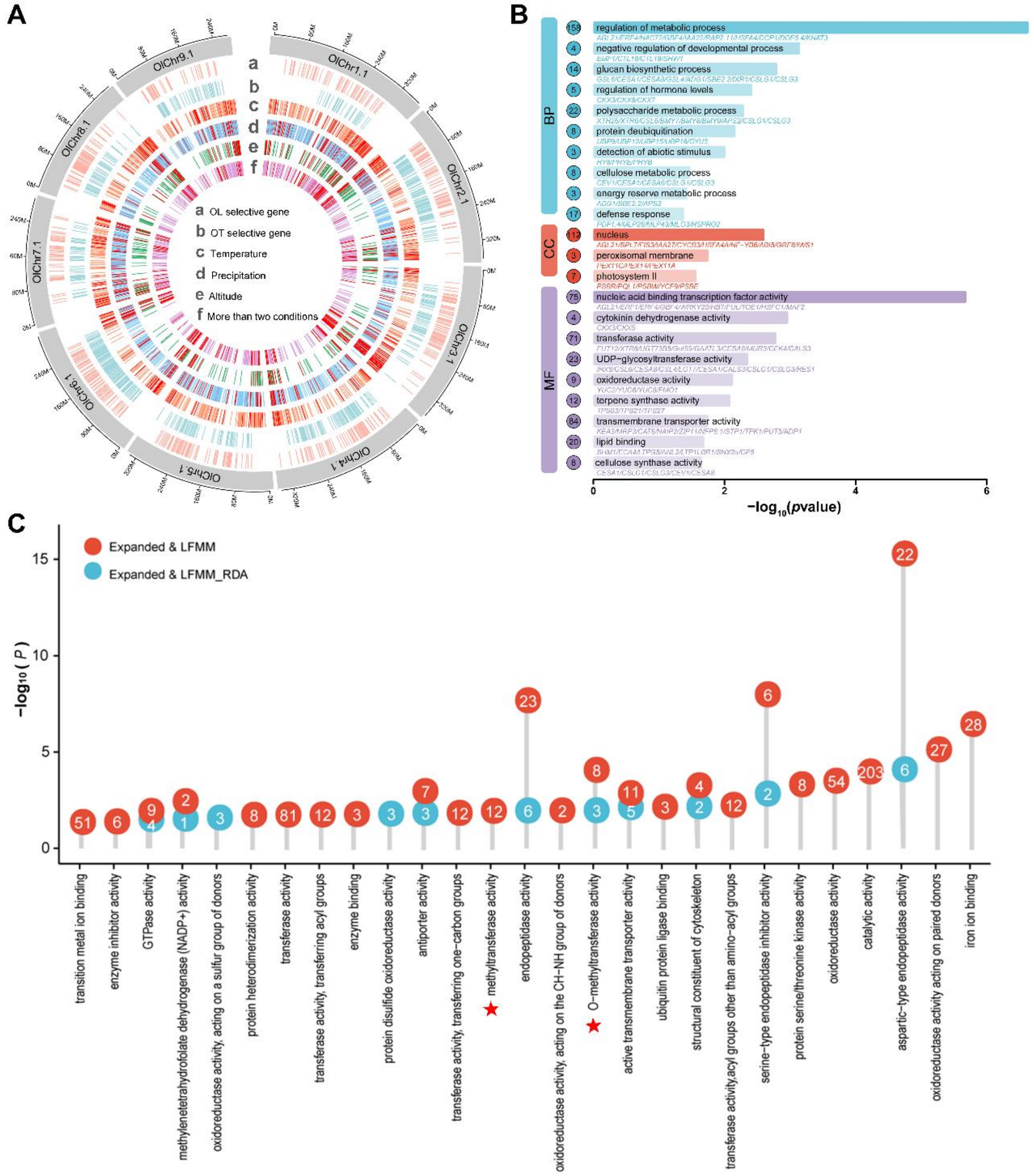


**Supplementary Figure 33. Adaptive and selective genes identified in *Opisthopappus*. A,** Circos plot showing the distribution of adaptive and selective genes across nine *O. longilobus* pseudochromosomes. Tracks from outer to inner indicate the following: **a, b,** the selective genes identified for *O. longilobus* and *O. longilobus*, respectively. **c-e,** the adaptive genes associated with temperature-, precipitation-related environmental variables and altitude, respectively; **f,** the adaptive genes co-located for at least two of the three above factors (temperature, precipitation and altitude). **B,** GO functional enrichment analysis of the 4,620 local environmental adaptive genes. The representative significant GO terms of biological process (BP), cellular component (CC) and molecular function (MF) are presented. The numbers inside the circle represent the number of genes in the corresponding GO term. **C,** The enriched GO terms in biological processes for the adaptive genes overlapping with expanded genes of *O. longilobus*. The blue and red points represent the GO terms enriched for 190 and 629 expanded genes identified by LFMM & RDA and LFMM approaches, respectively. The white numbers indicate the number of genes in the corresponding GO term. Detailed information is shown in Supplementary Tables 45 and 54.

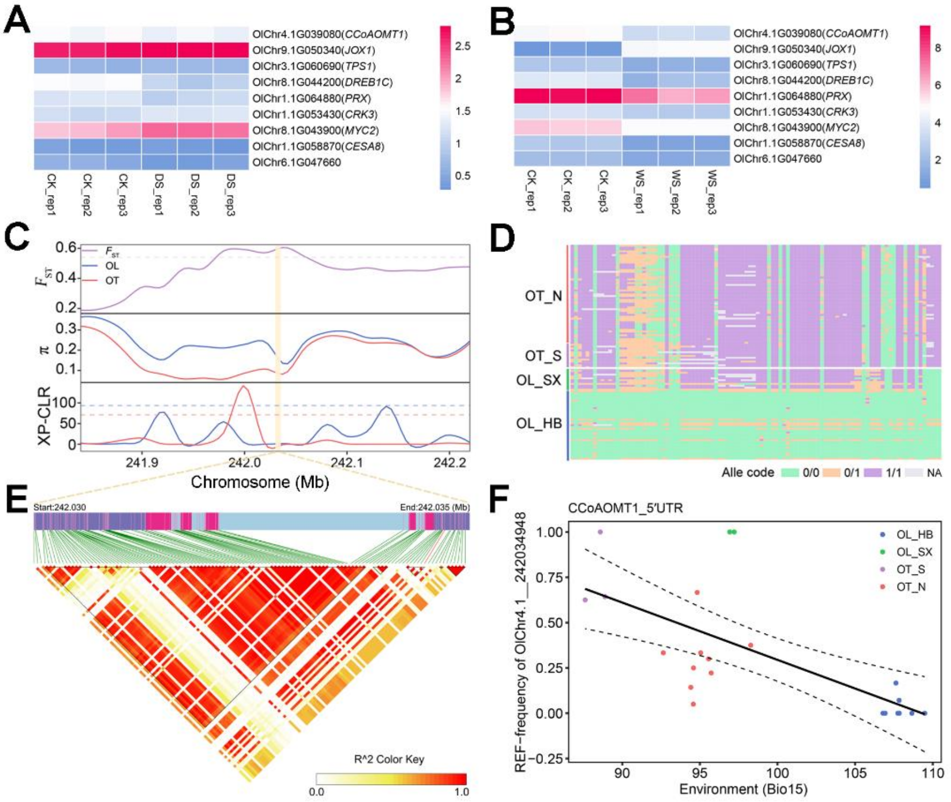


**Supplementary Figure 34. Expression levels of the environmental adaptive genes and selective signal for *CCoAMOT1*. A, B,** Gene expression profiles of nine selected adaptive genes (in $\log_2$(RPKM+1)) under drought stress (DS) and waterlogging stress (WS) conditions, respectively. **C,** Selective sweep on chromosome 4.1 (241.830-242.235 Mb), where adaptive gene *CCoAMOT1* is highlighted with yellow shadow. **D,** Genotype heatmap of *CCoAMOT1*. **E,** Linkage disequilibrium heatmap of *CCoAMOT1*. A total of 118 variants located in the genic region of *CCoAMOT1*, and one InDel loci OlChr4.1__242034948 (5’ UTR, c.-106_-105insC) associated with Bio15 (precipitation seasonality). **F,** Linear regression of allele frequencies and environment variable Bio15 across 22 populations for OlChr4.1__242034948. Different color points represent the four *Opisthopappus* subpopulations. The dashed line represents the 95% confidence interval.

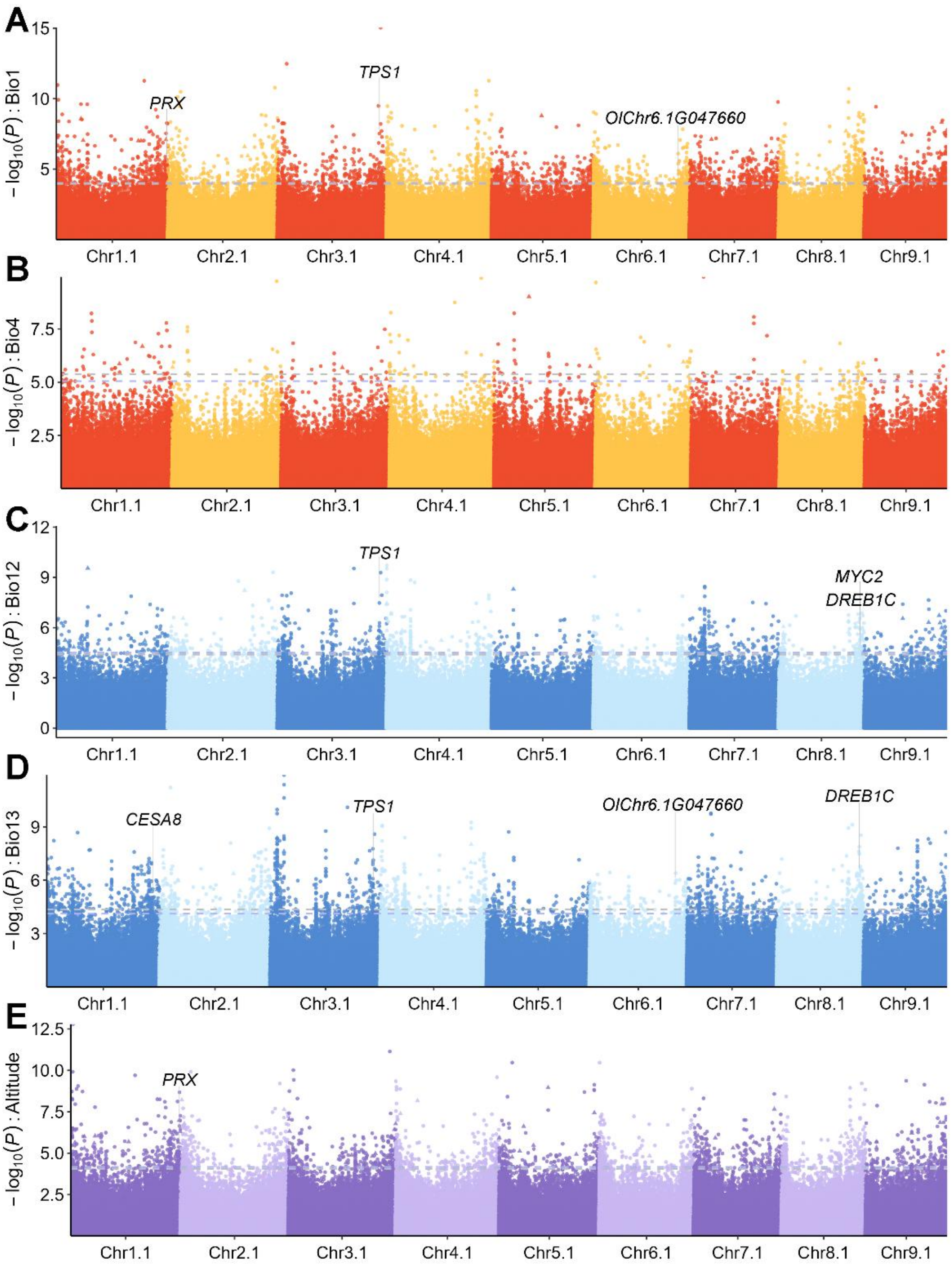


**Supplementary Figure 35. Manhattan plots show the genotype-environment associations identified for the representative variables (Bio3 and Bio15 are not included) by LFMM.** SNPs and InDels are represented by points and triangles, respectively. The blue and gray dashed lines indicates the 5% false discovery rate correction thresholds for SNPs and InDels, respectively.

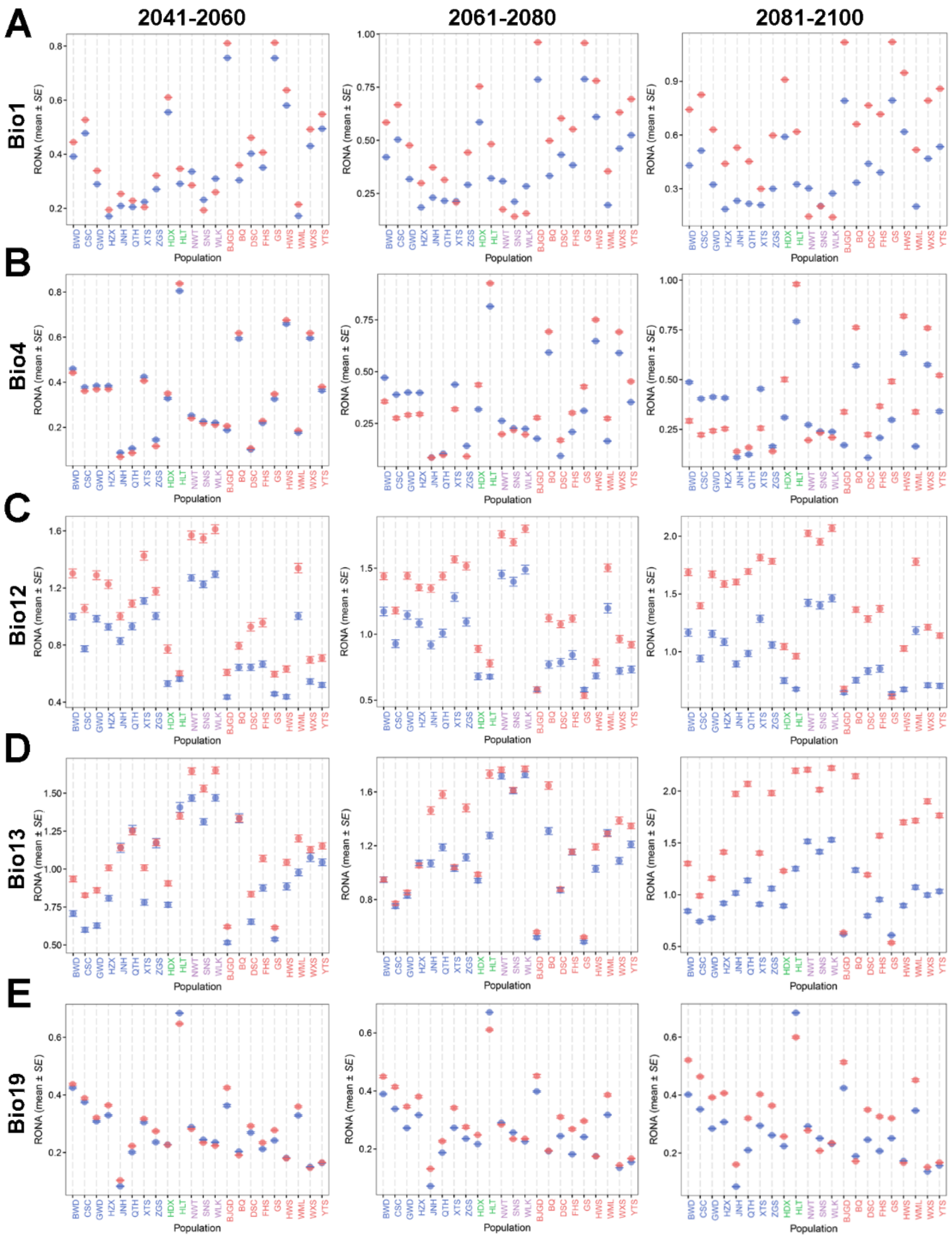


**Supplementary Figure 36. Identification of gene variants associated with local environmental adaptation in *Opisthopappus*.** Upper panel: The climate-associated variants are marked in schematic diagram of the gene structure with green vertical line segments, and the pink points indicate the proposed functional site. Lower panel: Linear regression of allele frequencies and environment variables across 22 *Opisthopappus* populations for the proposed functional variant. The four subpopulations are distinguished by color-coded points. The dashed line represents the 95% confidence interval.

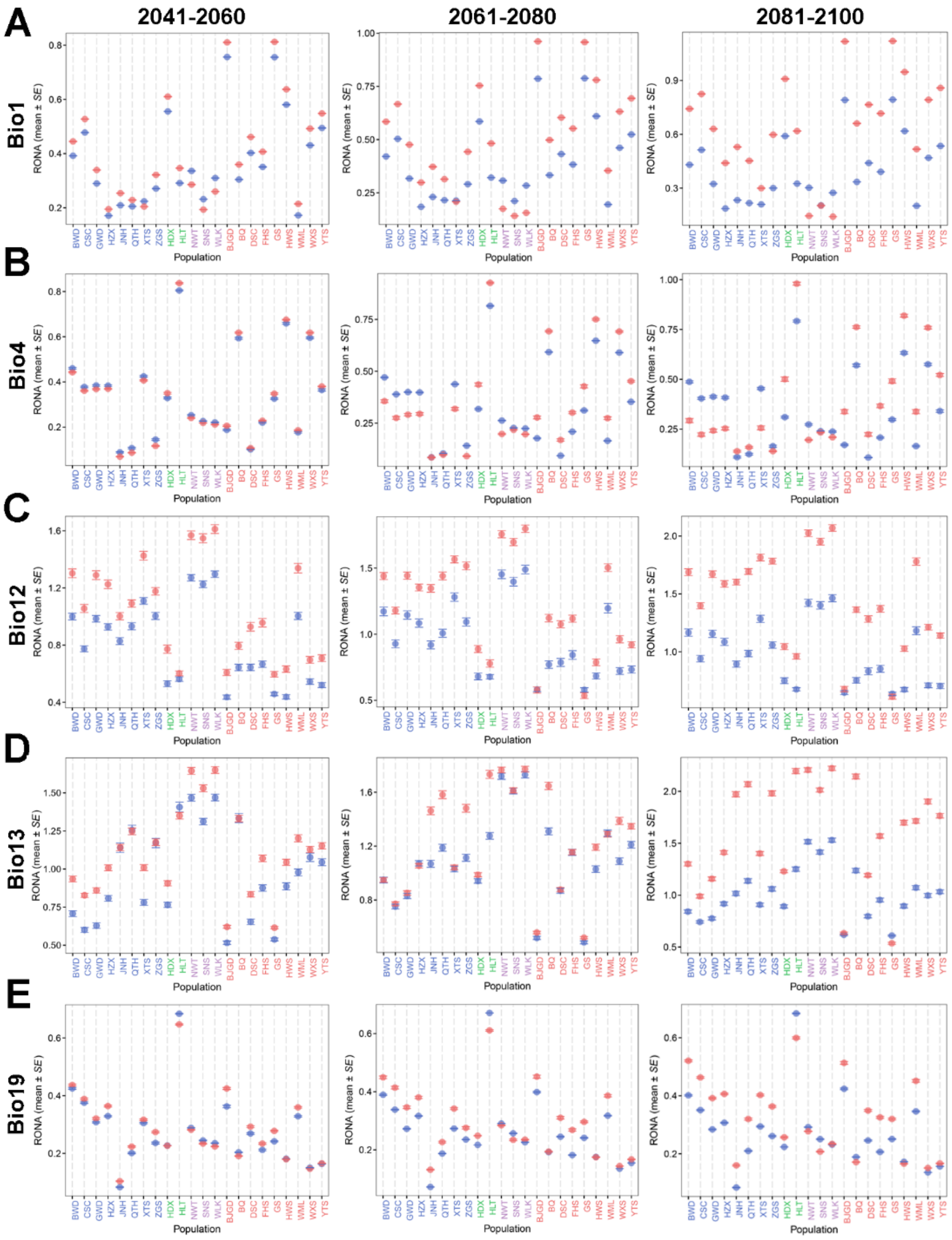


**Supplementary Figure 37. Comparison of RONA values across the 22 *Opisthopappus* populations under different future climate scenarios.** In **A-E,** The blue and red points represent the average RONA estimates for Bio1 (n = 916 climate-associated variants), Bio4 (n = 28 climate-associated variants), Bio12 (n = 262 climate-associated variants), Bio13 (n = 555 climate-associated variants), and Bio19 (n = 96 climate-associated variants) under SSP126 and SSP585, respectively. Error bars represent the standard error of the average RONA values calculated from different variants. Source data are provided in Supplementary Table 46.

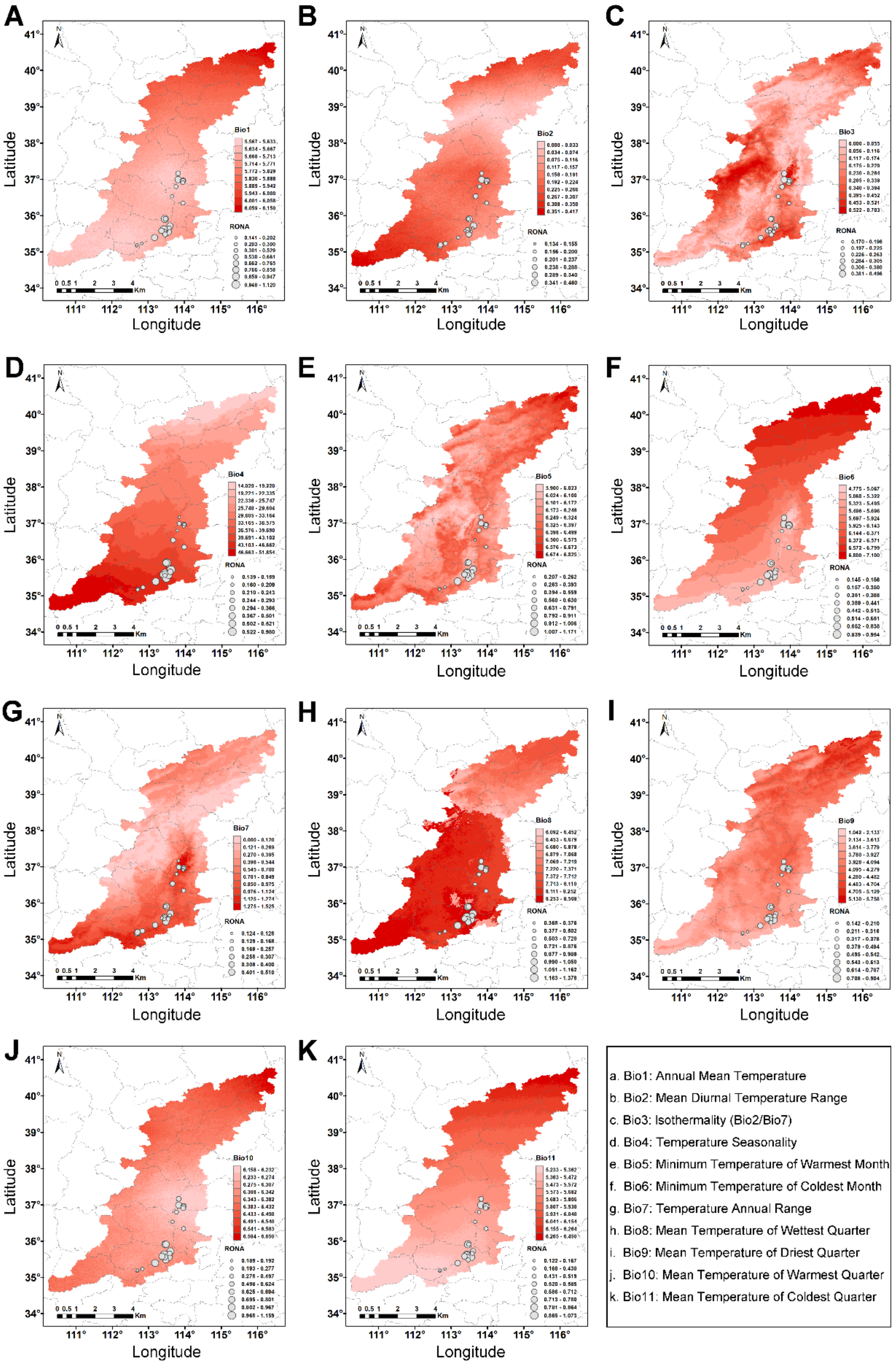


**Supplementary Figure 38. Comparison of the RONA estimates across 22 *Opisthopappus* populations for 11 temperature-related variables (Bio1-11) under SSP585 scenario in 2081-2100.** The raster colors on the maps indicate the degree of projected absolute climate change across the Taihang Mountain range, which were calculated by subtracting the average current climate data from average future climate data of 4 climate models. Areas with darker red are forecasted to experience more dramatic change. The point sizes on the map correspond to RONA values across *Opisthopappus* populations.

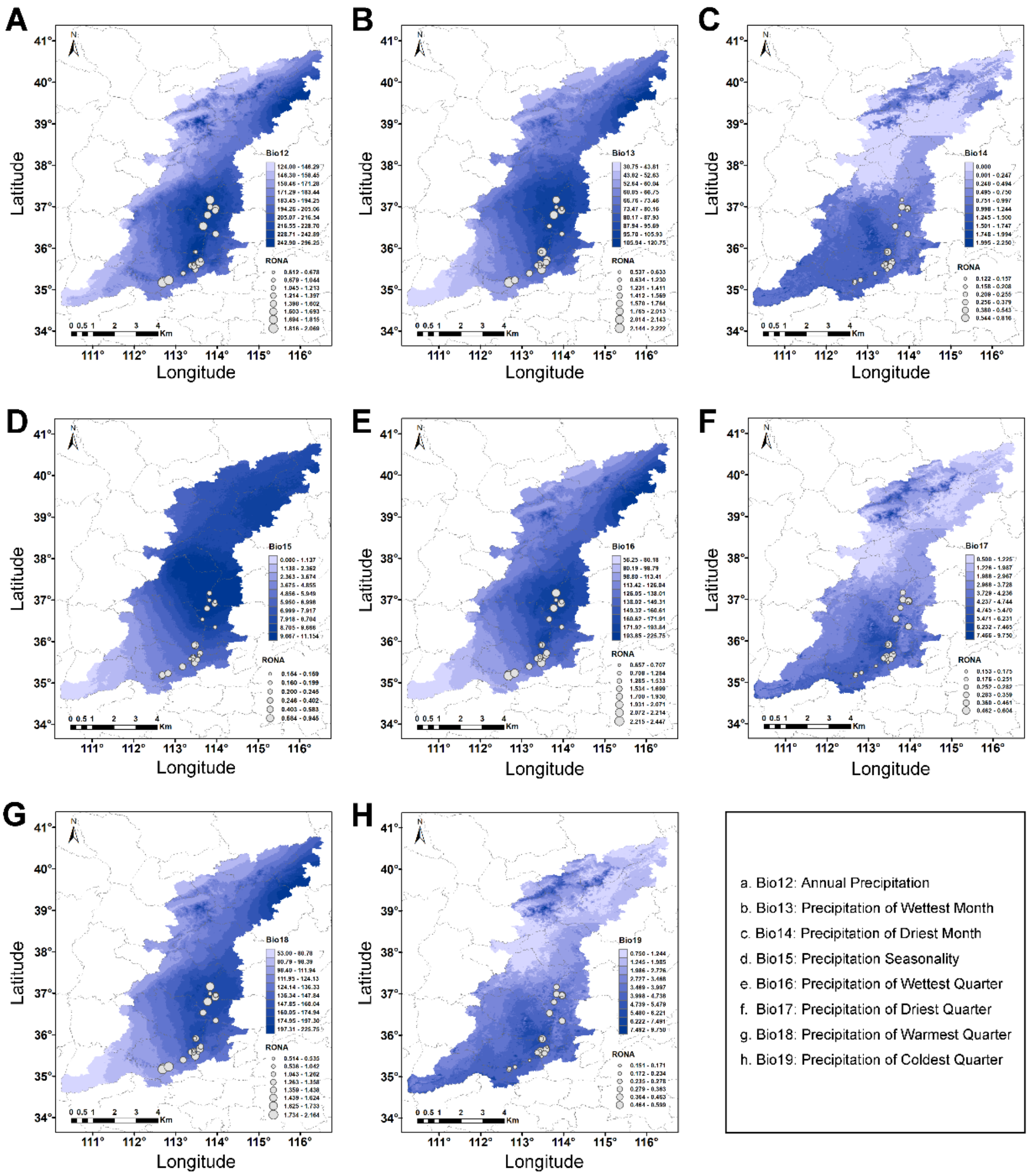


**Supplementary Figure 39. Comparison of the RONA estimates across 22 *Opisthopappus* populations for 8 precipitation-related variables (Bio12-19) under SSP585 scenario in 2081-2100.** The raster colors on the maps indicate the degree of projected absolute climate change across the Taihang Mountain range, which were calculated by subtracting the average current climate data from average future climate data of 4 climate models. Areas with darker blue are forecasted to experience more dramatic change. The point sizes on the map correspond to RONA values across *Opisthopappus* populations.

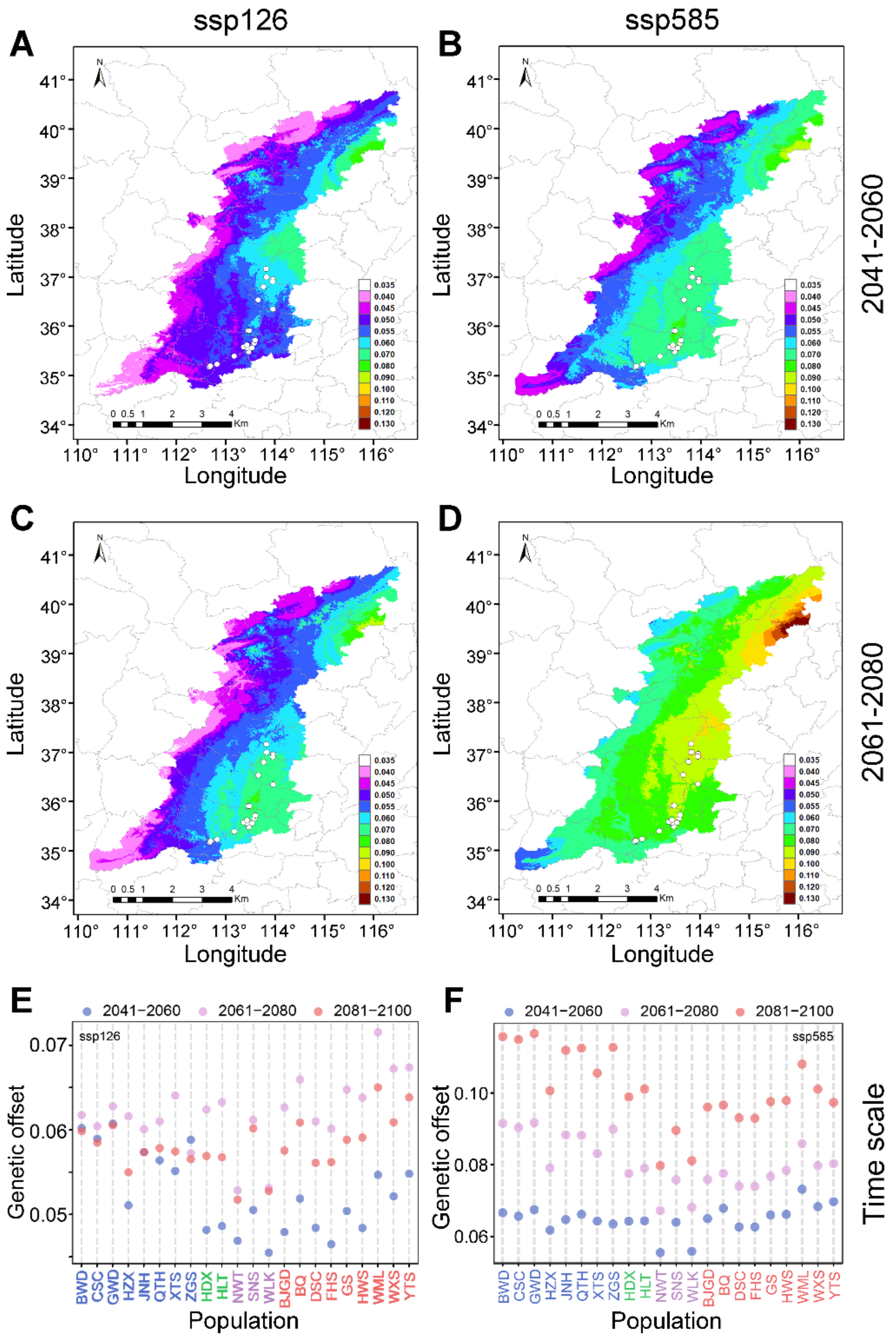


**Supplementary Figure 40. Predicted genetic offsets to future climate change in *Opisthopappus* based on 4,620 core adaptive loci. A-D,** Map of the GF-predicted genetic offsets averaged across four climate models across the Taihang Mountain range (n = 200,824 grids) under two shared socioeconomic pathways (SSP126 and SSP585) and two future periods (2041-2060 and 2061-2080) scenarios. The color gradient from white to terracotta refers to increasing levels of genetic offset, and the 22 sampled populations are denoted by circular markers on the map. **E, F,** Temporal genetic offset predictions for 22 *Opisthopappus* populations under SSP126 (low emissions) and SSP585 (high emissions) scenarios, respectively. Source data are provided in Supplementary Table 47.

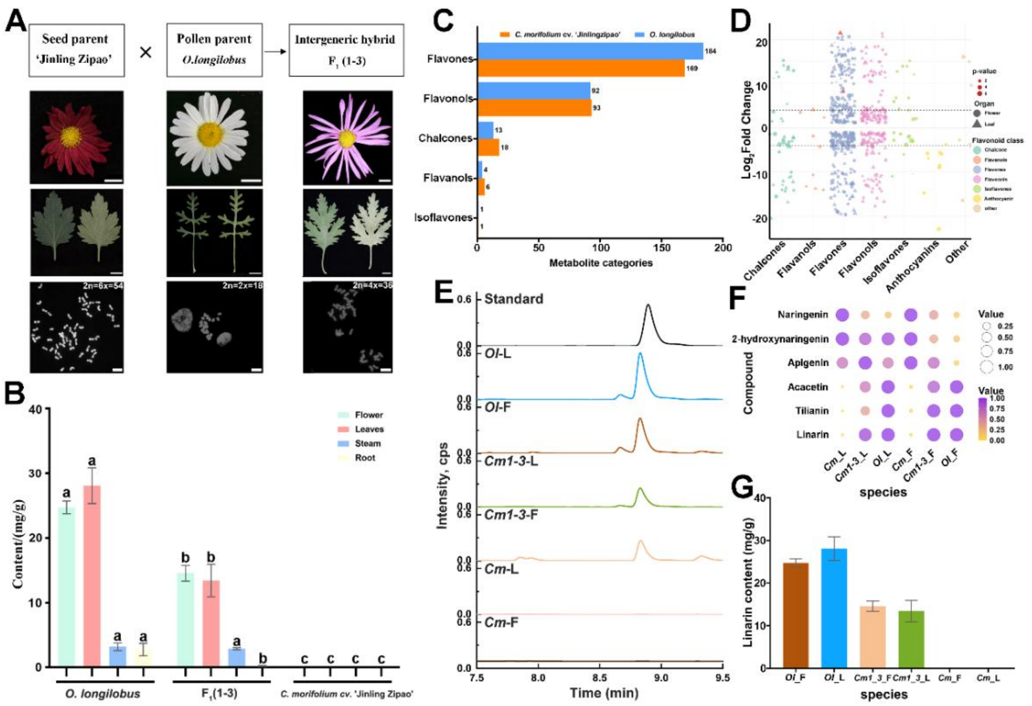

**Supplementary Figure 41. Intergeneric hybridization and metabolite analysis. A,** Flowchart of the intergeneric hybridization process. The flower, leaf, and chromosomal karyotype characteristics of *C. morifolium* cv. 'Jinling Zipao', *O. longilobus*, and their distant hybrid offspring $F_1$(1-3). **B,** Tissue-specific distribution of linarin content in *O. longilobus*, *C. morifolium* cv. 'Jinling Zipao', and $F_1$(1-3). The statistical significance was evaluated using two-way ANOVA across different tissues. All data are presented as mean ± standard deviation (mean ± *SD*, n = 3). Significance levels were set as follows: $P < 0.05$ indicates a significant difference (*), $P < 0.01$ indicates a highly significant difference (**), and $P < 0.001$ indicates an extremely significant difference (****). NS, no significance. **C,** Metabolite composition and proportion in *O. longilobus* and *C. morifolium* cv. 'Jinling Zipao' determined by flavonoid-target metabolome analysis (n = 3 biological replicates). **D,** Differential metabolite analysis between *O. longilobus* and *C. morifolium* cv. 'Jinling Zipao'. **E,** UPLC chromatographic analysis of linarin content in *O. longilobus* (*Ol*), *C. morifolium* cv. 'Jinling Zipao' (*Cm*), and hybrid $F_1$(1-3) (*Cm*1-3). F: flower; L: leaf. **F,** The relative levels of target metabolites in the flower (F) and leaf (L) of parents (*Cm* and *Ol*) and hybrid $F_1$(1-3) (*Cm*1-3) were quantified using flavonoid-targeted metabolomics. Metabolite levels in flower and leaf tissues were normalized to those in the corresponding tissues of *O. longilobus*. **G,** Statistical analysis of linarin content quantified by UPLC in *O. longilobus*, *C. morifolium* cv. 'Jinling Zipao', and hybrid $F_1$(1-3). Results are shown as means ± *SD* (n = 3 biological replicates). Source data are provided in Supplementary Tables 50 and 51.

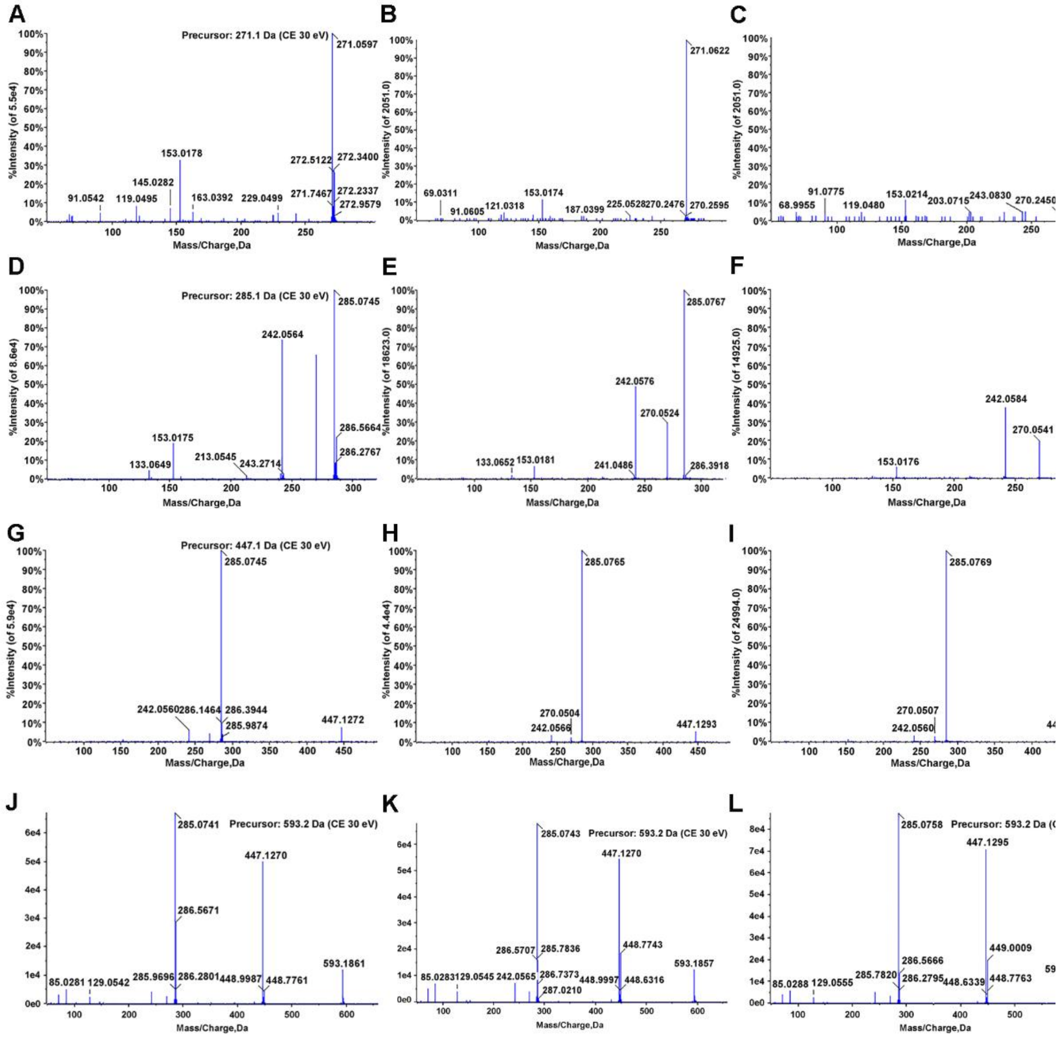

**Supplementary Figure 42. MS2 fragment ions analysis in *O. longilobus*. A, D, G, J,** MS2 fragment ions of apigenin, acacetin, tilianin and linarin standards. **B, C,** Mass spectra of apigenin in flower and leaf tissues. **E, F,** Mass spectra of acacetin in flower and leaf tissues. **H, I,** Mass spectra of tilianin in flower and leaf tissues. **K, L,** Mass spectra of linarin in flower and leaf tissues.

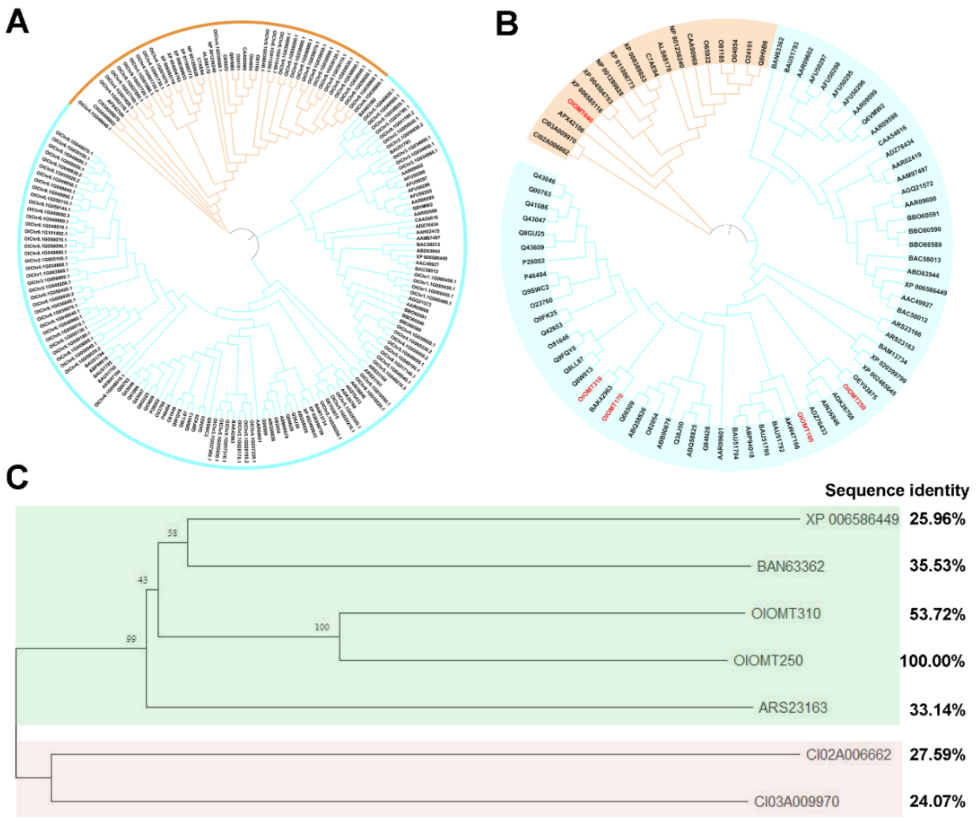


**Supplementary Figure 43. Analysis of OMT genes of *O. longilobus*. A,** A phylogenetic analysis of COMTs and CCoAOMTs identified in *O. longilobus* and other species. **B**, Phylogenetic tree of candidate OMTs in *O. longilobus* and other species. CCoAOMT and COMT genes are colored lightblue and orange, respectively. In **B,** The OMT genes marked in red indicate *O. longilobus* candidates clustered with known OMTs. **C,** Phylogenetic analysis based on protein sequence alignments of previously reported enzymes and the newly identified OlOMT250 and OlOMT310 in this study. Previously reported OMT enzymes used for phylogenetic tree construction included *Plagiochasma appendiculatum* (Pa4'OMT, Genebank: ARS23163), *Glycine max* (SOMT2, Genebank: XP006586449), *Carthamus tinctorius* (CtMROMT, Genebank: Ban63362), and *C. indicum* (CI03A009970 and CI02A006662). Detailed information about the reference OMT sequences used is available in Supplementary Table 53.

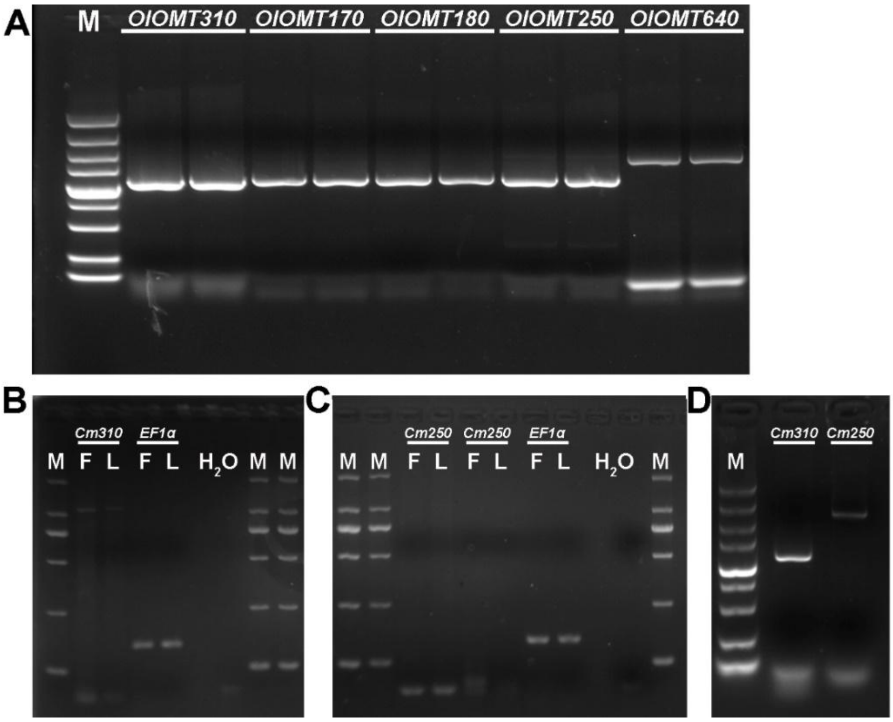


**Supplementary Figure 44. Cloning of candidate OMT genes in *O. longilobus* and *C. morifolium* cv. 'Jinling Zipao'. A,** Cloning of candidate OMT genes from *O. longilobus*. **B,** Cloning of *CmOMT310* gene from *C. morifolium* cv. 'Jinling Zipao' (cDNA as template). **C,** Cloning of *CmOMT250* gene from *C. morifolium* cv. 'Jinling Zipao' (cDNA as template). **D,** Cloning of *Cm310* and *Cm250* genes from *C. morifolium* cv. 'Jinling Zipao' (DNA as template). F and L denote flower and leaf tissues, respectively. *EF1α* is used as the reference gene (GenBank accession number: KF305681). For clarity in this figure, *CmOMT310* and *CmOMT250* are abbreviated as *Cm310* and *Cm250*, respectively.

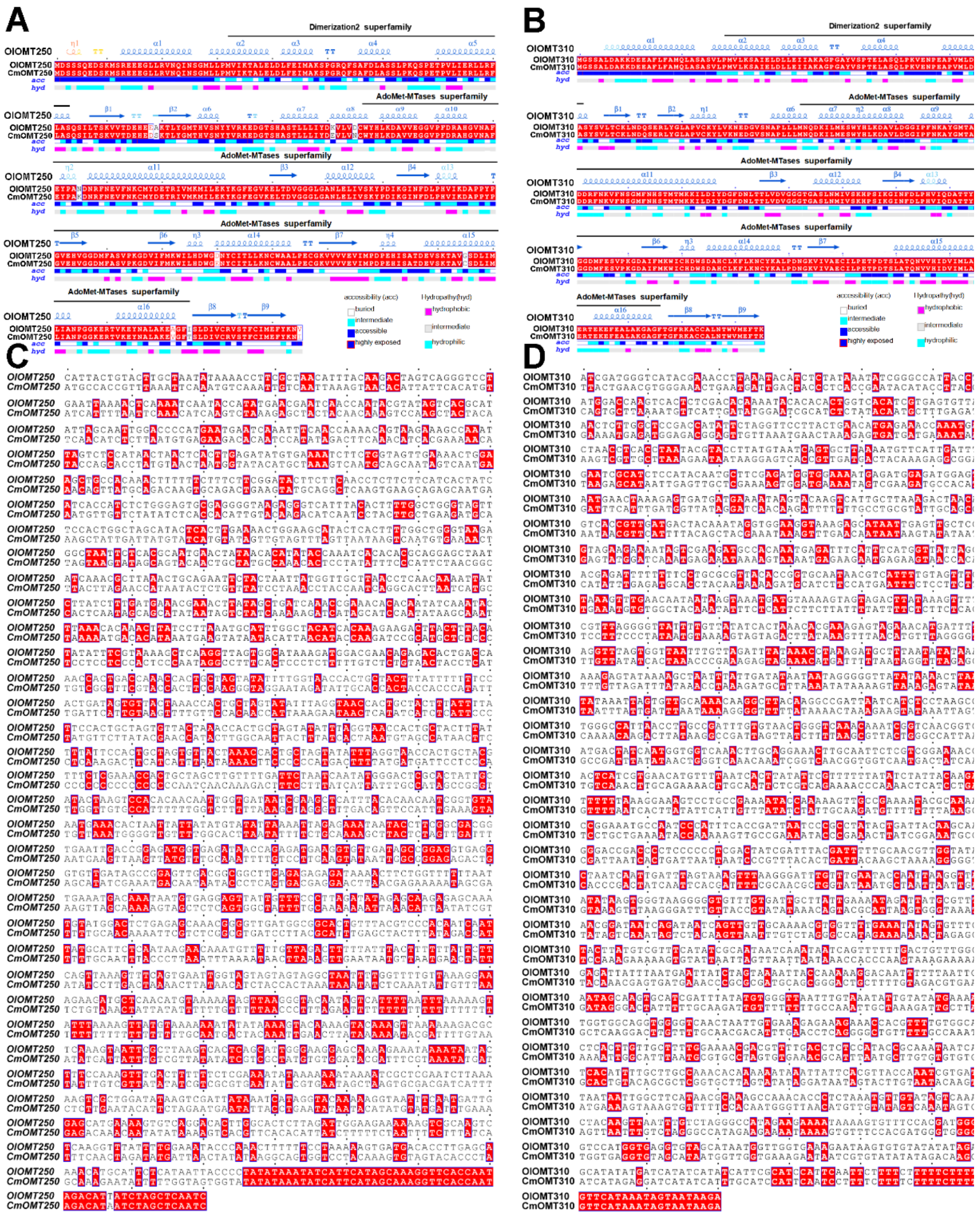

**Supplementary Figure 45. Sequence alignment of *OMT250* and *OMT310* between *O. longilobus* and *C. morifolium* cv. 'Jinling Zipao'. A, B,** Coding sequence (CDS) alignments of OMT250 and OMT310 between *O. longilobus* and *C. morifolium* cv. 'Jinling Zipao'. **C, D,** Promoter sequence alignments of *OMT250* and *OMT310* between *O. longilobus* and *C. morifolium* cv. 'Jinling Zipao'.

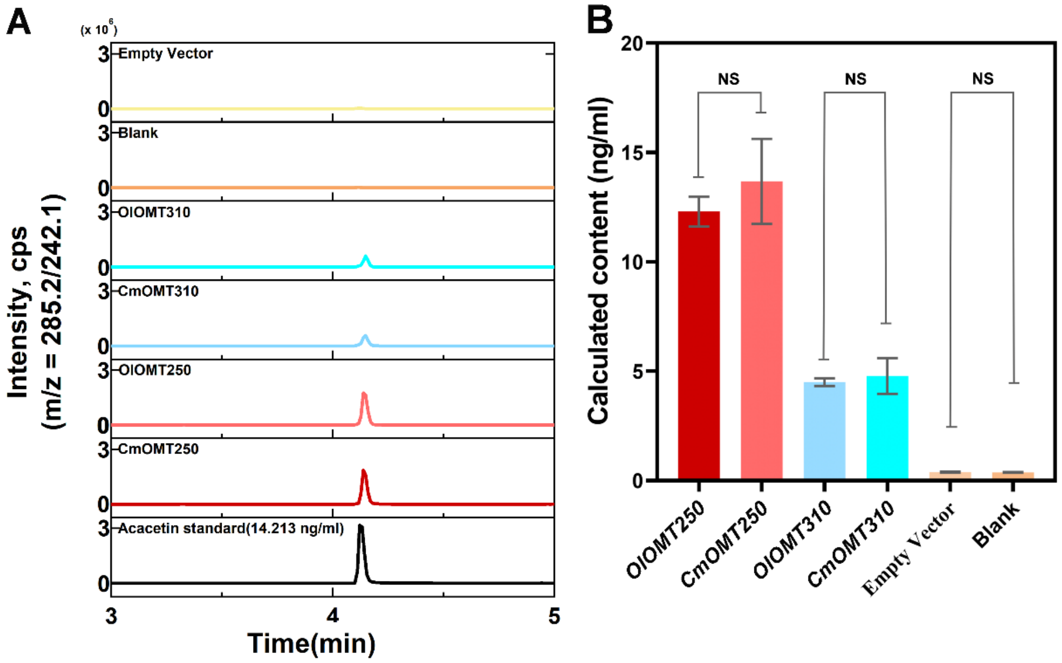


**Supplementary Figure 46. Comparative analysis of acacetin accumulation associated with OMT250/OMT310 between *O. longilobus* and *C. morifolium* cv. 'Jinling Zipao'. A,** Chromatographic analysis of acacetin accumulation associated with OMT250/OMT310 between *O. longilobus* and *C. morifolium* cv. 'Jinling Zipao'. **B,** Statistical analysis of acacetin accumulation associated with OMT250/OMT310 between *O. longilobus* and *C. morifolium* cv. 'Jinling Zipao'.

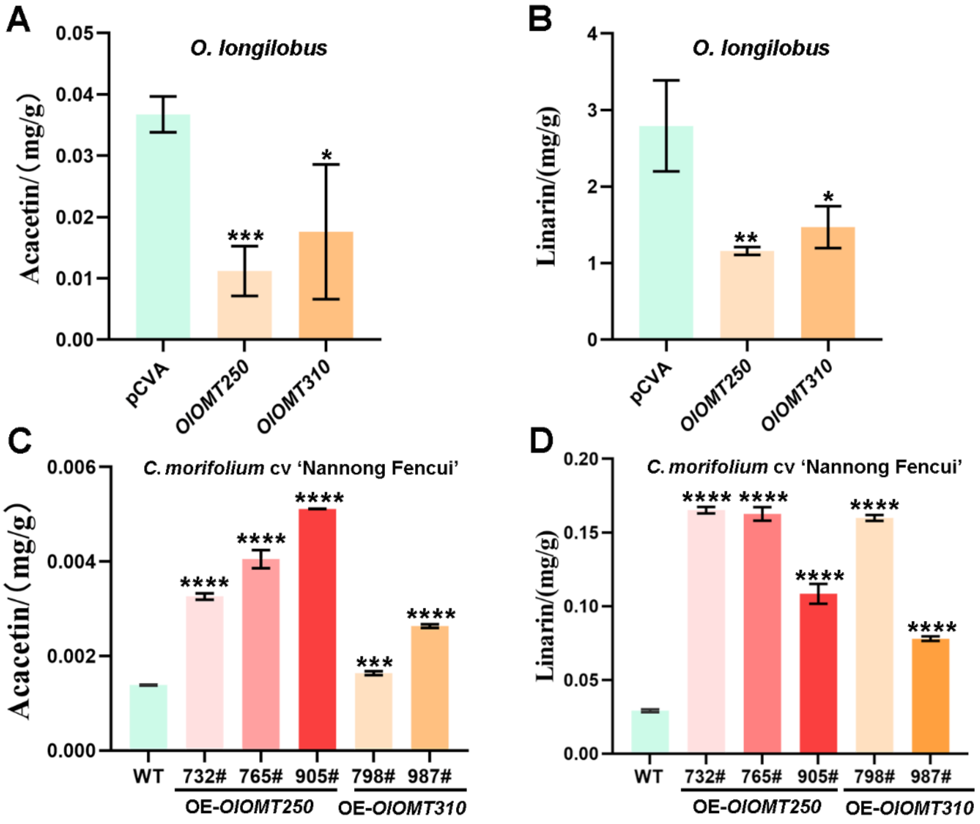


**Supplementary Figure 47. Relative contents of acacetin and linarin in *OMT250* and *OMT310* transient interference and overexpressing lines. A,** Quantification of acacetin content in seedling-stage transient VIGS plants of *O. longilobus* targeting *OlOMT250* and *OlOMT310* using the pCVA vector. **B,** Quantification of linarin content in seedling-stage transient VIGS plants of *O. longilobus* targeting *OlOMT250* and *OlOMT310* using the pCVA vector. **C,** Quantification of acacetin content in wild-type (WT) and OlOMT250/OlOMT310 overexpression lines. **D,** Quantification of linarin content in WT and *OlOMT250*/*OlOMT310* overexpression lines. Due to the lack of an established stable transformation system in *O. longilobus* and the chrysanthemum cultivar 'Jinling Zipao', transient gene silencing was performed in *O. longilobus* cuttings. In parallel, transient overexpression assays were conducted in the cultivated chrysanthemum 'Nannong Fencui', a genotype with low basal linarin accumulation, to facilitate functional assessment of candidate genes. All data are presented as mean ± *SD* (n = 3). Statistical significance was determined using Student's *t*-test: $P < 0.05$, significant (*), $P < 0.01$, highly significant (**), $P < 0.001$ indicates a highly significant difference (***), and $P < 0.0001$ indicates an extremely significant difference (****).

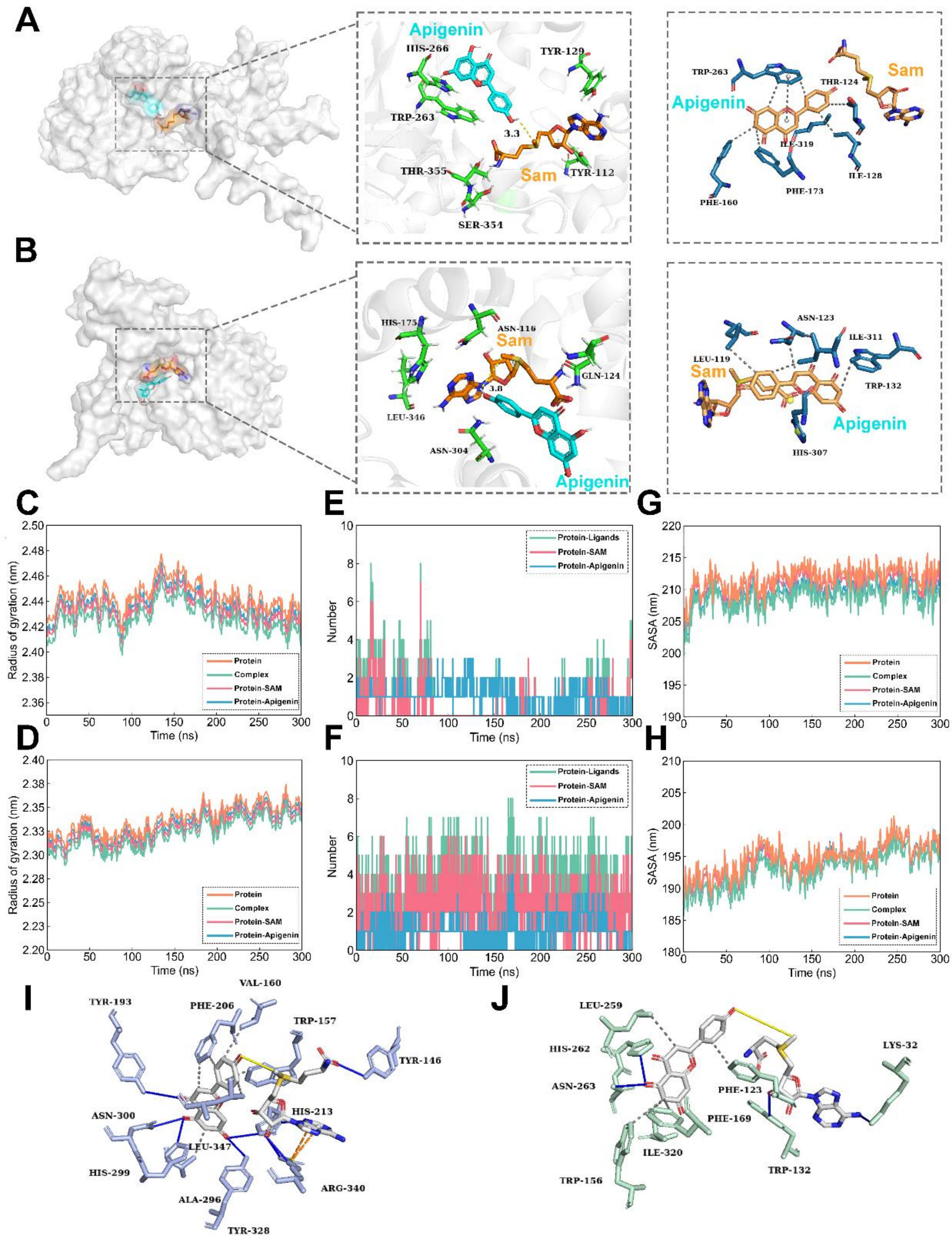


**Supplementary Figure 48. Molecular docking and dynamics analysis of OlOMT250, OlOMT310 and representative OMTs. A, B,** Molecular docking results of OlOMT250 and OlOMT310. **C, D,** Rotational radius of OlOMT250 and OlOMT310. **E, F,** Number of hydrogen bonds in OlOMT250 and OlOMT310. **G, H,** Accessible surface area of OlOMT250 and OlOMT310. **I,** Molecular docking results of Pa4'OMT protein. **J,** Molecular docking results of SOMT2 protein. Pa4'OMT and SOMT2 have been reported to catalyze the 4'-O-methylation step in apigenin biosynthesis (Kim et al., 2005; Liu et al., 2017).

### Supplementary Note

#### Supplementary Note 1. Genome sequencing and survey of *O. longilobus*

##### Genomic DNA extraction and quality assessment

The extracted DNA was quantified using a Qubit Fluorometer (Life Technologies, USA). DNA purity was assessed by measuring OD260/OD280 and OD260/OD230 ratios with a NanoDrop spectrophotometer (Thermo Fisher Scientific, USA), and DNA integrity was verified by 1% agarose gel electrophoresis. All sequencing was performed by BGI Genomics Corporation Ltd (Shenzhen, China).

##### Short-read library construction and sequencing

High-quality genomic DNA was fragmented to an average size of ~350 bp using a Covaris M220 ultrasonicator (Covaris, USA). Paired-end libraries were constructed using a standard protocol involving end repair, A-tailing, and adaptor ligation. Sequencing was carried out on the DNBSEQ-2000 platform (MGI Tech, China) with a paired-end 150 bp configuration, yielding approximately 390 Gb of raw short-read data. Low-quality reads, adapter contaminants, and reads containing ambiguous bases were removed using Fastp v0.20.0 (Chen, 2023) with platform-specific adapters (--adapter_sequence AAGTCGGAGGCCAAGCGGTCTTAGGAAGACAA; --adapter_sequence_r2 AAGTCGGATCGTAGCCATGTCGTTCTGTGAGCCAAGGAGTTG). Reads with an average Q-score < 15 or length < 150 bp were discarded. After filtering, 389.6 Gb of high-quality clean reads were retained for genome assembly.

##### PacBio library construction and sequencing

High-quality genomic DNA was used to construct 20 kb SMRTbell libraries, and PacBio sequencing was performed on the Sequel II platform. Genomic DNA was fragmented using a megaruptor system (Diagenode, Belgium) and size-selected for 13-16 kb fragments with the SageELF system (Sage Science, USA). DNA fragments were ligated with hairpin-shaped PacBio adapters to form SMRTbell templates for sequencing on the Sequel II platform using single molecule real-time (SMRT) technology. Raw subreads were processed with the CCS module (SMRT Link v10, CCS v6.2.0) to generate high-fidelity (HiFi) reads, using the parameters: "--min-passes 3 --min-rq 0.99 --min-length 500". In total, eight 20 kb libraries were constructed, yielding 248.84 Gb of high-quality HiFi clean data (Chin et al., 2013).

##### Hi-C library construction and sequencing

Fresh young leaves of *O. longilobus* were cross-linked with formaldehyde, and the cross-linked chromatin was lysed, digested with the restriction enzyme *MboI*, subjected to proximity ligation, and purified to obtain ligation products between spatially associated fragments. The Hi-C libraries were sequenced on the DNBSEQ-2000 platform, generating a total of 412.04 Gb of paired-end reads. Adapter contamination and low-quality reads were removed using fastp v0.20.0 (Chen, 2023), yielding 408.19 Gb of clean data. The filtered Hi-C reads were aligned to the reference genome using Juicer v1.5.6 (Durand et al., 2016), and uniquely mapped valid contacts were extracted for downstream scaffolding and chromosome anchoring.

**Transcriptome library preparation and sequencing**

**For the different developmental stage experiment.** To assist in genome annotation, multiple tissues across key developmental stages were collected (Supplementary Figure 1). Root, stem, and leaf tissues were sampled from the same maternal plant at the full-blooming stage, with fully expanded leaves taken from the third node from the apex. Flower buds were collected at three developmental stages: initial, color-showing, and pre-anthesis. Inflorescences were harvested at both early- and full-blooming stages. Transcriptome sequencing was performed using both the BGISEQ platform (short reads) and the PacBio Sequel I platform (full-length Iso-Seq reads).

**For the abiotic stress treatments.** To analyze environmental adaptability, *O. longilobus* plants at the 6-leaf growth stage were exposed to two types of abiotic stresses. Under drought treatment (20% PEG), plants were deprived of water for 3 days and the third fully expanded leaf (from top to bottom) was sampled. For waterlogging stress, plants were imposed by keeping the water 3 cm above the soil surface for 12 h and root tissues were collected for transcriptome sequencing. Meanwhile, control samples from plants under normal conditions were collected in parallel for each stress treatment (drought and waterlogging). Each treatment included three biological replicates, with three individual plants per replicate.

**For the identification of candidate OMT genes.** Plants of *O. longilobus*, *Chrysanthemum morifolium* cv. 'Jinling Zhipao', and their $F_1$ hybrid $F_1$(1-3) were cultivated in a controlled greenhouse at Nanjing Agricultural University (Nanjing, Jiangsu, China; 118.841170°E, 32.034760°N). Inflorescences at the early-flowering stage were collected for subsequent RNA-seq analysis. All samples were immediately frozen in liquid nitrogen and stored at -80°C, with three biological replicates (Supplementary Table 1).

Total RNA was extracted using a modified CTAB phenol/chloroform method developed by BGI Genomics (Shenzhen, China). RNA integrity and purity were assessed using an Agilent 2100 Bioanalyzer (Agilent Technologies, USA) and a NanoDrop spectrophotometer (Thermo Fisher Scientific, USA), and only high-quality RNA samples were used for library preparation. mRNA libraries were constructed using the MGIEasy RNA Library Prep Kit (MGI Tech, Shenzhen, China) and sequenced on the DNBSEQ-2000 platform to generate 150 bp paired-end reads. For full-length transcriptome sequencing, pooled total RNA from all sampled tissues was converted into double-stranded cDNA with the UMI-based PCR cDNA Synthesis Kit (MGI Tech, Shenzhen, China) and used to construct a SMRTbell library for sequencing on the PacBio Sequel I platform (Pacific Biosciences, USA).

**Karyotyping and genome size estimation**

Before genome assembly, the genome size of *O. longilobus* was estimated using flow cytometry and *K*-mer analysis based on clean short-read sequencing data.

**Ploidy determination.** Flow cytometric analysis was performed as previously described (Hare and Johnston, 2011). Fresh leaf tissue (~0.2 g) was chopped in 500 μL nuclei extraction buffer for 1 min to release nuclei, and the homogenate was filtered through a 50 μm Biosharp filters. The filtrate was stained with 2 mL RNase-containing buffer in the dark for 15 min. Nuclei suspensions were analyzed on a CyFlow Space flow cytometer (Sysmex Partec, Germany), and fluorescence data were processed using FloMax v2.3. Genome size estimations were independently conducted using maize (*Zea mays*) as the internal reference standard. The mean fluorescence intensities of $G_0/G_1$ nuclei for *O. longilobus* ($I_s$) and for the maize standard (Ick) were measured, and the genome size (F) of *O. longilobus* was estimated as $F = I_s$ / Ick × n (n, genome size of maize), yielding ~3.36 Gb (Supplementary Figure 2A-F).

***K*-mer-based genome survey.** *K*-mer analysis was performed using Jellyfish (v2.2.6; $K$ = 17) (Marçais and Kingsford, 2011) with 389.62 Gb of clean data (~129.9× coverage) to generate the *K*-mer frequency distribution. Based on this distribution, GCE v1.0.2 (ftp://ftp.genomics.org.cn/pub/gce) (Vurture et al., 2017)was used to estimate genome size, heterozygosity, and repeat content (Supplementary Figure 3). Smudgeplot v0.2.5 (Ranallo-Benavidez et al., 2020) was used to infer genome ploidy and heterozygosity based on the coverage distribution of heterozygous *K*-mer pairs.

**Karyotype analysis and FISH analysis.** Metaphase chromosomes of *O. longilobus* were prepared from root tip meristems using an enzymatic digestion method. The preparations were used for karyotype observation and chromosome counting. Fluorescence *in situ* hybridization (FISH) was performed to detect 5S rDNA and telomeric sites (He et al., 2022). The 5S rDNA probes were designed from published sequences (JH5SrDNA-1 and JH5SrDNA-2). The telomeric probe was synthesized using the *Arabidopsis thaliana* repeat $(TTTAGGG)_n$ and labeled with fluorescein-12-dUTP (Thermo Fisher Scientific, USA). Fluorescent signals were counterstained with DAPI and observed under an Olympus BX60 fluorescence microscope (Olympus, Japan) (Supplementary Figure 2G-I).

### Supplementary Note 2. Genome assembly of *O. longilobus*

#### Contamination screening and organelle read filtering

After self-correction and trimming of low-quality regions, all sequencing data were screened for microbial contamination, and no substantial signals were detected. To prevent organellar sequences from interfering with the nuclear genome assembly, HiFi reads were aligned to the reference chloroplast and mitochondrial genomes using Minimap2 v2.24 (Li, 2021). Reads assigned to organellar origin were removed, and only the remaining nuclear HiFi reads were retained for downstream assembly.

#### Organelle genome assembly

To obtain the chloroplast and mitochondrial genomes of *O. longilobus*, HiFi long reads were first aligned to the species' reference plastid genomes, and all reads mapping to chloroplast or mitochondrial sequences were extracted. These organelle-derived reads were then assembled de novo using Hifiasm v0.19.5 (Cheng et al., 2021), generating draft chloroplast and mitochondrial genome sequences. Functional annotation of the assembled organelle genomes was performed using GeSeq (https://chlorobox.mpimp-golm.mpg.de/geseq.html). During annotation, ARAGORN v1.2.38 was run with a minimum gene length threshold of 3000 bp, and tRNAscan-SE v2.0.7 was executed with a cut-off score of 15 for reporting tRNAs. BLAST searches were conducted using identity thresholds of 25% for protein queries and 85% for rRNA, tRNA, and DNA sequences. The resulting annotations were further inspected and manually curated based on structural visualization in Bandage v0.9.0 to ensure the accuracy of organelle genome structure (Supplementary Figure 10A).

#### Assembly quality evaluation

The quality of the *O. longilobus* genome was assessed in four dimensions: continuity, accuracy, completeness and haplotype phasing.

Genome continuity was evaluated by calculating the contig N50 length and the LTR Assembly Index (LAI). Long terminal repeats (LTRs) were detected by LTRharvest (GenomeTools v1.6.1) (Ellinghaus et al., 2008). LAI was calculated by LTR_retriever v2.9.8 (Ou et al., 2018) with default parameters (Supplementary Table 14).

The PacBio long reads and RNA-Seq reads were mapped to the genome assembly with minimap2 v2.30 and HISAT2 v2.1.0 (Kim et al., 2015), respectively. PacBio reads were mapped with minimap2 under the settings "-ax map-hifi -secondary=no". SAMtools v1.15.1 (Danecek et al., 2021) was then used to calculate the genome coverage and mapping rate. The detailed mapping information of the samples could be found in Supplementary Table 6. Assembly quality was evaluated for structure accuracy by assessing the completeness of the Embryophyta_odb10 dataset with BUSCO v5.2.2 (Supplementary Table 8) (Waterhouse et al., 2018).

Haplotype phasing accuracy was assessed using Merqury v1.3 (Rhie et al., 2020) based on haplotype-specific *K*-mers (hap-*K*-mers). PacBio HiFi reads were used to generate hap-*k*-mer databases for Hap1 and Hap2, and hap-mer together with switch-error rates was calculated to quantify phasing quality (Supplementary Figure 6C; Supplementary Table 7). Finally, we manually checked and curated order and orientation errors based on Hi-C contact maps using Juicebox to achieve a chromosome-level haplotype-resolved assembly of *O. longilobus* genome (Supplementary Figure 6C).

**Telomere and centromere identification**

Telomeric regions were identified using the QuarTeT v1.2.0 (https://github.com/aaranyue/quarTeT) (Lin et al., 2023) teloexplorer module with the plant-specific model (-c plant -m 10), which detects telomere sequence $(AAACCCT)_n$ at chromosome termini. Although the majority of chromosomes exhibited well-assembled telomeric ends, there were a few cases where this sequence was missing. Centromeric regions were predicted with the QuarTeT centrominer module using the assembled genome and TE annotation under default settings, based on local enrichment of tandem repeats and transposable elements (Supplementary Figure 7C, D; Supplementary Tables 10 and 11).

**SV validation**

Hi-C reads were mapped to the haplotype assemblies using Juicer, and the contact matrices were subsequently inspected and visualized with Juicebox. In order to identify the shared inversions among haplotypes, we first determined the chromosomal correspondences between haplotypes using JCVI v1.5.7 (Goll et al., 2010), and subsequently selected the candidate regions that showed inversion signals in the Hi-C interaction maps. To align the HiFi reads to the two haplotypes of *O. longilobus*, minimap2 was used with the parameter set "-ax map-hifi", and the resulting alignments were visualized using the Integrative Genomics Viewer (IGV, v2.19.5) (Robinson et al., 2011; Thorvaldsdóttir et al., 2013). In addition, minimap2 was also employed with the parameters "-ax asm5, --eqx" to align the sequences across the inversion regions, and SyRI v1.4 (https://github.com/schneebergerlab/syri) (Goel et al., 2019) was subsequently used to further identify inversion and translocation events (Supplementary Figure 6B).

### Supplementary Note 3. Genome annotation of *O. longilobus*

**Genome repeat annotation**

For homology-based TE identification, the genome was searched against the Repbase TE database (https://www.girinst.org/repbase) using RepeatProteinMask v4.0.7 (http://www.repeatmasker.org/; parameters: -no_low -no_is -norna -engine ncbi -parallel 1) and RepeatMasker v4.0.7 (http://www.repeatmasker.org/; parameters: -engine ncbi -no LowSimple -pvalue 0.0001) (Tarailo-Graovac and Chen, 2009). To identify long terminal repeat retrotransposons (LTR-RTs), the LTR_retriever v2.9.8 (Ou and Jiang, 2017) pipeline was employed to integrate candidate LTR-RTs detected by LTRharvest (GenomeTools v1.6.1; parameters: -similar 90 -vic 10 -seed 20 -minlenltr 100 -maxlenltr 7000 -mintsd 4 -maxtsd 6 -motif TGCA -motifmis 1). A species-specific TE library was constructed by incorporating LTR-RTs identified above and additional transposable elements predicted using LTR_harvest and RepeatModeler v2.0.1 (http://www.repeatmasker.org/RepeatModeler.html; parameters: -database mydb -pa 6) (Flynn et al., 2020). This library was then combined with the Repbase database, and the merged dataset was utilized by RepeatMasker v4.0.7 to identify and classify transposable elements. The final set of repetitive sequences in the *O. longilobus* genome was obtained by integrating the ab initio-predicted TEs and those identified by homology through RepeatMasker v4.0.7. Additionally, the package Tandem Repeats Finder v4.09.1 (Benson, 1999) was used to identify tandem repeats in the genome with the parameters '2 7 7 80 10 50 2000 -d -h' (Supplementary Table 12). The locations of centromere and telomere were inferred from the outputs generated by Tandem Repeats Finder.

**Structural annotation of genes**

The genome of the *O. longilobus* was masked for repetitive elements by generating a *de novo* repeat library with RepeatModeler and used to mask the genome with RepeatMasker. The protein-coding genes were predicted from the repeat-masked genome using GeMoMa v1.9 (http://www.jstacs.de/index.php/GeMoMa) (Keilwagen et al., 2019), which integrates evidence from protein homology, transcripts, and ab initio predictions. The homology-based evidence was derived by aligning protein sequences from six plant species (*A. annua*, *C. lavandulifolium*, *C. morifolium*, *C. nankingense*, *C. seticuspe*, and *H. annuus*) to each *O. longilobus* genome assembly. Homology-based gene prediction was also carried out using the software GeMoMa v1.9. In parallel, we performed an evidence-based gene model prediction by mapping *O. longilobus* transcriptome datasets onto the assembled genome sequences, using HISAT2 v2.1.0 and GMAP v2020-06-02 (Wu and Watanabe, 2005) pipeline. For gene model prediction, GeMoMa (parameters: AnnotationFinalizer.r=NO AnnotationFinalizer.u=YES GeMoMa.g=25 GeMoMa.ge=5 GeMoMa.sil=false GeMoMa.m=15000) was used, integrating protein-coding gene alignments, intron position conservation, and RNA-seq evidence to refine intron boundaries (Supplementary Table 16).

**ISO-Seq full-length transcriptome data processing**

A SMRTbell library with an average insert size of 3.6 kb was constructed and sequenced, generating 56.69 Gb of PacBio subreads. The long-read data were processed with the SMRT analysis suite, resulting in 1,389,656 full-length non-chimeric (FLNC) reads. Isoforms were clustered using the Iterative Clustering and Error Correction (ICE) algorithm, polished with Quiver, and deduplicated with CD-HIT-EST (v4.8.1, parameters: -c 0.98 -aL 0.90 -aS 0.98 -AL 100 -AS 30 -T 6 -G 0) (Li and Godzik, 2006) to generate high-quality full-length transcripts for genome annotation. Then, the Gffread (Pertea and Pertea, 2020) was used to extract mRNA sequences from the genome assembly. TransDecoder v3.0.1 (https://github.com/TransDecoder/TransDecoder) was used to predict coding sequences and peptide sequences from mRNA sequences.

**RNA sequencing data analysis**

Short-read RNA-seq data were quality-filtered using SOAPnuke (v1.5.2; parameters: -l 15 -q 0.2 -n 0.001) (Chen et al., 2018) to remove adapters, low-quality reads, and reads containing excessive ambiguous bases (N). The sets of RNA sequencing data were each approximately ~6 Gb in size. For genome-guided transcript assembly, RNA-seq reads were mapped to the repeats-masked *O. longilobus* genome using HISAT2 v2.1.0 with the parameters "--sensitive --no-discordant --no-mixed -I 1 -X 1000". Transcript reconstruction was performed using StringTie v2.2.1 (Pertea et al., 2015) with the parameters "-f 0.3 -j 3 -c 5 -g 100 -s 10000", followed by merging assemblies across samples and removing redundant transcripts.

**Identification of non-coding RNA**

The annotation of tRNAs was performed by tRNAscan-SE v1.3.1 (Lowe and Eddy, 1997). rRNA sequences from *A. thaliana* and *O. sativa* were used as reference sequences to annotate rRNA. BLAST v2.2.26 (Altschul et al., 1990) was used for the mapping of rRNAs with the parameters "-p blastn -e 1e−5 -v 10000 -b 10000". MicroRNA (miRNA) and snRNA annotations were identified using the CMsearch program of the INFERNAL software package v1.1.2 (Nawrocki and Eddy, 2013) by searching against the Rfam database (e.g., Rfam_12.0/Rfam/RF00906.cm and Rfam_12.0/Rfam/RF00097.cm) (Supplementary Table 15).

**Functional annotation of protein-coding genes**

The function of the proteins encoded by all predicted genes was annotated using InterProScan v5.59-91.0 (Zdobnov and Apweiler, 2001) by searching publicly available database. For functional annotation, all the predicted genes were aligned to six public databases, including SwissProt (http://www.uniprot.org/), GO (Gene Ontology, http://geneontology.org/), TrEMBL (http://www.uniprot.org/), KEGG (Kyoto Encyclopedia of Genes and Genomes, http://www.genome.jp/kegg/), InterPro (https://www.ebi.ac.uk/interpro/), NR (Non-Redundant Protein Sequence Database, https://ftp.ncbi.nlm.nih.gov/blast/db/FASTA/nr.gz), KOG (Eukaryotic

Orthologous Groups, https://ftp://ftp.ncbi.nih.gov/pub/COG/KOG) using DIAMOND (v2.1.9) (Buchfink et al., 2015) with an E-value of 1e-5 as the cutoff (Supplementary Table 20). Gene Ontology identifiers for each gene were obtained using Blast2GO v4.1 (Conesa et al., 2005) according to the blast results combined with InterPro GO entries. Existing Gene Ontology terms were then mapped to enzyme codes by Blast2GO and predicted proteins were submitted to KAAS to get KO numbers for KEGG pathway annotation. The program PlantTFDB 5.0 (https://planttfdb.gao-lab.org/) was used to detect transcription factors in the *O. longilobus* genome (Supplementary Table 21).

**Estimation of the LTRs insertion time**

LTRharvest embedded in GenomeTools v1.6.1 was employed to detect structurally complete LTR retrotransposons using the parameters "-similar 90 -vic 10 -seed 20 -minlenltr 100 -maxlenltr 7000 -mintsd 4 -maxtsd 6 -motif TGCA -motifmis 1", enabling sensitive identification of young LTR elements (Supplementary Figure 12A; Supplementary Table 13). For each intact LTR-RT, insertion time was inferred from the sequence divergence between its 5' and 3' LTRs. High-quality intact LTR-RTs were further curated, and their insertion ages estimated using LTR_retriever v2.9.8 with the substitution rate parameter set to -u 7e-9 (Wen et al., 2022; Cao et al., 2025). The paired 5' and 3' LTR sequences were aligned to calculate nucleotide divergence ($\lambda$), followed by estimation of the DNA substitution rate ($K$) using the Kimura two-parameter model $K = -0.75 \ln (1 - 4\lambda/3)$. Finally, insertion time ($T$) was calculated as $T = K / (2r)$, where $r$ corresponds to the general substitution rate of $7 \times 10^{-9}$ substitutions per site per year reported for the Asteraceae family (Supplementary Figure 12B-D).

### Supplementary Note 4. Genome evolution of *O. longilobus*

**Phylogenetic analysis**

The protein sequences of *Oryza sativa* (rice) and *Zea mays* (maize); basal angiosperms represented by *Amborella trichopoda*; core eudicots including *Chenopodium quinoa* (Caryophyllales), *Aquilegia coerulea* (Ranunculales), *Vitis vinifera* (Vitales), *Solanum lycopersicum* (Solanales), *Daucus carota* (Apiales), and *Coffea canephora* (Rubiaceae); and Asteraceae species from the order Asterales: *Helianthus annuus*, *Lactuca sativa*, *Cynara cardunculus*, *Artemisia annua*, *Artemisia argyi*, *Chrysanthemum lavandulifolium*, *Cichorium intybus*, *Chrysanthemum nankingense*, *Chrysanthemum seticuspe*, *Chrysanthemum morifolium*, *Glebionis coronaria*, *Erigeron canadensis*, *Chrysanthemum makinoi*, *Cichorium endivia*, *Arctium lappa*, *Stevia rebaudiana*, *Smallanthus sonchifolius*, *Taraxacum mongolicum*, *Taraxacum kok-saghyz*, *Carthamus tinctorius*, *Mikania micrantha* and *Erigeron breviscapus* were downloaded from Ensembl Plants (http://plants.ensembl.org/index.html), and NCBI (https://www.ncbi.nlm.nih.gov/)

(Supplementary Table 22). All the genes of these species were filtered as follows: (a) the longest transcript was selected for genes with multiple splice variants, (b) the genes encoding proteins with fewer than 30 amino acids were filtered out, and (c) sequences containing frameshifts or coding regions whose lengths were not multiples of three were removed. To study the evolution of the *O. longilobus*, the first step is to identify orthologous genes from selected closely related species genomes and to further get the *O. longilobus* specific gene families. An all-versus-all comparison of protein sequences was performed using DIAMOND v2.1.9. Then, OrthoFinder v2.5.4 (Emms and Kelly, 2015) was used to cluster the similarities gene pairs into groups. Single-copy genes for each plant species for a given orthogroup were aligned using MAFFT v7.505 (Katoh and Standley, 2013), and alignments were concatenated to create a super-alignment matrix. The concatenated alignment was subsequently used to construct a maximum likelihood phylogenetic tree using FastTree v2.1 (Price et al., 2010), IQ-TREE v2.2.6 (Minh et al., 2020) and RAxML v8.2.12 (Stamatakis, 2014). For the species-level phylogeny across angiosperms, *Amborella trichopoda* was used as the outgroup, whereas *Scaevola taccada* was used as an outgroup for the Asteraceae-focused phylogeny.

The Bayesian Relaxed Molecular Clock (BRMC) method was utilized to estimate the divergence time of species by MCMCtree, a part of PAML package v4.5 (Yang, 1997). It was executed using the following parameters: "burn-in = 5,000,000, sample frequency = 5,000, sample number = 1,000,000". For divergence time estimation, the model was calibrated using various fossil records selected from previous publications and TimeTree (http://www.timetree.org/). Specifically, the following divergence intervals were applied: *A. trichopoda* and *Z. mays*: 179.9-205.0 Mya, *A. coerulea* and *O. sativa*: 142.1-163.5 Mya, *V. vinifera* and *C. canephora*: 111.4-123.9 Mya, *S. lycopersicum* and *C. canephora*: 72.5-97.6 Mya, *O. sativa* and *Z. mays*: 41.4-51.9 Mya, *C. cardunculus* and *H. annuus*: 38-50 Mya, *C. cardunculus* and *L. sativa*: 29 Mya, *C. seticuspe* and *C. nankingense*: 2.7-3.0 Mya, *S. taccada* and *C. cardunculus*: 80 Mya, *L. sativa* and *T. kok-saghyz*: 9.3-22.5 Mya, *T. mongolicum* and *C. cardunculus*: 36.6-45.1 Mya, *H. annuus* and *M. micrantha*: 16.0-27.0 Mya, *M. micrantha* and *S. rebaudiana*: 3.1-30.3 Mya, *E. breviscapus* and *C. cardunculus*: 7.4-8.6 Mya, *E. breviscapus* and *H. annuus*: 28.5-40.6 Mya, *G. coronaria* and *A. annua*: 6.8-34.6 Mya, *G. coronaria* and *E. breviscapus*: 23.59 Mya, *A. annua* and *C. morifolium*: 6.0-7.0 Mya. The phylogenetic tree with calibrated divergence time was visualized using FigTree (http://tree.bio.ed.ac.uk/software/Figuretree/).

**Expansion and contraction of gene families**

The expansion and contraction of gene families were inferred by comparing family size changes between ancestral and extant species using the CAFÉ program v4.2.1 (De Bie et al., 2006). Modeling of gene family size was performed using CAFE and the gene birth and death rate was estimated with orthologous groups that were conserved in all selected species. CAFE parameters were set to *p*-value threshold = 0.05, and auto searching for the λ value. The algorithm in CAFE takes a matrix of gene family sizes in extant species as input and uses a probabilistic graphical model (PGM) to ascertain the rate and direction of changes in gene family size across a given phylogenetic tree.

**Genome synteny and whole-genome duplication**

Cross-species synteny between *O. longilobus* and related species (*C. morifolium*, *C. lavandulifolium*, *C. nankingense*, and *C. seticuspe*) was visualized using the Dual Synteny Plot module of MCScanX implemented in TBtools v2.154 (Chen et al., 2020) (Supplementary Figure 9). We identified and localized whole-genome duplication (WGD) events by combining intra- and inter-species synteny analysis and *Ks* distribution. Using the JCVI synteny pipeline, LAST output was filtered with a c-score threshold of 0.7 to retain significantly matching sequences within the *O. longilobus* genome. Dotplots for genome pairwise synteny visualization was generated using the command 'python -m jcvi.graphics.dotplot' (Supplementary Figure 11C, D).

WGDI (Sun et al., 2022) with the '-icl' parameter was used to identify the intergenomic synteny blocks between *O. longilobus* and others, as well as intragenomic synteny blocks within each species. The *Ks* between collinear genes was estimated by Nei_Gojobori approach in PAML with the parameter '-ks' of WGDI, and the '-bk' parameter was applied to generate a dot plot of collinear genes and *Ks* values, visualizing intra- and inter-species synteny. Additionally, the *Ks* peaks were fitted using the '-pf' parameter, and the density distribution curve of *Ks* was displayed using the '-kf' parameter (Supplementary Figure 11A, B). The location of the WGD event was identified based on the comparison of the *Ks* peaks between paralogs within species and orthologs between species. Different modes of gene duplication as whole-genome duplicates (WGD), tandem duplicates (TD), proximal duplicates (PD), transposed duplicates (TRD), or dispersed duplicates (DSD) were identified using DupGen_finder (Qiao et al., 2019) with default parameters (https://github.com/DXXDR/DupGen_finder; Supplementary Figure 11E, F).

**Positive selection analysis**

For positive selection analyses, *O. longilobus* was set as the foreground lineage, with *C. morifolium*, *C. lavandulifolium*, *C. nankingense*, and *C. seticuspe* defined as background lineages. Single-copy orthologs were identified with OrthoFinder v3.0.1b1. Protein multiple sequence alignments (MSAs) were generated using MAFFT v7.526 and trimmed with Gblocks v0.91b (Castresana, 2000). Codon MSAs were generated using ParaAT v2.0 (Zhang et al., 2012) via pal2nal.pl (http://www.bork.embl.de/pal2nal/), followed by removal of gap-rich positions. Gene trees were inferred with RAxML v8.2.10 under the best-fit JTT+I+F model, and positive selection was assessed using the branch model implemented in codeml (PAML v4.10.7) (Supplementary Table 29).

**Phylogeny analysis of family Asteraceae based on chloroplast genome sequence**

We downloaded 18 chloroplast genomes from Asteraceae species from the NCBI database (https://www.ncbi.nlm.nih.gov/), including *Cynara cardunculus* (KM035764), *Arctium lappa* (NC_042724), *Carthamus tinctorius* (KX822074), *Lactuca sativa* (NC_007578), *Taraxacum mongolicum* (NC_031396), *Cichorium* *intybus* (MK569377), *Cichorium endivia* (NC_082974), *Helianthus annuus* (NC_007977), *Smallanthus sonchifolius* (MT872397), *Stevia rebaudiana* (OM927972), *Mikania micrantha* (NC_031833), *Erigeron breviscapus* (MK279916), *Erigeron canadensis* (MT806101), *Glebionis coronaria* (MW874476), *Artemisia annua* (KY085890), *Artemisia argyi* (NC_030785), *Chrysanthemum lavandulifolium* (NC_057202.1), and *Chrysanthemum morifolium* (MT919681). Here, the *O. longilobus* chloroplast assembly was integrated with 18 previously published chloroplast genomes to yield a comprehensive phylogenomic analysis of *Opisthopappus* and other related Asteraceae genera (Supplementary Figure 10B). Among them, the chloroplast genome of *Cynara cardunculus* was used as the outgroup. The quadripartite structure of the chloroplast genome (LSC, SSC, and IRs) was oriented using cpstools v2.6 (Huang et al., 2024) with the parameters "-m LSC, -m RP, -m SSC". The consistency of region orientation was validated using MUMmer v3.23 (Kurtz et al., 2004) with the following parameters: "nucmer -mum, delta-filter -m, show-coords -T -r -l", and visualized with mummerplot. All chloroplast genome sequences were aligned using MAFFT v7.505 with the *-auto* option. Conserved blocks from the alignment were extracted using GBLOCKS v0.91b. The best-fit evolutionary model was selected with ModelTest v0.1.7 (Posada and Crandall, 1998). Phylogenetic reconstruction was performed in RAxML-NG v1.2.2 under the maximum likelihood framework using the TVM+I+G4 substitution model with 1,000 bootstrap replicates. The resulting phylogenetic tree was visualized and annotated using iTOL (https://itol.embl.de/).

**GO and KEGG enrichment analysis**

GO and KEGG enrichment analysis of gene sets were performed in R software (https://www.r-project.org) against *O. longilobus* genome as reference. *P* values of the enrichment analysis were calculated with a hypergeometric test using the phyper function, followed by multiple-testing correction using the Benjamini–Hochberg false-discovery rate (FDR) method via *P*.adjust. In addition, *q*-values were calculated using the qvalue package in R. Significant enrichment was defined as *P* value $\leq 0.05$.

**Homology-based annotation**

As for the model plant, gene numbers starting with ATXG are widely used in *Arabidopsis thaliana*. In order to minimize the difference from previous gene annotations, we use a reciprocal BLAST v2.12.0 to map the Araport gene annotation onto each haplotype *O. longilobus* genome. BLAST databases for both *O. longilobus* (query) and *A. thaliana* (reference; TAIR10) were constructed using makeblastdb. Reference sequences were first aligned to the query database with a stringent E-value cutoff ($1 \times 10^{-5}$) to identify candidate homologs. These candidates were then aligned back to the *A. thaliana* database to define reciprocal best hits (RBHs) and assign the corresponding *A. thaliana* gene IDs. Query genes without RBH support were re-searched using a relaxed cutoff ($1 \times 10^{-2}$), and those still lacking significant hits were designated as unannotated.

### Supplementary Note 5. Whole-genome resequencing of *Opisthopappus* populations

#### DNA extraction, library construction, and sequence quality control

The genomic DNA was extracted from leaves of 115 *Opisthopappus* individuals using a Qiagen DNeasy plant kit, and the samples were sent to BGI Genomics Co., Ltd (Shenzhen, China) for sequencing library preparation according to the manufacturer's specifications. In brief, the DNA samples were fragmented by Covaris to a size of ~300 bp. Qubit dsDNA HS Assay kit, 500 assays (Thermo Fisher Scientific, USA) was used to measure the quantity of purified DNA samples. Then the DNA fragments were end-polished, A-tailed, and ligated with the full-length adapters for BGI sequencing with further PCR amplification. PCR products were then purified using the Agencourt AMPure XP-Medium system. The libraries were analyzed for size distribution using an Agilent 2100 Bioanalyzer, and were quantified using real-time PCR. Subsequently, we used the DNBSEQ-T7 platform to generate 150 bp paired-end reads.

Adapter and low-quality reads were removed with SOAPnuke software using the following criteria: (1) reads with 0.1% unidentified nucleotides (N); (2) reads with > 50% bases aligned to the adapter; (3) reads with >50% bases having Phred quality < 10; and (4) putative PCR duplicates generated via PCR amplification during the library construction process. Consequently, we obtained 12.34 Tb (~107.31 Gb per sample) high-quality genomic data (Supplementary Table 32).

#### Linkage disequilibrium analysis and ROH detection

To estimate and compare the LD pattern among different groups, the squared correlation coefficient ($r^2$) between pairwise SNPs without LD filtering was calculated using PopLDdecay v3.42 software (Zhang et al., 2019). The program parameters were set as "-MaxDist 300 -MAF 0.05 -Miss 0.25". The average $r^2$ value was computed for pairwise SNPs in a 300 kb window and averaged across the whole genome. LD decay curves were generated for each group using R package ggplot2 and LD decay distance was determined by the maximum $r^2$ dropped by half. Local LD heatmap was constructed by R package LDBlockShow (Dong et al., 2021).

The ROHs were detected by PLINK v1.9 with the following parameters "--homozyg --homozyg-density 50 --homozyg-gap 100 --homozyg-kb 100 --homozyg-snp 50 --homozyg-window-het 5 --homozyg-window-snp 50 --homozyg-window-threshold 0.05". Then, ROHs were classified into different categories based on their length, and a longer ROH suggests the presence of recent inbreeding.

### Supplementary Note 6. Genotype-environment association analysis

#### Collection of bioclimatic variables

In genome-environment associations analysis, the 19 bioclimatic variables layer (Bio1-Bio19) were download from WorldClim v2.1 database (https://www.worldclim.org/data/worldclim21.html) at 30s spatial resolution (~1 km) using the longitude and latitude information for 115 resequenced samples. The tiff files were transferred into bioclimatic data in R package raster v3.6.32. The Pearson's correlation coefficients were calculated among the 19 bioclimatic variables and visualized in R packages corrplot. We selected the top climatic factors based on weighted $R^2$ importance using a machine learning gradient forest (GF) model in the R package gradientForest v.0.1.37 (Ellis et al., 2012) by building a relationship between climatic variables and genetic variants with 500 regression trees.

Here, we used a subset of five hundred thousand SNPs that randomly selected from the high-quality LD-prunned SNPs with the aim of reducing computing resources. To avoid multicollinearity, seven important environmental variables (Supplementary Figure 30) with absolute correlation coefficient < 0.6 were retained for RDA analysis.

In genomic offset analysis, future climate projections were obtained from the WorldClim CMIP6 (Coupled Model Intercomparison Project Phase 6) (https://www.worldclim.org/data/cmip6/cmip6_clim30s.html) with a spatial resolution of 30s (~1 km). The four general circulation models (GCMs) were: ACCESS-CM2 (Australian Community Climate and Earth System Simulator Coupled Model version 2), BCC-CSM2-MR (Beijing Climate Center-Climate System Model version 2-Medium Resolution), CMCC-ESM2 (Centro Euro-Mediterraneo per I Cambiamenti Climatici-Earth System Model2), and GISS-E2-1-G (Goddard Institute for Space Studies Model E version 2.1 coupled with the GISS Ocean). SSP126 and SSP585 are two shared socioeconomic pathways (SSPs) that represent the low and high future emission scenarios, respectively. Specifically, the SSPs are combinations of shared socioeconomic pathways and representative concentration pathways (RCPs). SSP126 is the abbreviation for SSP1-RCP2.6 scenario. SSP1 assumes a gradual shift toward a more sustainable world, with emphasis on human well-being and reduced inequality. RCP2.6 represents one mitigation scenario leading to a very low forcing level. Similarly, SSP585 is the shortest form for SSP5-RCP8.5 scenario. SSP5 assumes rapid economic growth driven by fossil fuels, with high energy demand and limited efforts to mitigate greenhouse gas emissions. RCP8.5 indicates a very highbaseline emission scenario (Hou et al., 2024).

#### IBD and IBE analyses

We employed the isolation-by-distance (IBD) and isolation-by-environment (IBE) analyses to quantity the relative importance of environmental and geographical factors in shaping the spatial distribution of genetic scenarios as well as to investigate the role of adaptive variants. The 4,620 adaptive variants and the same number of neutral variants that randomly selected from LD-pruned dataset were separately used to calculated genetic distance ($F_{ST}/(1-F_{ST})$). The geographical distance matrix was computed from an initial coordinate matrix using the distm function in R package geosphere v1.5.20. The environmental distance matrix was obtained with the vegdist function in R package vegan v2.7.1 using the euclidean method. The mantel and partial mantel tests (excluding geography or environment influence) were implemented in R package vegan with significance determined using 999 random permutations. And the matrix correlations between geography and environment (autocorrelation), genetic distance and geography (IBD), and genetic distance and environmental (IBE) were separately analyzed.

### Supplementary Note 7. Mining of the linarin metabolism pathway and associated candidate genes

#### Chemicals

Apigenin, acacetin, tilianin, and linarin were sourced from Beijing Huayueyang Biotechnology (Beijing, China), while S-adenosyl-L-methionine (SAM) was obtained from Shanghai Yingxin Laboratory (Shanghai, China). The purity of all standards was confirmed to be ≥ 98%.

#### Metabolite profiles measured by LC/LC–MS

The capitula were collected from *Opisthopappus* species and their close relatives at full bloom (Supplementary Table 49). We also sampled capitula, leaves, roots, and stems from annual *O. longilobus*, *C. morifolium* cv. ‘Jinling Zipao’, and their intergeneric hybrid $F_1$(1-3). Briefly, 0.1 g of dried powder of each sample was used to extract bioactive components with methanol, followed by a 60 min sonication period utilizing an ultrasonic cleaner (Billerica, MA). Subsequently, the extracts were centrifugated at 4,000 rpm for 20 min, and the supernatants were collected. The extracts were filtered through a 0.22 μm organic membrane and stored at 4°C until Ultra Performance Liquid Chromatography (UPLC) analysis. UPLC analysis was performed on a Waters ACQUITY UPLC H-Class system (Waters, Milford, MA, USA). Samples were separated on an ACQUITY UPLC HSS T3 column (100 × 2.1 mm, 1.8 μm) with a detection wavelength of 326 nm and an injection volume of 2 μL.

For UPLC analysis of *Opisthopappus* species and their close relatives, the mobile phase consisted of solvent A ($V_{acetonitrile}$:$V_{water}$ = 2:98, containing 0.1% formic acid) and solvent B ($V_{acetonitrile}$:$V_{water}$ = 50:50, containing 0.1% formic acid). The gradient elution program was as follows: 0.0-1.5 min, 88% A; 1.5-3.0 min, 75% A; 3.0-8.0 min, 70% A; 8.0-12.0 min, 64% A; 12.0-15.0 min, 0% A; 15.0-20.0 min, 88%

A. The flow rate was 0.30 mL/min. The column temperature was 40°C.

For UPLC analysis of annual *O. longilobus*, 'Jinling Zipao', and their $F_1$ progeny, the mobile phases consisted of ultrapure water with 0.1% formic acid (A) and acetonitrile with 0.1% formic acid (B). Samples were separated under a gradient elution program as follows: 0.00 min, 85% A; 0.00-11.00 min, 65% A; 11.00-13.00 min, 40% A; 13.00-14.00 min, 5% A and held until 16.00 min; 16.00-18.00 min, 85% A. The flow rate was 0.80 mL/min, and the column temperature was set to 30°C (Supplementary Figure 41B, E, G; Supplementary Table 51).

For targeted flavonoid metabolite analysis, the third and fourth fully expanded leaves from approximately one month age rooted cuttings, along with capitula at full bloom of *O. longilobus*, 'Jinling Zipao', and their $F_1$ progeny, were collected. All tissues were vacuum freeze-dried, and 50 mg of dried powder was weighed into centrifuge tubes. Each sample was extracted with 1.2 mL of 70% methanol and centrifuged at 12,000 rpm for 3 min. The supernatants were filtered through a 0.22 μm organic membrane and stored at 4°C until UPLC-MS/MS analysis. UPLC-MS/MS analyses were performed using an ExionLC AD UPLC system coupled to an Applied Biosystems 4500 QTRAP mass spectrometer equipped with an electrospray ionization (ESI) source operating in multiple reaction monitoring (MRM) mode.

Samples were separated on an Agilent SB-C18 column (1.8 μm, 2.1 × 100 mm). The mobile phase consisted of ultrapure water with 0.1% formic acid (A) and acetonitrile with 0.1% formic acid (B). The analytes were eluted using a gradient for mobile phase B as follows: 5% at 0.00 min; a linear increase to 95% from 0.00-9.00 min; held at 95% from 9.00-10.00 min; decreased to 5% from 10.00-11.10 min; and maintained at 5% until 14.00 min for column re-equilibration. The flow rate was set to 0.35 mL/min, and the column temperature was maintained at 40°C. The injection volume was 4 μL. The ESI source operation parameters were as follows: ion source, electrospray; source temperature (TEM) 550°C; IonSpray voltage (IS) 5,500 V in positive mode and -4,500 V in negative mode. Curtain gas (CUR), ion source gas 1 (GS1), and ion source gas 2 (GS2) were set at 25, 50, and 60 pound per square inch (psi), respectively; the collision gas (CAD) was set to a medium level. Collision-induced dissociation (CID) was applied at a high setting to promote efficient fragment ion generation. Data acquisition was performed on a triple quadrupole (QQQ) mass spectrometer in multiple reaction monitoring (MRM) mode, using nitrogen as the collision gas at medium pressure. For each metabolite, precursor-product ion transitions were optimized by adjusting the declustering potential (DP) and collision energy (CE) to ensure specific and sensitive detection (Supplementary Figure 41C, D, F). Data analysis was performed using MultiQuant software.

The extraction procedure and untargeted metabolite profiling followed the published method (Tian et al., 2025), with the same plant materials used for targeted flavonoid metabolite profiling. Briefly, 50 mg of vacuum-dried tissue powder was weighed into a centrifuge tube and extracted with 1 mL of 80% methanol.

MS/MS fragment-ion spectra was performed on a UPLC–TripleTOF 5600+ system (AB SCIEX). Metabolites were separated on an ACQUITY UPLC HSS T3 column (2.1 × 100 mm, 1.8 μm) using a binary solvent system of 0.1% formic acid in water (A) and acetonitrile (B) at a flow rate of 0.3 mL/min. The gradient was: 5% B (0-2 min), 5-35% B (2-6 min), 35-95% B (6-12 min), 95% B (12-14 min), 95-5% B (14-14.1 min), and 5% B for re-equilibration (14.1-20 min). The column temperature was maintained at 40°C, and the injection volume was 2 μL. Mass spectrometry was conducted in ESI negative mode with an ion spray voltage of -5500 V and a source temperature of 550°C. Gas settings were 55 psi (GS1), 55 psi (GS2), and 35 psi (CUR). Declustering potential was 80 V, and collision energy was 35 eV. TOF-MS and MS/MS spectra were acquired over m/z 50-1600. Data acquisition was performed using SCIEX OS (Figure 5A).

Metabolite annotation followed standard high-resolution MS workflows. Peak tables were constructed and molecular features were matched against public and in-house spectral libraries (HMDB, MoNA, GNPS, PMhub) and further supported by in silico fragmentation using MSFinder v3.52 (Lai et al., 2018) and SIRIUS v5.6.3 (Böcker et al., 2008; Zhang et al., 2012). Network-based spectral grouping and feature consolidation were performed following the MetCirc pipeline (https://github.com/PlantDefenseMetabolism) (Naake and Gaquerel, 2017).

**Identification and phylogenetic analysis of OMT genes**

We used homologous alignment and hmmsearch to identify O-methyltransferase (OMT) proteins of *O. longilobus*. Firstly, we downloaded reported OMT protein sequences from the UniProt database (https://www.uniprot.org/) and used them as reference genes to compare with *O. longilobus* proteins using BLASTP with an E-value cutoff of 1e-10. For further identified OMTs, Pfam domains (http://pfam.xfam.org/, PF00891, PF08100, and PF01596) were used to search in the *O. longilobus* genome by HMMER v3.2.1 (Eddy, 2011). To further minimize false positives, an OMT HMM was built with hmmbuild from a ClustalW v2.1 (Thompson et al., 1994) alignment of curated OMT sequences. This model was then used to re-scan the *O. longilobus* genome with hmmsearch, and matches were retained at an E-value cutoff of 1e-5. OMT protein sequences were downloaded from the UniProt database to build a reference dataset, which was used to query the *O. longilobus* proteome by BLASTP (E-value < $1\times10^{-10}$) to identify putative OMT members; the candidates were then combined with those retrieved by HMMER, and conserved domains were annotated against Pfam v37.0 to remove false positives and retain bona fide OMT family genes.

After this, these genes were identified by building a phylogenetic tree. The Neighbor-joining tree was built using MEGA11 software (Tamura et al., 2021) after sequence alignment in MUSCLE. The resulting topology allowed classification of the *O. longilobus* OMT genes into COMT, and CCoAOMT subfamilies (Supplementary Figure 43A, B).

**Gene expression analysis**

Clean RNA-seq reads were aligned with HISAT2 v2.2.1, processed using SAMtools v1.15.1, and summarized at the gene level with featureCounts v2.0.1 (Liao et al., 2013). Differential expression analysis was performed using DESeq2 v1.42.1 (Love et al., 2014).

To characterize the expanded OMT gene family in *O. longilobus*, low-expression genes (RPKM ≤ 1) were filtered out based on transcriptome data. Genes showing expression patterns consistent with metabolite accumulation (lowest in 'Jinling Zipao', intermediate in $F_1$ hybrid, and highest in *O. longilobus*) and exhibiting correlation coefficients greater than 0.6 were retained as candidates for functional validation. To identify genes involved in the linarin-metabolism biosynthesis pathways, the key parameters of BLASTP v2.2.26 were E-value < 1e-5 and the identity > 90%. Gene expression patterns heatmaps across floral tissues were visualized using TBtools v2.154. Gene expression values were visualized after normalization using the min-max method.

### Supplementary Note 8. Functional validation of candidate OMT genes

#### Heterologous protein expression

Total RNA derived from the transcriptome samples used for identifying candidate OMT genes served as the template for downstream analyses. RNA extracted from transcriptome sequencing served as the template for cDNA synthesis, which was performed using the Evo M-MLV reverse transcriptase (Accurate Biology). *OlOMT250* and *OlOMT310*, along with their closest homologues, were amplified from the cDNA of *O. longilobus* and *C. morifolium* cv. 'Jinling Zipao' using specific primers (Supplementary Table 55). Both *OlOMT250* and *OlOMT310* were cloned into the pCold II vector, a His-tag expression vector. The *E. coli* BL21 (DE3) strain containing the appropriate pCold II_*OlOMT250* and pCold II_*OlOMT310* plasmids was employed for protein production.

Single colonies were inoculated into 5 mL of LB medium with ampicillin and cultured overnight at 37°C. The overnight cultures were then diluted 1:50 into 100 mL of LB/ampicillin in 250 mL flasks and grown at 37°C until reaching an $OD_{600}$ of 0.5-0.8. Following this, the cultures were cooled to 14°C, and protein expression was induced with the addition of isopropyl-β-D-thiogalactopyranoside (IPTG) to a final concentration of 0.5 mM. Induced cultures were incubated at 14°C for 16 h with shaking at 80 rpm. Cells were harvested via centrifugation (4000 rpm, 4°C, 20 min), washed twice with 1x phosphate-buffered saline (pH 7.5), and subsequently disrupted by sonication. The homogenates were centrifuged at 12000 rpm for 30 minutes at 4°C to separate soluble proteins from insoluble debris. His-tagged proteins were purified using a HisPur Ni-NTA spin column (Thermo Scientific). The purity and molecular mass of the recombinant proteins were assessed through 12% SDS-PAGE followed by staining with InstantBlue Coomassie (Abcam). Protein concentrations were determined via the Bradford assay using a Qubit™ 4.0 fluorometer (Thermo Fisher Scientific, USA). All experiments were conducted with three biological replicates.

**Creation of transgenic plants**

The primers (Supplementary Table 55) were designed by the Web MicroRNA Designer (http://wmd3.weigelworld.org/cgi-bin/webapp.cgi) to amplify amiR-*OlOMT250* and *OlOMT310* interference fragments. The PCR products were inserted into the pCVA vector, and transient expression was achieved by vacuum-infiltrating cuttings of *O. longilobus* with *Agrobacterium* suspensions.

To further validate the function of the candidate OMT genes identified above, genetic transformation experiments were performed using the *C. morifolium* cv. 'Nannong Fencui', which is characterized by a low accumulation of linarin. To obtain OE-*OlOMT250* lines or OE-*OlOMT310* lines of chrysanthemum, the full-length CDS of *OlOMT250* or *OlOMT310* was inserted into a pORE-R4-35AA vector. These vectors were subsequently transformed into *Agrobacterium tumefaciens* strain GV3101, and transgenic plants were obtained via leaf disc transformation.

**RT-qPCR analysis**

RNA extraction was performed using a commercial RNA isolation kit (Huayueyang Biotechnology, Beijing, China), and first-strand cDNA was synthesized with the Evo M-MLV RT Kit with gDNA Clean for qPCR II (Accurate Biotechnology, Changsha, China). Quantitative real-time PCR (RT-qPCR) was conducted on a Roche LightCycler480 II system using the SYBR Green Premix Pro Taq HS qPCR Kit (Accurate Biotechnology, Changsha, China). *CmEF1α* (GenBank: KF305681) and *OlEF1α* (OlChr9.1G031550.3) were used as internal reference genes. Relative transcript levels were calculated according to the $2^{\wedge-\Delta\Delta Ct}$ method (Livak and Schmittgen, 2001). All assays included three biological replicates, each with three technical replicates.

**Phenotypic characterization**

Overexpression lines (OE-*OlOMT310* and OE-*OlOMT250*) and wild-type *C. morifolium* cv. 'Nannong Fencui' were grown under uniform greenhouse conditions. Transient gene-silencing lines (pCVA-*OlOMT310* and pCVA-*OlOMT250*) were generated in *O. longilobus* using *Agrobacterium vacuum*-infiltration, with pCVA-empty as the control. Metabolic products generated in these plant materials were subsequently analyzed using *in vitro* enzyme assays, followed by targeted quantification on a Waters ACQUITY UPLC H-Class system coupled to a Triple Quad 6500+ mass spectrometer.

**UPLC-ESI-QQQ-MS analysis**

UPLC–MS/MS analysis was carried out using a Waters ACQUITY UPLC H-Class system paired with a Triple Quad 6500+ mass spectrometer equipped with an electrospray ionization (ESI) source operating in positive mode. The source temperature was maintained at 500°C, with both GS1 and GS2 set to 50 psi. MRM transitions were monitored for acacetin (Q1 = 285.2 m/z, Q3 = 242.1 m/z) and linarin (Q1 = 593.0 m/z, Q3 = 447.1/285.0 m/z). The dwell time was 150 ms, with a collision energy of 35 eV, a declustering potential of 10 V, and a collision cell exit potential of 11 V. Chromatographic separation was achieved on an ACQUITY UPLC HSS T3 column (100 × 2.1 mm, 1.8 μm) at 40°C. The mobile phases used were water with 0.1% formic acid (A) and acetonitrile (B), delivered at a flow rate of 0.4 mL/min with a 2 μL injection volume. The gradient program included 5% B for 0-1.0 min; a linear increase to 95% B from 1.0-.4 min; and re-equilibration to 5% B from 5.4-7.5 min.

**Structural analyses and binding energy calculation**

**Molecular docking analysis.** Molecular docking analysis was conducted using AutoDock Vina v1.2.5 (Trott and Olson, 2010). Protein structures of the target OMTs were predicted with AlphaFold2 v2.2.0 (Jumper et al., 2021) using the monomer_casp14 and monomer_ptm models. The predicted structures were processed in PyMOL v2.6.0a0 (https://pymol.org) to remove non-protein components and retain only the receptor backbone. Receptor preparation, including hydrogen addition, charge optimization, and conversion to AutoDock 4D4 atom types, was performed using AutoDockTools v1.5.7 (Morris et al., 2009). Ligand structures were obtained from PubChem (https://pubchem.ncbi.nlm.nih.gov/) in SDF format and converted to MOL2 after energy optimization in PyMOL. The putative binding pocket was predicted using POCASA v1.1 (https://g6altair.sci.hokudai.ac.jp/g6/service/pocasa), and a 60 × 60 × 60 Å³ grid box was applied for docking. Docking was performed with a maximum of 20 conformations. The protein-ligand complex was visualized in PyMOL, focusing on key interactions including hydrogen bonding, hydrophobic contacts, and π-π stacking. These results were used to infer the most plausible binding mode and to guide subsequent computational analyses.

**Molecular dynamics simulation.** Molecular dynamics simulations of the protein-ligand complexes were performed using GROMACS v2023.1. (Abraham et al., 2015) Ligand geometries were optimized in Gaussian 16 at the HF/6-31G* using an SCF convergence criterion of XQC and a maximum of 80 optimization cycles, followed by calculation of Merz-Kollman (MK) charges. Ligand parameters were generated with antechamber model in AMBERtools20 (Case et al., 2023) using BCC charge (Jakalian et al., 2000; Jakalian et al., 2002) correction and the GAFF force field (Wang et al., 2004), and subsequently converted into GROMACS-compatible formats. Protein models were parameterized using the OPLS-AA force field (Jorgensen et al., 1996), and missing residues as well as disulfide bonds were corrected with pdb2gmx. The resulting complexes were solvated in an SPC/E water box (Berendsen et al., 1987), and counterions ($Na^+$/$Cl^-$) were added to neutralize the system. Energy minimization was performed using steepest descent followed by conjugate gradient until the maximum force reached ≤ 100 kJ/mol/nm. System equilibration was carried out under NVT (300 K) and NPT (1 bar) ensembles. Production MD simulations were run for 300 ns with a 2 fs timestep, and trajectory frames were saved every 1 ps. Post-simulation analyses included root-mean-square deviation (RMSD), hydrogen bond counts, radius of gyration (Rg), and solvent-accessible surface area (SASA). All structural visualizations were performed using DuIvyTools v0.6.0 (https://github.com/CharlesHahn/DuIvyTools).
